# The diversity, evolution, and conservation of African plants

**DOI:** 10.64898/2026.09.20.752981

**Authors:** Wyckliffe Omondi Omollo, Xueqin Wang, Liguo Zhang, Xinru Zhang, Yuxuan Liu, Tao Xiong, Zijia Lu, Bing Liu, Shengwei Wang, Moses Kirega Gichua, Robert Wahiti Gituru, Alexandre Antonelli, Qing-Feng Wang, Zhi-Duan Chen, Miao Sun

## Abstract

Africa hosts remarkable biodiversity across diverse biomes, yet its evolutionary history at continental-scale remains poorly understood. We integrate a time-calibrated phylogeny covering 93% of Africa’s vascular plant genera with continental-scale distribution data to jointly evaluate phylogenetic diversity (PD), endemism, and diversification rates (DR). We reveal a pronounced decoupling between DR and accumulated evolutionary history across Africa. Mountain, arid, and semi-arid areas emerge as centers of neo-endemism and elevated diversification, whereas lowland and humid regions, particularly the Guineo-Congolian zone, are dominated by older lineages with low recent diversification. Several DR hotspots do not overlap with Global Biodiversity Hotspots, indicating a mismatch between rapid diversification and accumulated evolutionary history. High net diversification and speciation rates during the last 15 million years were likely driven by cooling, aridification, and mountain uplift that created new ecological gradients and isolation barriers. These abiotic changes, interacting with lineage-specific traits, shaped Africa’s modern flora and highlight conservation gaps, especially in arid regions.

## INTRODUCTION

Mainland Africa (excluding neighboring islands, hereafter “Africa”) is an ecologically and evolutionarily diverse continent, covering approximately 20% of the Earth’s land area. Spanning a broad range of climatic zones and biomes, from north to south: the hyper-arid Sahara, seasonally dry savannas, the tropical rainforests of the Congo Basin and the Mediterranean-type shrublands of the Cape Floristic Region (CFR) (Supplementary Fig. 1)^1–3^. It is home to seven Global Biodiversity Hotspots^4^ and harbors over 48,000 described native vascular plant species, distributed across 290 families and 4,023 genera^5,6^. Of these, 60% (28,800) of species, 40% (1,611) of genera, and 4% (12) of families are endemic to the continent. Tropical Africa is estimated to harbor between 30,000 and 35,000 plant species^5^, including the Congo Basin—the world’s second-largest tropical rainforest after the Amazon^7^. South Africa, with over 22,000 species, boasts the highest number of endemic species in Africa (about 13,265), while northern Africa has about 7,052 species, of which about 2,510 are endemic to the region^5,8^.

This immense diversity reflects Africa’s complex geological history, climate instability, and topographic heterogeneity, including the Gondwanan breakup (ca. 120–90 million years ago, Ma), which isolated regions and fostered distinct floras^9–12^. These processes promoted biome-specific diversification and left a legacy of shared plant lineages with regions once connected to Africa, such as Australia, Madagascar, South America, and Southeast Asia^13,14^. The Miocene (ca. 23–5.3 Ma) was a critical period in African plant evolution, marked by rainforest expansion followed by cooling, declining atmospheric CO_2_, and tectonic uplift that promoted aridification and the expansion of open habitats^10,15^.

Several hypotheses have been proposed to explain broad patterns of plant diversity, endemism, and diversification across African ecosystems. In tropical rain forests (TRF), the fragmentation-refugia hypothesis posits that historical climate oscillations drove cycles of forest expansion and contraction, potentially promoting vicariant speciation in TRF-restricted plant lineages^10,16–19^. These dynamics likely fragmented a once continuous pan-African rainforest into isolated refugia in West-Central and East Africa during arid periods^10,20,21^. Similar processes may also have affected savannas and montane ecosystems, contributing to diversification across multiple biomes^22^.

Major extinction pulses associated with the Eocene–Oligocene transition (ca. 33.9 Ma), Miocene aridification (23–13.8 Ma), and late Pliocene climate shifts likely erased substantial lineage diversity across African ecosystems^18,19,23–29^. Pollen fossils indicate that families such as Casuarinaceae, Chloranthaceae, Sarcolaenaceae, and Winteraceae were present in Africa at least during the Oligocene–Miocene, and possibly earlier, but are now extinct on the continent^30,31^, suggesting that extinction has played an important role in structuring modern floras. However, signals of these ancient extinction events remain underexplored outside rain forest systems e.g.,^18,19^ partly due to the continent’s scarce and temporally biased fossil record of angiosperm macrofossils^32^.

Clade-specific phylogenies have shown that humid western African lineages are often phylogenetically older, whereas dry-adapted eastern and southern African lineages typically reflect more recent diversification associated with Neogene aridification and tectonic uplift e.g.,^33–35^. Subsequent studies further linked biome shifts, extinction, and diversification across individual plant groups^10,19^. However, whether these patterns scale to the entire African flora remains unknown. Resolving this question requires continent-wide, time-calibrated phylogenies.

Beyond climatic and extinction dynamics, tectonic processes have been important drivers of diversification across plant and animal groups. In the Americas, the rise of the Andes triggered rapid speciation in TRF clades^36,37^. Similarly, in Africa, Cenozoic uplifts, especially in East Africa and along the eastern escarpment of southern Africa, generated pronounced topographic complexity, driving ecological heterogeneity, geographic isolation, and diversification of regional floras^10,15,38,39^. These links between deep-time geological and climatic changes are also evident in Madagascar, which despite its close proximity to Africa and shared geology, has evolved a high level of plant and animal endemism^40^. Yet, the relative contribution to large-scale continental patterns of vascular plant diversification remains insufficiently resolved. Together, these processes provide a framework to understand the evolutionary accumulation and distribution of African plant diversity.

Despite Africa’s rich evolutionary history, biodiversity faces escalating threats from deforestation, climate change, overexploitation, and desertification (Supplementary Fig. 2)^41–44^. Moreover, according to the IUCN Red List v.2025-2 (www.iucnredlist.org), many African plant species are classified as threatened – within the categories “Critically Endangered”, “Endangered” or “Vulnerable” – with numerous others listed as “Near Threatened” or “Data Deficient”^45^ (Supplementary Fig. 2), highlighting the urgent need for comprehensive research and effective conservation measures in Africa.

Understanding the ecological and evolutionary processes that generate and maintain biodiversity is crucial for identifying and conserving regions of high evolutionary potential^4,46^. Historically, biodiversity evaluations relied on species counts and endemism, with endemism being central to conservation priorities^47^. However, these metrics do not capture evolutionary history, leading to a shift toward phylogenetically informed approaches^48–52^. Phylogenetic metrics such as phylogenetic diversity (PD), phylogenetic endemism (PE), diversification rates (DR), and categorical analysis of neo- and paleo-endemism (CANAPE) provide complementary insights into the origins, persistence, and diversification of lineages, enabling the identification of regions that function as evolutionary refugia or centers of recent radiation^53–55^. Although these approaches have been applied to individual African clades and regional floras e.g.,^5,12,18,19,52,56–58^ a continent-wide synthesis integrating phylogenetic diversity, endemism, and diversification across the whole vascular flora remains lacking.

In this study, we generate a dated, genus-level tree of life for the vascular plants of Africa, integrate it with continental-scale distributional data to investigate the evolutionary and ecological processes that have shaped the continent’s botanical diversity. Building on expanded taxon sampling and distribution data, we develop a composite approach integrating phylogenetic diversity (PD), endemism, and diversification rates (DR, as an explicit diversity dimension) to evaluate biodiversity patterns across evolutionary timescales. Rather than proposing new conceptual models of African biome evolution, our aim is to synthesize and extend previous clade-specific phylogenetic insights within a continent-wide comparative context. Specifically, we aim to: 1) identify regions of high genus-level phylogenetic and evolutionary diversity of woody and herbaceous plant lineages across Africa using this composite framework; 2) assess spatiotemporal patterns of lineage diversification across Africa and their links to major macroevolutionary events; 3) evaluate the relative roles of abiotic and biotic drivers in shaping diversification across Africa’s biomes; and 4) identify conservation priority areas by integrating phylogenetic and diversification-based metrics. By linking Africa’s diverse ecological landscapes with its deep evolutionary history, this study provides an integrative macroevolutionary framework for plant biodiversity research and conservation.

## RESULTS

### Phylogenetic relationships, divergence time estimates and patterns

All orders and 285 families (98.27%) of African vascular plants were sampled (except Apodanthaceae, Hydroleaceae, Lonchitidaceae, Mayacaceae, and Triuridaceae). Our molecular dataset contained 21,815 species (45%) and 3,719 genera (93%) of the known vascular plants in Africa, of which 1,611 genera (40%) are endemic. The overall topology of our tree was largely consistent with the APG IV^59^, gymnosperm phylogenetic framework^60^, PPG I^61^, and recent global phylogenies, with only minor discrepancies in a few angiosperm relationships (Supplementary Fig. S4A). Eleven orders contained at least one endemic family.

Divergence time estimates indicate that most of African vascular plant genera (63%) originated during the Neogene and Quaternary (23 Ma to present) (Supplementary Fig. S5A), while 30% diverged during the Late Cretaceous and Paleogene (100–66 Ma and 66–23 Ma, respectively), and 7% originated between the Carboniferous and the Early Cretaceous (299–145 Ma) (Supplementary Fig. S5A). At higher taxonomic levels, most families (85%) originated during the Cretaceous (145–66 Ma), with a peak in the Late Cretaceous (100–66 Ma) (Supplementary Fig. S5A). South Africa, eastern Africa, and parts of northern Africa are centers of diversity for genera that originated before the Miocene (Supplementary Fig. S5B and D). In contrast, western Africa contains a higher proportion of genera that originated after the Miocene (Supplementary Fig. S5C and E).

Our divergence-time estimates were consistent with those from recent global-scale phylogenetic studies across taxonomic levels^62–65^ (Supplementary Fig. S6A–D), and showed minimal deviations in node ages for shared genera and families (Supplementary Fig. S6E–H), supporting the robustness of the phylogenetic framework and associated macroevolutionary inferences.

### Spatial patterns of diversity, endemism, age, and diversification rates

Poales, Gentianales, Fabales, Lamiales, Asterales, Malpighiales, and Asparagales accounted for most observed GR and PD (Supplementary Fig. S4B, C). However, DR varied independently of richness and PD. Asterales and Gentianales combined high GR and PD with high DR, while Fabales and Poales, despite being among the richest and most phylogenetically diverse orders, exhibited only moderate DR (Supplementary Fig. S4D). In contrast, Apiales, Brassicales, and Proteales showed comparatively high DR values despite lower GR and PD.

At the family level, Asteraceae with 360 genera (145 endemic); Poaceae, 307 (58); Rubiaceae, 159 (77); Apocynaceae, 136 (75); Aizoaceae, 116 (106); and Brassicaceae, 103 (20), combined high GR with elevated DR. In contrast, some families (e.g., Fabaceae) showed very high GR (306 genera, 134 endemic) but comparatively moderate DR. Conversely, Loranthaceae and Melastomataceae displayed high diversification rates despite relatively low GR, consistent with recent, lineage-specific diversification bursts. Within gymnosperms and ferns, Cupressaceae 4 (1) and Polypodiaceae 29 (none endemic) were the most diverse, but did not contribute substantially to high DR.

Overall, DR was moderately correlated with GR (BAMM: *r* = 0.63, DR statistic: *r* = 0.73) and PD (BAMM: *r* = 0.56, DR statistic: *r* = 0.66) while GR and PD were strongly correlated (*r* = 0.95; Supplementary Fig. S7A–C). Endemism across orders broadly mirrored GR and PD patterns. Gentianales (164/187 total endemic genera sampled), Asterales (160/211), Caryophyllales (150/158), Fabales (137/140), Asparagales (113/123), and Malpighiales (97/111) contained the highest numbers of endemic genera (Supplementary Fig. S4E).

Spatially, orders with high PD were concentrated within Africa’s major biodiversity regions spanning northern, western, eastern, and southern Africa (Supplementary Fig. S4, numbered 1–9). For example, Asparagales, Asterales, Brassicales, Caryophyllales, and Poales predominated in southern and northern Africa, whereas Malpighiales, Sapindales, and Gentianales were concentrated in western Africa; genera from the latter orders generally exhibited older divergence times (fig. S4). Across all vascular plants, five major regions of high diversity were consistently identified for both woody and herbaceous genera and all diversity metrics (Supplementary Fig. 1A–I): (1) the Mediterranean Basin (particularly areas surrounding the Atlas Mountains of Morocco, Algeria, and Tunisia), (2) the Guineo–Congolian region, (3) the Eastern Afromontane region (including the Ethiopian Highlands, Eastern Arc Mountains, and Albertine Rift), (4) the coastal forests of East Africa (from southern Kenya to northern Mozambique), and (5) the Cape Floristic Region (CFR), Succulent Karoo, and Maputaland–Pondoland–Albany. The Guineo– Congolian lowland forests showed comparatively lower richness of herbaceous genera (Fig. 1B, E, and H), while central and arid northern Africa showed only moderate diversity and endemism.

**Fig. 1.**
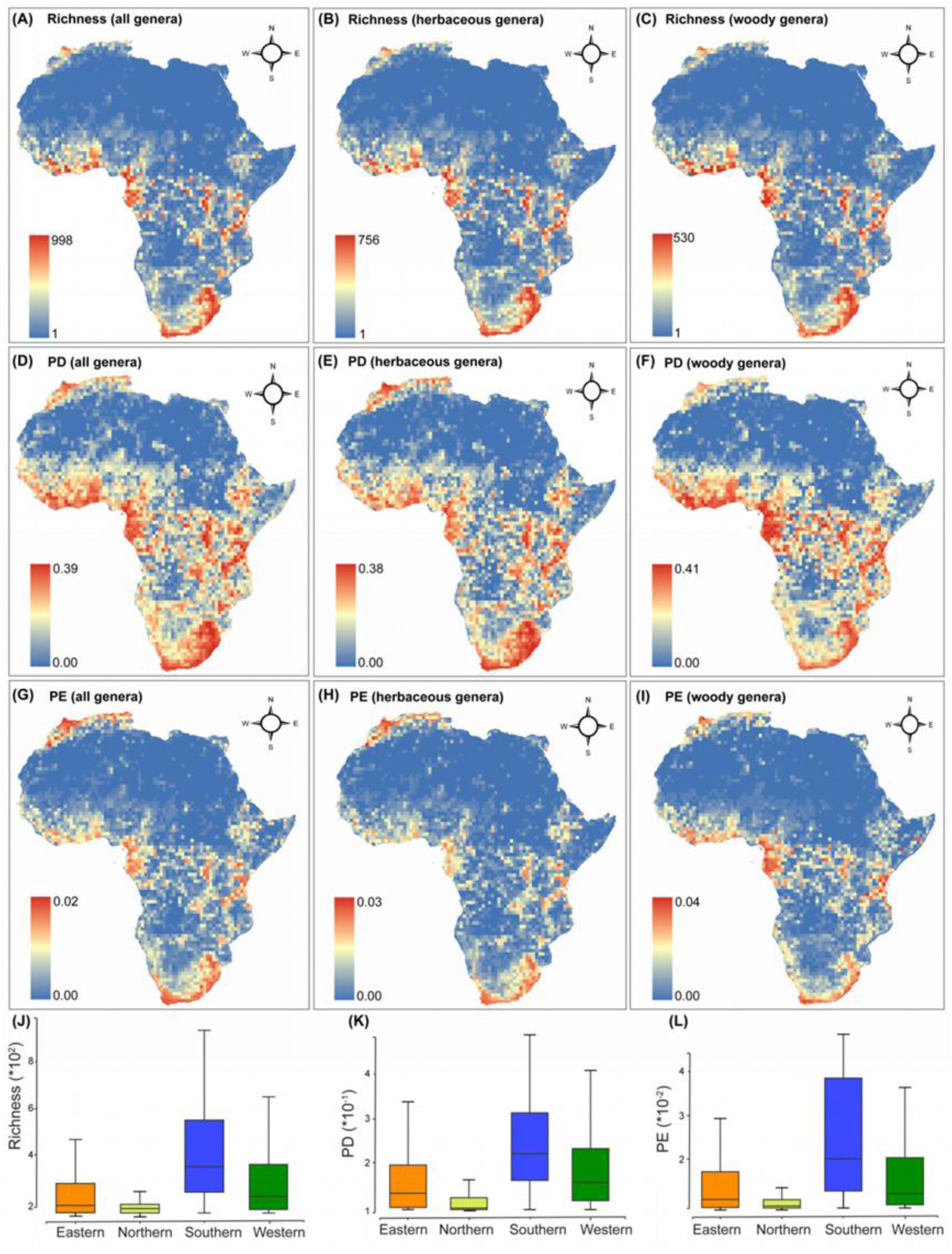
Patterns of diversity and endemism in African vascular plant genera. A–C, Richness for all genera (A), herbaceous genera (B), and woody genera (C). D–F, Phylogenetic diversity for all genera (D), herbaceous genera (E), and woody genera (F). G–I, phylogenetic endemism for all genera (G), herbaceous genera (H), and woody genera (I). The analyses include 3,719 vascular plant genera (herbaceous genera, *n* = 2,101; woody genera, *n* = 1,325; genera with both woody and herbaceous species, *n* = 293). J–L, Boxplots of generic richness (J), PD (K), and PE (L) of the three main biodiversity hotspots in Africa; median—solid line in the box, box—interquartile range (25% and 75%), whiskers—5% and 95% intervals. Maps are displayed in the WGS84 coordinate system.

Mapping tip rates (BAMM and DR statistic) revealed diversification patterns largely decoupled from present-day diversity (Fig. 2A–F). High DR occurred primarily in southern and eastern Africa, including the CFR, Succulent Karoo, Maputaland–Pondoland–Albany, the Eastern Afromontane region, and arid to semi-arid regions of northern and central Africa, while humid and tropical Guineo-Congolian forests showed consistently low DR despite high GR and PD. Across grid cells, PD showed a moderate positive association with DR (BAMM: *r* = 0.65; DR statistic: *r* = 0.65), and PE showed similar patterns (*r* = 0.63–0.64; fig. S8A–D), although these relationships reflected broad continental gradients rather than spatial congruence of hotspots.

**Fig. 2.**
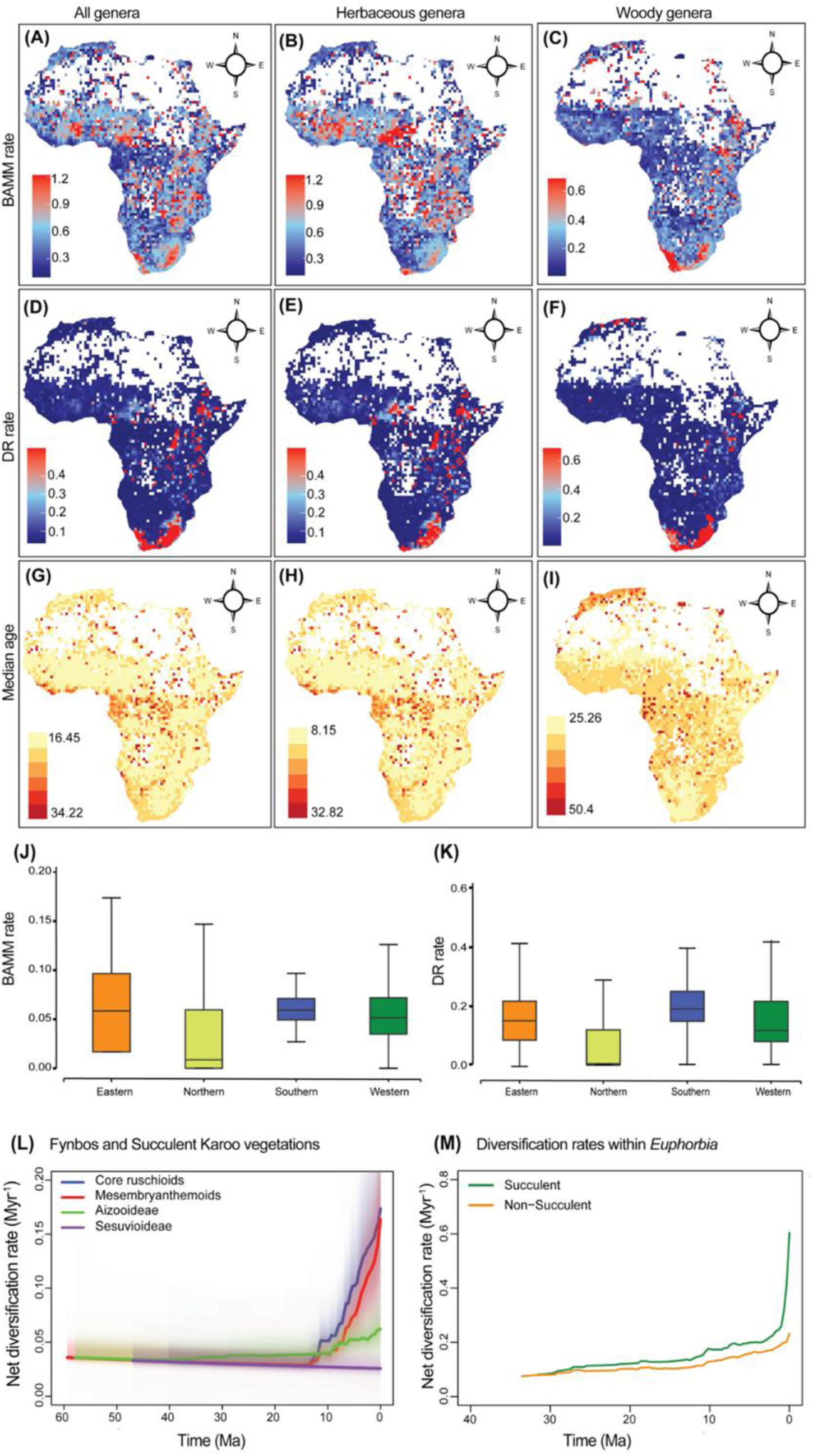
Spatial patterns of diversification and median age across vascular plant genera in Africa. Tip rates were estimated using Bayesian Analysis of Macroevolutionary Mixtures (BAMM) and diversification rate (DR) statistic (red = faster rates; blue = slower rates) for vascular plant genera across Africa. (A–C) BAMM rates for: all genera (A); herbaceous genera (B); and woody genera (C). (D–F) DR rates for: all genera (D); herbaceous genera (E); and woody genera (F). (G–I) Median ages for: all genera (G); herbaceous genera (H); and woody genera (I). J–K, Boxplots of BAMM (J) and DR statistic (K) of the three main biodiversity hotspots in Africa; median—solid line in the box, box—interquartile range (25% and 75%), whiskers—5% and 95% intervals. L–M, rate-through-time (RTT) plots showing net diversification rates for genera restricted to the Fynbos and Succulent Karoo vegetation (L), and for succulent and non-succulent genera within *Euphorbia* (M). All maps are displayed in the World Geodetic System 1984 (WGS84).

Assessment of SES-PD indicated phylogenetic clustering (low SES-PD) in the CFR, Succulent Karoo, and northern Africa, whereas strong phylogenetic overdispersion (high SES-PD) characterized the Guineo-Congolian region, Eastern Afromontane regions, and eastern South Africa for both herbaceous and woody genera (fig. S9A–C). Consistently, net relatedness index (NRI) showed positive clustering in arid and Mediterranean regions, while nearest taxon index (NTI) indicated clustering in southern Africa, particularly in the CFR and Succulent Karoo (fig. S9D–I).

Median lineage age analyses showed that western Africa, particularly the Guineo-Congolian region, harbors older lineages, whereas southern regions, especially the CFR and Succulent Karoo, are dominated by younger taxa (Figs. 2G–I and fig. S10–11). Tropical and humid regions consistently contained older genera than arid and temperate regions, while herbaceous and succulent genera exhibited younger median ages than woody and non-succulent genera, respectively (figs. S10M–O; S12A–D). CANAPE further identified centers of neo-endemism in eastern and southern Africa, including the CFR and Succulent Karoo, whereas paleo-endemism was concentrated in central and western regions (fig. S13A). Neo-endemic regions showed higher diversification rates and younger lineages compared to paleo-endemic regions (fig. S13B–E).

### Temporal diversification of African vascular plants

Analyses of diversification rates inferred from our molecular phylogeny revealed pronounced temporal heterogeneity within the African flora (Fig. 3A). Speciation and net diversification rates declined from the Devonian (ca. 380–359 Ma) to the Middle Jurassic (ca. 174–163 Ma) (Fig. 3B), remained relatively stable through much of the Cretaceous (145–66 Ma), and then increased gradually during the Cenozoic (66–0 Ma), particularly during the Neogene (23–2.6 Ma) and most prominently in the Miocene (23–5.3 Ma). Net diversification rates reached their highest values during the Miocene and Pliocene (5.3–2.6 Ma), reflecting a recent surge in lineage formation (Fig. 3B).

**Fig. 3.**
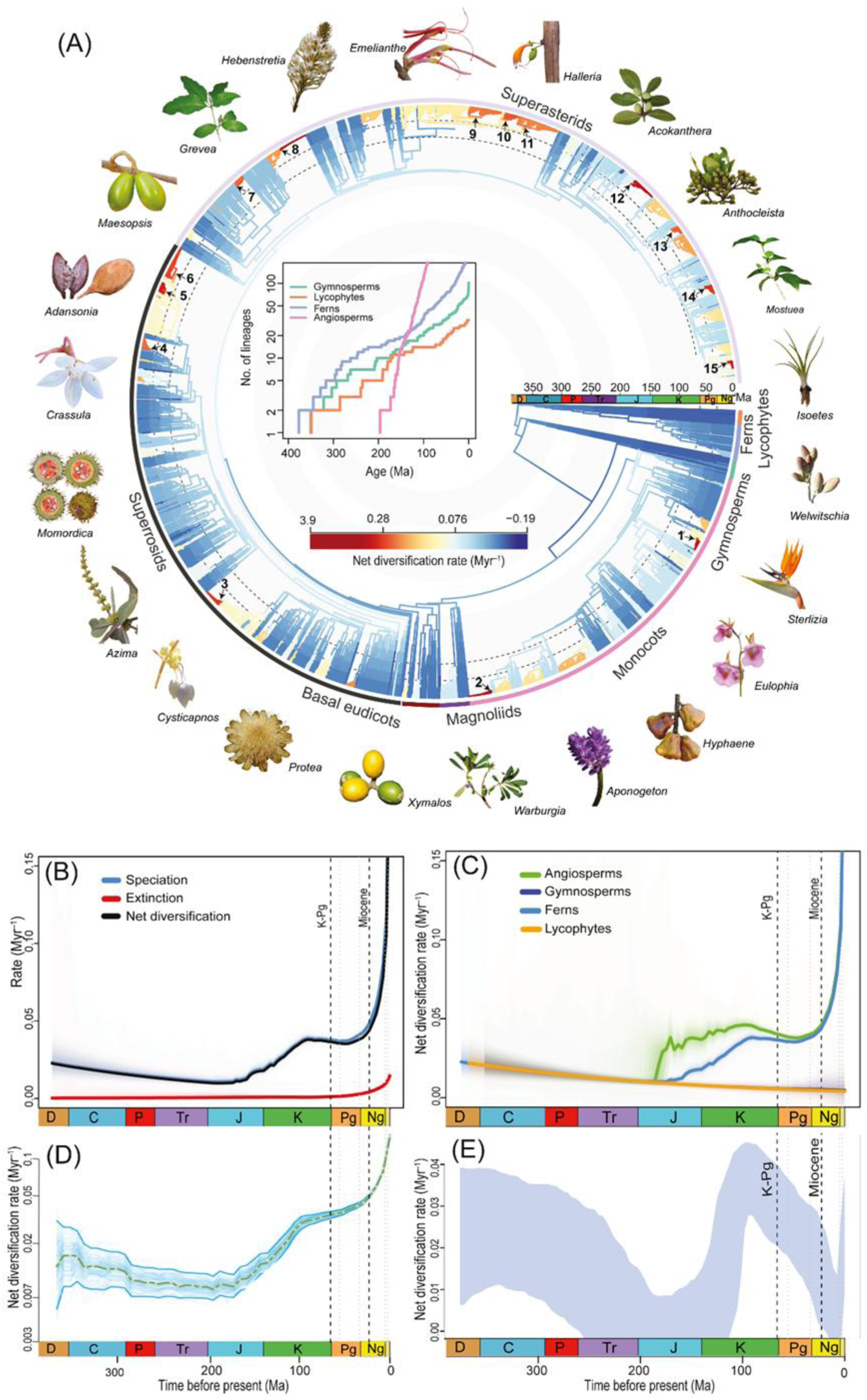
Net diversification dynamics of vascular plant genera in Africa. (**A**) Circular phylogenetic tree of vascular plant genera, colored by net diversification rates inferred using Bayesian Analysis of Macroevolutionary Mixtures (BAMM), with warmer colors (red to yellow) indicating higher diversification rates, and cooler colors (blue to light blue) indicating lower rates. Black arrows (labeled 1–15) mark the major diversification rate shifts detected across the tree. Representative genera are illustrated around the tree, grouped by major clades: angiosperms (including basal eudicots, magnoliids, monocots, and superasterids), gymnosperms, ferns, and lycophytes. The inset shows lineage-through-time (LTT) plots for the four major clades. (**B**) Rate-through-time (RTT) plot showing speciation, extinction, and net diversification rates for all vascular plant genera. Shaded areas represent 95% Bayesian credibility intervals. (**C**) RTT plot showing net diversification rates for each major vascular plant group, also with 95% Bayesian credibility intervals. (**D-E**) Net diversification rates from the cladogenetic diversification rate shift (CLADS) (D) and Compound Poisson Process on Mass-Extinction Times model (CoMET) (E) analysis for all vascular plant genera through time. Vertical dashed lines mark the Cretaceous– Paleogene boundary (K–Pg, 66 Ma) and the onset of the Miocene (23 Ma). Geological period abbreviations: D, Devonian; C, Carboniferous; P, Permian; Tr, Triassic; J, Jurassic; K, Cretaceous; Pg, Paleogene; Ng, Neogene. Photo credits: Bing Liu and Wyckliffe Omondi.

Among major clades, angiosperms exhibited the most pronounced rise in diversification, beginning near the Cretaceous–Paleogene boundary (ca. 66 Ma) and accelerating during the Miocene (Fig. 3C and fig. S14A). Ferns showed a moderate mid-Cenozoic rise followed by further increases from the Miocene, whereas gymnosperms and lycophytes maintained consistently low and stable rates throughout (Fig. 3C and fig. S14B–D). We identified 15 major shifts in net diversification rates within angiosperms during the Middle Miocene (ca. 15–10 Ma) (Fig. 3A, black arrows numbered 1–15), particularly in Aizoaceae, Fabaceae, Poaceae, Brassicaceae, Apocynaceae, Scrophulariaceae, and Melastomataceae lineages (Fig. 3A).

All diversification models (BAMM, ClaDS, CoMET, and RevBayes) rejected constant-rate diversification and inferred increasing speciation and net diversification toward the present (Figs. 3B–E; figs. S15–S16). CoMET further identified a pronounced diversification-rate shift within the last 10 Ma (2 ln BF > 6; fig. S16), consistent with the Miocene diversification pulses inferred by BAMM. Extinction rates remained comparatively low throughout most of the evolutionary history of African vascular plants, with only weak evidence for extinction-rate shifts and no support for discrete mass-extinction events (figs. S16–S17). These patterns were robust across alternative extinction-rate models (fig. S17A–C).

### Current environmental associations

The environmental effects on GR mirrored those on PE (fig. S18). Both were highest in areas with high mean annual precipitation (MAP) and low mean annual temperature (MAT), with MAT being the strongest predictor. In contrast, GR and PE decreased with increasing elevation, number of soil types, aridity, and human footprint.

DR showed weak but significant positive associations with higher elevation, greater human footprint, and increased aridity (figs. S19–S20). These patterns suggest that recent lineage diversification tends to be elevated in topographically complex, drier, and more disturbed environments, consistent with their relatively recent development in Africa. Precipitation seasonality and elevation had weak but significant effects on lineage age, with older lineages occurring in regions with more seasonal rainfall and lower elevations (fig. S21). Endemism patterns reinforced these contrasts: neo-endemism was concentrated in warmer, higher-elevation, and more arid regions with greater human impact, whereas paleo-endemism was associated with wetter, less arid, and less disturbed environments. Mixed endemism predominated under intermediate environmental conditions (fig. S22).

### Paleoenvironment- and trait-dependent diversification

Environment-dependent models indicated that elevation had the strongest effects on diversification, whereas CO_2_ and time had minimal influence (table S3). The best-supported elevation-dependent model (BElevVarD_ElevVar_EXPO; AICc = AICc = 31,407.396; Akaike weight = 0.985) showed a negative association between speciation and elevation variation (α = - 1.1274), and a similarly negative relationship between extinction and elevation variation (β = - 1.0885), resulting in an overall decline in net diversification with increasing elevation variability (fig. S23). Among temperature-dependent models, the best-supported model was BCSTDTempVar_EXPO (AICc = 31,510.473), which showed negative relationship between extinction and temperature variation (β = -0.3415), while effects on speciation were weak. In contrast, CO_2_-dependent and time-dependent models showed substantially weaker support and effect sizes.

Consistent with these model-based results, we detected negative correlation between global paleotemperature and atmospheric CO_2_ and tree-wide net diversification rates inferred using both BAMM and ClaDS (figs. S24, S25). Although elevation effects differed between methods— BAMM suggesting a weak positive association and ClaDS a negative one—analyses of younger clades (<50 Ma) consistently showed positive relationships between elevation variability and diversification, indicating stronger elevation-driven diversification at shallower evolutionary timescales (tables S4 and S5).

Trait-based analyses revealed that succulence, life form, aridity, and habitat significantly impact African lineage diversification (table S6). Succulent and herbaceous lineages had higher net diversification rates than non-succulent and woody lineages, respectively (Fig. 4A, B), while lineages adapted to arid, semi-arid, or temperate environments diversified faster than those in humid and tropical regions, also with no overlap in the 95% HPD intervals (Fig. 4C, D). Similar patterns were observed for speciation rates (fig. S26).

**Fig. 4.**
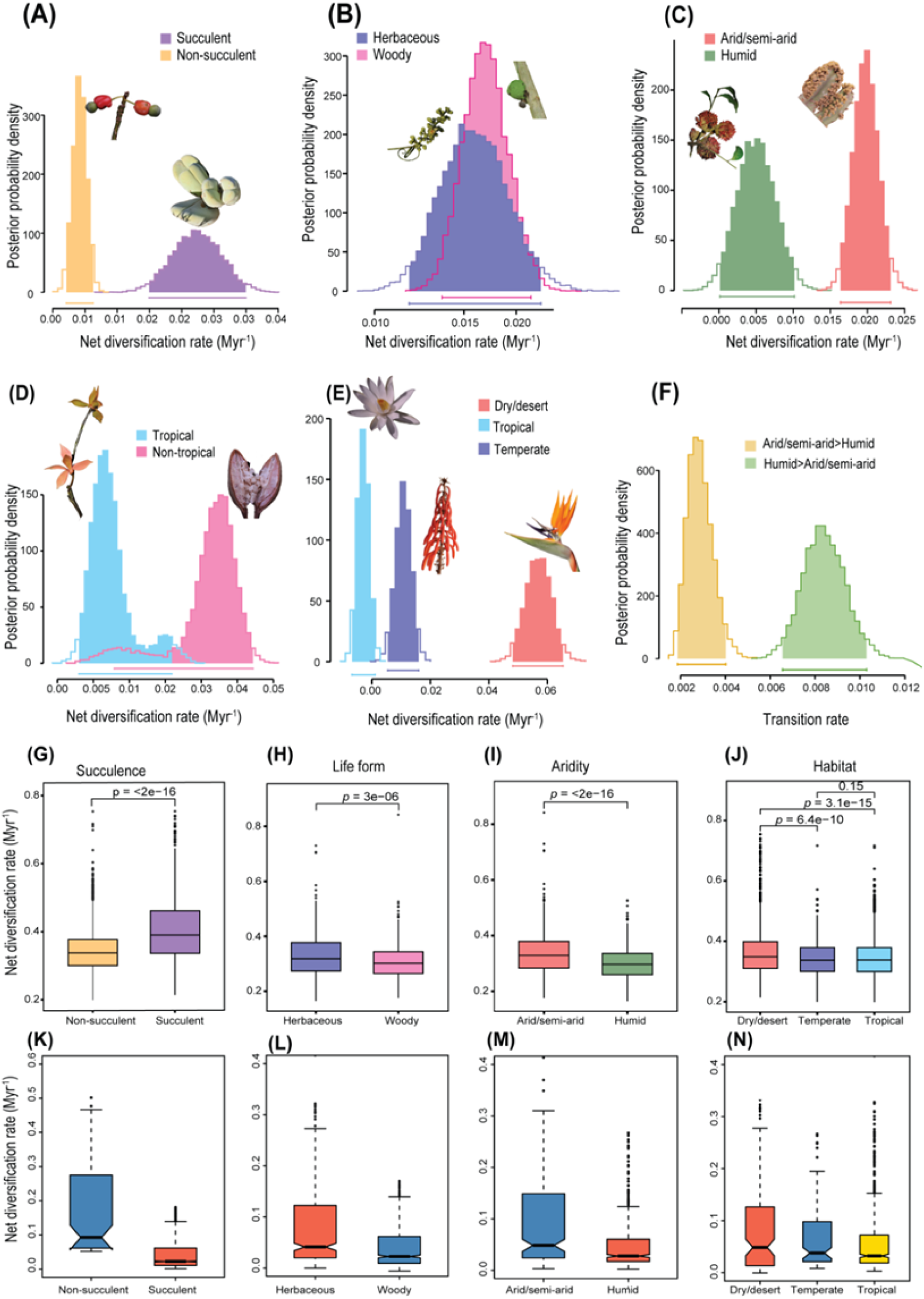
Biotic and abiotic drivers of diversification for vascular plant genera in Africa. (A–F) Diversification analyses using binary state speciation and extinction (BiSSE; A–F) and multistate speciation and extinction (MuSSE; E) models demonstrate the effects of two abiotic and two biotic traits on net diversification rate: Succulence (A), life form (B), aridity (C), and habitat (D, E). (F) transition rate between humid and arid/semi-arid as inferred by BiSSE. (G–H) Diversification analyses inferred by BAMM. (K–N) Net diversification rates inferred from cladogenetic diversification rate shift (ClaDS). Photo credits: Bing Liu.

Together, these results indicate that both abiotic (paleoelevation, paleotemperature) and biotic (traits, habitat preference) factors shaped diversification patterns across African vascular plants.

## DISCUSSION

By integrating phylogenetic diversity, endemism, and diversification-rate metrics across the African flora, our study provides a continent-scale evolutionary framework for understanding the assembly of African plant diversity. The results uncover consistent temporal and spatial signatures of diversification across lineages, showing that most extant genera arose during the Neogene, whereas family-level divergences peaked in the Late Cretaceous. Together, these patterns indicate deep evolutionary foundations coupled with uneven, lineage-specific diversification through time. Humid and tropical forests harbor older lineages, while arid and montane regions contain younger, rapidly diversifying assemblages. Hotspots of DR only partially overlap with areas of high PD, highlighting a decoupling between long-term lineage accumulation and recent radiations.

Phylogenetic clustering in arid and temperate areas suggests ecological filtering and niche saturation, whereas overdispersion in humid and tropical regions reflects the coexistence of distantly related lineages. Temporally, speciation rates increased markedly from the Miocene onward, peaking during the late Miocene–Pliocene, reflecting the influence of climatic shifts, aridification, and landscape reorganization. Herbaceous, succulent, and dry-adapted lineages diversified faster than woody or mesic taxa, highlighting the roles of persistence, ecological adaptation, and diversification in shaping Africa’s flora across space and time.

### Spatial decoupling of diversification rates and evolutionary history

Our analyses reveal that Africa’s principal reservoirs of evolutionary history (PD, PE, and GR) are concentrated in the northern Mediterranean Basin, Guineo-Congolian region, Eastern Afromontane and East African coastal forests, the CFR, Succulent Karoo, and Maputaland-Pondoland-Albany. Although these regions have previously been recognized as biodiversity hotspots e.g.,^52,56–58,66–68^, our continental synthesis reveals a clear macroevolutionary pattern: centers of phylogenetic diversity and endemism are largely distinct from centers of recent diversification. This spatial decoupling suggests that the contemporary distribution of African plant diversity reflects the interplay of two contrasting evolutionary processes—long-term persistence of ancient lineages and geographically distinct pulses of recent diversification. As a result, regions that maximize evolutionary history are not necessarily those generating biodiversity most rapidly today.

In the CFR, Eastern Afromontane, and semi-arid southern Africa, high diversity coincides with high DR, consistent with diversification associated with environmental heterogeneity and abiotic factors as climate, soils, fire regimes, and dispersal limitation^69,70^. Elevated DR in CFR and Succulent Karoo is consistent with recent radiations ca. 18 Ma, in clades including Iridaceae, Proteaceae, Restionaceae, and genera such as *Erica* and *Phylica*, coinciding with Miocene cooling and the onset of winter rainfall and fire regimes, facilitating arid- and fire-adapted lineages^71–73^.

By contrast, Guineo–Congolian forests exhibit high PD but low DR suggesting that diversity in these regions is primarily associated with long-term persistence rather than recent radiations^52,74^. Many tropical montane systems similarly combine deep evolutionary histories with localized and recent radiations^52,75^. Hyper-arid northern deserts and southern Africa exhibit low genus-level diversity but moderate DR, likely due to historical aridification and human impacts, including altered fire regimes and land-use^76^ (fig. S2).

This spatial decoupling between PD and DR can be summarized into three broad scenarios: (i) high PD–low DR regions, dominated by ancient lineages, consistent with tropical conservatism^77^; (ii) low PD–high DR regions shaped by recent diversification in arid and montane systems; and (iii) regions with both high PD and high DR, indicating prolonged stability punctuated by episodic radiations, as observed in the CFR and Eastern Afromontane regions^72,78^. Endemism patterns reinforce these scenarios. Neo-endemic taxa are associated with elevated DR in dynamic montane and semi-arid systems, reflecting recent *in situ* diversification and limited range expansion, whereas paleo-endemics contribute disproportionately to PD while exhibiting low contemporary DR, supporting the cradle–museum framework^48^. While these patterns align with previous regional and clade-based studies^52^, our continent-wide synthesis shows that cradle and museum dynamics often coexist spatially across African biomes.

Regions such as the CFR and Succulent Karoo exhibit phylogenetic clustering of young and closely related lineages such as *Pentameris* and *Muraltia*^79^, while the Guineo–Congolian region exhibits phylogenetic overdispersion, with co-occurring lineages that are distantly related and evolutionarily ancient e.g., *Strephonema* and *Pentadiplandra*^27,52^. CANAPE analyses indicate widespread mixed endemism across Africa, particularly in CFR, Succulent Karoo, and montane systems (Eastern Afromontane, Cape Fold, Atlas, and Rif Mountains), highlighting their dual roles as evolutionary cradles and museums^23,50,52^. The Guineo-Congolian forests show few cells as centers of endemism despite high richness, emphasizing the distinction between species richness and spatially restricted evolutionary uniqueness.

Lineage ages indicate that lowland tropical forests in western and central Africa harbor some of the oldest floristic elements, dating to the Cretaceous (dominated by Malpighiales, Sapindales, and Gentianales; figs. S4, S5, S10, and S11). This pattern is consistent with Linder^3^ and global studies on palms^80^ and Malpighiales^81^, and broader phylogenetic evidence for the ancient origins of tropical rainforest floras^82^, reinforcing the antiquity of tropical angiosperm-dominated forests. Together, these results show that the distribution of African plant diversity reflects both the long-term persistence of evolutionary lineages and geographically localized episodes of recent rapid radiations. Although consistent with previous phylogenetic and macroecological studies, our genus-level framework captures broad-scale evolutionary structure and may smooth finer-scale species-level patterns and recent diversification dynamics. Nevertheless, it reveals a pronounced spatial decoupling between regions that harbor deep evolutionary history and those characterized by rapid lineage accumulation.

### Trends in the diversification of vascular plants over time

Our results indicate that most families (95%) diversified more extensively during the Late Cretaceous. In comparison, only a smaller portion of genera (10%) show this pattern (Fig. S5), reflecting finer-scale temporal heterogeneity at lower taxonomic levels. The marked increase in diversification rates during the Neogene, particularly from the Miocene onward, is consistent with climatic and geological changes including East African Rift uplift, Sahara formation, and regional aridification^10,15^. Rift uplift likely promoted diversification by generating environmental heterogeneity, and geographic isolation through steep ecological gradients and scarp formation that isolated populations and facilitated species differentiation (fig. S28)^36,38,83^.

Angiosperms accelerated in diversification after the Miocene (Figs. 3C and fig. S14A), consistent with global Cenozoic radiations^64^. Gymnosperms, originating earlier (ca. 380 Ma) and flourishing during the Mesozoic^84^, declined from the Late Cretaceous to the present. This decline has been attributed to: (1) the rise of angiosperms, and (2) mass extinction events at 251.9, 201.3, and 66 Ma. Although extinction-rate estimation from phylogenies is challenging^85,86^, our episodic speciation–extinction analyses model detected extinction signals^87^ (fig. S16), but no major peaks coinciding with these mass extinctions (Figs. 3 and fig. S16B), aside from a slight increase in diversification rates around the Permian–Triassic boundary (ca. 251.9 Ma), consistent with previous findings^88^. While the precise timing of angiosperm origins remains debated e.g.,^65,89–92^, the temporal coincidence between angiosperm radiation and gymnosperm decline supports the “active displacement hypothesis”, whereby competitive replacement and Cenozoic climate cooling increased gymnosperm extinction rates in Africa and globally^88,93^.

In contrast, ferns showed higher diversification rates than gymnosperms and lycophytes, despite ancient origins, with most extant diversity arising from relatively recent radiations^94^. Their diversification closely tracks angiosperm expansion, likely reflecting ecological opportunities created by angiosperm-dominated forests in forest understories and canopies^95,96^*^;^*^97^. Consistent with this, shade-adapted fern lineages show higher diversification rates than open-habitat taxa (fig. S29), a pattern broadly aligned with fossil evidence indicating that closed-canopy angiosperm forests became widespread after ca. 65.5 Ma, despite sparse African fossil records and uncertainty in angiosperm origins^65,98^. Overall, fern diversification likely reflects a combination of environmentally driven extinction and opportunistic origination in expanding ecospace rather than niche availability alone^97^. Similar competitive dynamics may underlie the comparatively low diversification of African lycophytes, reflecting intrinsic constraints and competition with rapidly diversifying angiosperms, though these groups remain under-sampled^99^.

### Abiotic and biotic synergies in African plant evolution and diversification

DR hotspots are concentrated in arid and temperate regions, whereas humid and tropical forests exhibit lower DR but high PD (Fig. 2), consistent with the time-for-speciation hypothesis^55,100,101,102^. Genera restricted to arid and semi-arid regions are significantly younger (< 23 Ma) and exhibit higher net diversification and speciation, with CANAPE analyses revealing a concentration of neo-endemics in dry regions (Fig. 4 and figs. S12A, S22, S25C). Transitions from humid to dry habitats were more frequent than the reverse (Fig. 4F), indicating that recurrent aridification repeatedly created ecological opportunities for diversification, as also reported in African Melastomataceae and Fabaceae during the late Miocene^35,103^. BAMM identified major diversification shifts in several families, including Melastomataceae, within the clade of *Rosettea* and *Feliciotis* during the Middle Miocene (ca. 15–10 Ma). These patterns further support the role of Neogene aridification as a major driver of diversification by (1) generating novel ecological niches, increasing spatial heterogeneity, and (2) filtering for dry-adapted lineages^104,105^.

The expansion of the modern arid fynbos biome (from ca. 10 Ma) and the Succulent Karoo (< 5 Ma), which contain approximately one third of the world’s succulent species, are linked to increasing aridity along southern Africa’s west coast, driving mesic flora extinctions favoring arid-adapted lineages^72,78^. These shifts coincided with C_4_ photosynthesis bursts, increased diversification in herbaceous groups e.g., Poaceae and Asteraceae, formation of the Sahara Desert, and origin of succulent lineages like Didiereaceae in Madagascar^107–110^. Major radiations of ice plants (Ruschioideae) and diversification events in African *Euphorbia* also occurred between 11–2 Ma (Fig. 3L and M)^111, 112^.

Trait analyses reveal higher net diversification and speciation in succulent and herbaceous genera (Fig. 4 and fig. S25), reflecting shorter generation times, higher ploidy, and drought-adapted morphologies^113–115^. Morphological innovations, including wide-band tracheids (WBTs) and stone-like leaves in Aizoaceae genera (e.g., *Delosperma* and *Argyroderma*) likely facilitated Miocene diversification in arid habitats^112,116^. Today, arid, semi-arid, and most temperate regions across Africa are largely dominated by herbaceous, succulent, and dry-adapted lineages (fig. S30), underscoring the impact of Cenozoic aridification and open-habitat expansion on African floristic evolution.

Estimating extinction from molecular phylogenies is challenging^85,86^. Nevertheless, extinction-rate estimates have been used to investigate large-scale macroevolutionary patterns^65,102,117^. Our analyses indicate elevated extinction between ca. 50 and 6 Ma, followed by increased speciation (Fig. 3; figs. S15–S16). These extinction pulses coincide with global cooling and Neogene aridification, which fragmented humid habitats and promoted the expansion of more seasonal ecosystems, a pattern consistent with diversification dynamics observed in tropical rainforest clades during the Eocene–Miocene^10,19^.

Diversification–elevation relationships are strongly timescale dependent in our analyses. Models fitted to the full phylogeny show a negative association between diversification and elevation (ClaDS, RPANDA), consistent with deep-time constraints on high-elevation systems, whereas analyses restricted to younger clades (<50 Ma) reveal positive relationships, indicating pulses of diversification associated with mountain uplift and recent montane radiations^118^. Eastern Afromontane region exhibits high species richness, endemism, and elevated DR particularly in areas affected by repeated Cenozoic uplift (Figs. 1 and 2). Tectonic events, including Central African and East African uplifts (fig. S28), increased topographic complexity, promoting diversification and adaptive radiations in alpine lineages such as *Lobelia*, *Arabis alpina*, *Alchemilla*, and *Dendrosenecio*^10,119–121^. Similarly, tectonic stability and topographic complexity in the Cape Fold and Atlas Mountains promoted diversification in Proteaceae e.g., *Protea* and *Leucadendron*, Ericaceae, notably *Erica*^22,71,72,102^ and other montane and Mediterranean lineages, including *Erysimum*, and *Helianthemum*^122^, underscoring the role of mountain building and landscape heterogeneity in driving plant diversification across Africa. High-elevation systems thus may constrain diversification over long timescales, but act as arenas for recent and spatially restricted evolutionary radiations.

Overall, African plant diversification reflects the interplay of abiotic and biotic factors, consistent with macroevolutionary patterns observed in other regions. Aridification, topography, and climate fluctuations created ecological opportunities that, in combination with trait variation, have been associated with rapid diversification in arid and temperate regions, while humid and tropical areas accumulated older clades.

### Evolutionary insights for the conservation of African flora

This study provides a continent-wide synthesis of biodiversity patterns across Africa and shows how it accumulated through evolutionary processes (Fig. 5), providing an evolutionary framework for conservation strategies informed not only by species richness but also by the processes sustaining biodiversity^53^. Quantifying similarities and differences among biodiversity hotspots using complementary metrics—PD, CANAPE, and DR—enhances our ability to design protected-area networks capable of conserving multiple dimensions of biodiversity. Because these metrics capture complementary aspects of biodiversity, their combined use provides a more complete basis for conservation prioritization than species richness alone (Fig. 5 and fig. S26B).

**Fig. 5.**
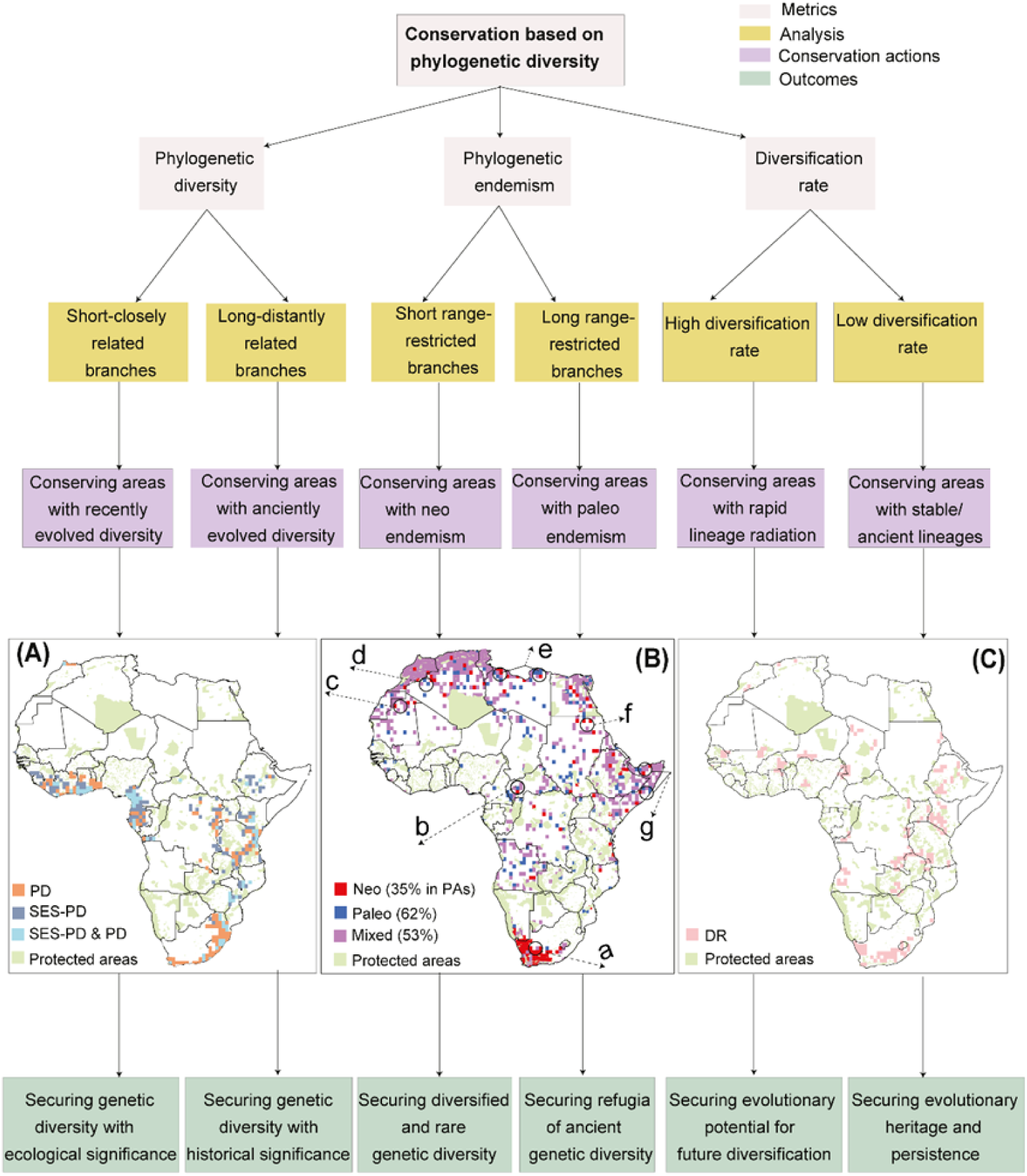
Evolutionary dimensions of biodiversity and their conservation relevance for African vascular plant genera. Conceptual framework (modified from González-Orozco & Parra-Quijano^161^) illustrating three complementary metrics that capture evolutionary diversity across African vascular plant genera and their implications for conservation. The figure highlights the advantages of incorporating evolutionary information into conservation planning. (A-C) Grid cells representing the top 5% of phylogenetic diversity (PD, brown) and standardized PD (SES-PD, blue), with overlapping cells in gray. Protected areas are highlighted in light green (A) (B) Conservation priorities identified using categorical analysis of neo- and paleo-endemism (CANAPE): neo-endemism (red), paleo-endemism (blue), and mixed endemism (purple). Letters (a–g) indicate some selected priority areas. (C) Map showing hotspots of diversification rate (DR; pink) overlaid on protected areas, illustrating regions of active evolutionary diversification relative to existing conservation coverage. Maps of protected areas are adapted from the World Database on Protected Areas (WDPA; https://www.protectedplanet.net/, accessed June 2025).

Overlaying these metrics with Africa’s protected-area network identifies several evolutionarily important yet unprotected regions, particularly in arid and semi-arid areas of Libya, Somalia, Sudan, Algeria, Mauritania, and parts of South Africa (Fig. 5B, labeled a–g). Because hotspots of rapid diversification often differ from areas of high PD or endemism (Fig. 5B and fig. S26), conservation strategies based on a single metric risk overlooking regions critical either for long-term lineage persistence or for ongoing evolution. Although arid regions are often perceived as species-poor and homogeneous^123^, our analyses demonstrate their central role in generating and maintaining Africa’s botanical diversity, consistent with diversification patterns observed in other seasonally dry and Mediterranean-type ecosystem globally.

Despite recent progress, most African landscapes remain outside formal protection (figs. S27J–K), necessitating inclusive solutions. Even within protected areas, it is crucial to ensure effective and community-led conservation, which like in Madagascar^124^can only be achieved through sufficient resource allocation. By explicitly linking evolutionary history, diversification dynamics, and spatial biodiversity patterns, this study provides an evolutionary perspective relevant to implementation of the “30 × 30” target of the Kunming–Montreal Global Biodiversity Framework, ensuring that protected areas safeguard both ancient lineages and the processes that continue to generate biodiversity.

### Limitations and Future Prospects

While our analyses provide a continent-wide synthesis of diversification dynamics in African vascular plants, diversification and extinction estimates derived from molecular phylogenies are sensitive to incomplete sampling and model assumptions, particularly for extinction, which remains difficult to estimate from extant-only trees^85,86^. Nevertheless, extinction-rate estimates have been widely used to investigate macroevolutionary patterns and extinction risk, despite ongoing methodological debate.^65,102,117^. In addition, stem ages may predate colonization of Africa, meaning that some inferred temporal patterns reflect lineage origins outside the continent rather than in situ diversification. These challenges are compounded by the sparse and uneven African plant fossil record, limiting direct validation of molecular inferences, especially for poorly sampled groups such as lycophytes.

Because our dataset is based on an Africa-only, genus-level phylogeny, most genera are represented by a single species. A genus-level approach was adopted to reduce taxonomic error, capture broad environmental signals, and maximize continental coverage, with sampling-fraction corrections applied where possible. Similar genus-level approaches have been employed in previous macroevolutionary studies^65,102^. Although this approach limits resolution of very recent species-level radiations, the major spatial and temporal patterns recovered here are robust across analytical frameworks and sensitivity analyses, consistent with previous macroecological studies^65,125^.

The growing availability of phylogenomic, functional traits, and paleobotanical data promises to transform our understanding of African plant evolution. Their integration within a continental-scale framework will provide a more complete reconstruction of diversification and extinction dynamics, reveal how environmental and ecological processes interact through time, and further illuminate the evolutionary mechanisms that have shaped one of the world’s most distinctive floras.

## Materials and Methods

### Species distribution data

We extracted species occurrences for vascular plants in Africa from the Global Biodiversity Information Facility (GBIF; www.gbif.org) and from 11 other online data sources: African Plant Database (APD, http://www.villege.ch/musinfo/bd/cjb/africa/recherche.php), POWO (http://www.plantsoftheworldonline.org), tropical African vascular plant database (RAINBIO, https://gdauby.github.io/rainbio/index.html), Botanical Database of Southern Africa (BODATSA, http://posa.sanbi.org/), Global Tree Search database (GlobalTreeSearch, https://tools.bgci.org/global_tree_search.php), *Protea* Atlas Project for Africa (https://www.proteaatlas.org.za/sugar5.htm), the World Flora Online (WFO, www.worldfloraonline.org), eflora Maghreb (https://efloramaghreb.org), North Africa Trees (https://www.northafricatrees.org/en/), Flore du Maroc, (http://www.floramaroccana.fr), and Integrated Digitized Biocollections (iDigBio, https://www.idigbio.org/). Additionally, we consulted regional, national, and local floras, as well as vascular plant checklists from both the literature and online sources, some of which were sourced by Qian et al.^5^.

Distributional data for each genus were compiled and manually checked for two main types of errors: (i) coordinate errors (e.g., missing, zero, country centroids, or clearly mislocated records) and (ii) taxonomic inconsistencies (e.g., outdated or unaccepted names). Corrections were applied where reliable information was available; otherwise, records that could not be corrected were excluded from the analyses. To ensure that all plant names in our records are accepted, we used “rWCVP”, a companion R package for the World Checklist of Vascular Plants (WCVP)^126^, to standardize the species names. We further improved data quality by removing common errors arising from georeferencing and data recording processes using the R package CoordinateCleaner^127^. Records with missing coordinates, country centroids, biodiversity institutions, zero coordinates, duplicated entries and obvious spatial outliers were removed. These processes ensured that only records from Africa were retained, excluding data from neighboring islands such as Madagascar, the Canary Islands, Madeira, Cape Verde, São Tomé and Príncipe, the Comoros, Seychelles, Mauritius, and Réunion.

By these procedures, we finally retained 1,001,645 district-level records for 48,137 species from 4,011 genera for vascular plants in Africa. Because occurrence records are spatially biased toward well-sampled regions (e.g., South Africa and parts of East Africa), which may influence estimates of richness and diversification patterns, we aggregated record into 100 km × 100 km equal-area grid cells to reduce sampling bias and improve comparability at continental scales. This grid size was selected to balance spatial resolution and sampling completeness while minimizing noise from uneven sampling. Grid cells were defined as polygon units and all raster layers were projected to the same coordinate system prior to extraction to avoid spatial misalignment. The final dataset’s coordinate information was projected using ArcGIS v.10.8. All spatial analyses were conducted in an equal-area projection (Africa Albers Equal Area Conic, AAEAC) to ensure unbiased area representation across latitudes, and results were subsequently visualized in geographic coordinates (WGS84). Information on species growth forms was assembled from RAINBIO, POWO^6^, Zanne et al.^128^, and Plant Trait Database (TRY)^129^. For taxa with unclear growth form status, information was obtained from regional floras and local botanical literature.

### Phylogenetic analyses

Sequences of six loci (*atpB*, *matK*, *ndhF*, *rbcL*, *trnL-trnF*, and ITS) were used to reconstruct the vascular plant phylogeny of Africa. The datasets included sequences from GenBank (http://www.ncbi.nlm.nih.gov/genbank/, accessed between August-October 2024). All sequences for genera native to Africa available in GenBank were downloaded using the ‘rentrez’ package in R v.3.2.4^130, 131^. We took infrageneric circumscriptions (subgenus and/or section) into consideration and gave priority to species with more targeted DNA sequences available in GenBank when choosing the representative species for each genus. Two liverworts (*Pellia endiviifolia* (Dicks.) Dumort. and *Aneura mirabilis* (Malmb.) Wickett & Goffi-net), and two mosses (*Syntrichia ruralis* (Hedw.) F.Weber & D. Mohr and *Physcomitrium patens* (Hedw.) Mitt.), were selected as outgroups.

The taxonomic accuracy of sequences was checked against recently published well-supported molecular phylogenetic results for infrafamilial relationships by conducting preliminary maximum likelihood (ML) analyses. If species with unreasonable placement were identified, these sequences were replaced or manually removed. These procedures were repeated until no unexpected lineage positions were detected. In cases where there were multiple sequences for a given species, only the longest sequence was kept for each genetic marker. If two sequences had equal lengths, the most recently published one was selected.

Sequences for each locus were aligned in three steps: first, using MAFFT v.7.305^132^; second, manually adjusting the alignment in BioEdit v.7.2.5^133^; and third, reorganizing the sequences based on their phylogenetic positions, followed by further adjustments where necessary. The final alignment included 3,249 sequences for *atpB*, 13,136 for *matK*, 6,123 for *ndhF*, 12,932 for *rbcL*, 9,640 for *trnL-trnF*, and 2,917 for ITS. To test the effect of missing data, we also generated a phylogenetic tree using *matK* and *rbcL* with less missing sequence data to test whether missing genetic data affect patterns of phylogenetic diversity, endemism, and DR. This sensitivity analysis explicitly evaluates whether missing genetic data bias downstream macroevolutionary inferences, thereby strengthening confidence in the robustness of our phylogenetic framework. The patterns of diversity and DR obtained from the *matK* and *rbcL* tree were highly congruent with those derived from the combined six-locus tree (r > 0.92, *p* < 0.001) (fig. S3), suggesting that our results are robust despite the higher missing data in the other four loci. Since high concordance mentioned above and the two-gene phylogeny had lower sampling density and resolution, the six-locus tree was used for all subsequent analyses due to its greater accuracy and completeness.

The concatenated six-locus matrix consisted of 21,815 species, representing 3,719 genera native to Africa from 285 families (93% of all vascular plant genera and 98.27% of all families). A partitioned (based on the six loci) ML analysis was conducted for the final concatenated data set using RAxML v.8.0.22^134^. Each locus was treated as an independent partition under the same GTR + GAMMA substitution model, with model parameters estimated separately for each partition. The optimal ML tree was inferred with 100 bootstrap replicates, with transition rates and base frequencies estimated using the GTR model, and among-site rate heterogeneity was modeled with a discretized gamma. FigTree v.1.3.19 (http://tree.bio.ed.ac.uk/software/figtree/)^135^ was used to view the resulting bipartition tree and generate a nexus formatted file suitable for our diversity analyses. We compared our topology with the orders according to Angiosperm phylogeny group (APG) IV^59^, Yang et al.^60^ for gymnosperms, and Pteridophyte Phylogeny Group (PPG) I^61^ for ferns, to confirm the phylogenetic position of each genus.

### Divergence time estimation and statistical analyses

We used the penalized likelihood method, as implemented in treePL^136^ (https://github.com/blackrim/treePL), to estimate divergence times of vascular plants in Africa, based on the optimal maximum likelihood phylogram obtained with RAxML v.8.0.22^134^, after excluding the outgroups. We validated the available fossils and selected 70 calibrations for dating analyses (table S1). Fossil calibrations were selected based on strict criteria, including reliable taxonomic placement, stratigraphic consistency, and prior use in published phylogenetic studies, to minimize calibration uncertainty. Most of the fossils we used have been widely applied in previous divergence time analyses e.g.,^49,62,64,65,101^. The ‘prime’ option was applied to identify the best optimization parameters, and a ‘thorough’ analysis was then carried out with the optimal parameters determined above (opt = 1, optad = 1 and optcvad = 4). Cross-validation based on χ² values indicated that a smoothing value of 1.0 provided the best model fit and was used in subsequent dating analyses. Confidence intervals for age estimates were calculated from 100 bootstrap replicates following Magallón et al.^62^ and Lu et al.^49^. Uncertainty in divergence time estimates was explicitly incorporated through bootstrap replicates, allowing downstream analyses to account for temporal uncertainty.

To evaluate the reliability of the estimated divergence times in our study, correlation analyses were conducted to compare our estimated divergence times with those of recent global-scale time trees^62–65^. These comparisons provide an external validation of our time estimates and help assess potential biases arising from taxon sampling or methodological differences. Among these, Smith and Brown^63^ has the largest and most up-to-date time tree for seed plants with 79,881 species. The stem ages of families and genera shared between these studies were extracted for Spearman’s rank correlation analyses in R following Lu et al.^49^. Briefly, for monophyletic families and genera, stem ages were extracted directly by tracing their stem node. For families and genera that are polyphyletic or paraphyletic, the stem age of each monophyletic lineage was extracted, and the oldest one was selected to represent the age of the family and genus. We then calculated the proportions of genera present in Africa with stem ages before and after the Miocene to explore divergence signatures of the African flora, recognizing that some lineages may have originated outside the continent before colonizing it. To further examine divergence patterns over evolutionary time, we estimated the number of genera originating within each five-million-year interval^49^. These descriptive time-slice analyses provide a complementary temporal context for interpreting the downstream diversification patterns inferred from rate-based methods, helping to highlight key periods of evolutionary change in relation to Africa’s geological and climatic history.

### Phylogenetic diversity and DR metrics

Using a time-calibrated phylogeny representing 93% of Africa’s vascular plant genera and extensive distribution records, we implemented an integrated analytical framework that jointly evaluates PD, endemism, and DR. By integrating multiple complementary metrics rather than relying on a single index, this framework reduces the risk of biased inference and allows a more comprehensive characterization of biodiversity patterns. Rather than defining new indices, this framework integrates established metrics to capture how plant biodiversity is structured across both evolutionary time and geographic space. We evaluated patterns of PD, phylogenetic structure, and endemism by calculating standardized metrics and conducting categorical analyses of phylogenetic endemism (CANAPE) following the methodological framework described by Mishler et al.^48^. Biodiverse v4.9^137^ was used to calculate generic richness (GR), phylogenetic diversity (PD) and phylogenetic endemism (PE) for vascular plant genera. Phylogenetic diversity (PD) was calculated as the sum of branch lengths connecting all genera within each grid cell^138^. PE was calculated as the sum of branch lengths connecting species present in a grid cell, weighted by the inverse of range size of their descendant species, to quantify the geographic concentration of evolutionary history^51^. To account for richness effects and identify assemblages with significantly higher or lower PD than expected by chance, we calculated the standardized effect size of phylogenetic diversity (SES-PD) using the R package *picante* under the taxa.labels null model with 999 randomizations. SES-PD was therefore calculated by dividing the difference between the observed (PD_observed_) and expected phylogenetic diversity (PD_random_) by the s.d. of the null distribution (s.d.(PD_random_)):

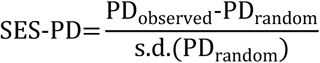

We also calculated the phylogenetic structure for vascular plant genera in Africa (clustering or overdispersion) using the net relatedness index (NRI) and the nearest taxon index (NTI)^79^ as follows:

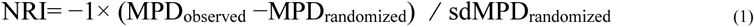

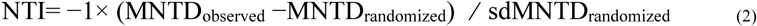

NRI was based on the mean phylogenetic distance (MPD), an estimate of the average phylogenetic relatedness between all possible pairs of taxa within a grid cell, while NTI was based on the mean nearest taxon distance (MNTD), an estimate of the mean phylogenetic relatedness between each species and its nearest relative within a grid cell. Positive values of NRI and NTI indicate phylogenetic clustering (taxa are more closely related than expected), whereas negative values indicate phylogenetic overdispersion (taxa are more distantly related than expected) in a grid cell^79^. Both SES.MPD and SES.MNTD were calculated using the R package “picante”^139^.

Importantly, PD and endemism patterns are interpreted alongside diversification dynamics, allowing recent lineage accumulation (captured by DR) to be assessed in the context of long-term evolutionary history. DR is therefore used here as a quantitative measure of recent lineage accumulation contributing to present-day diversity, with full details of DR estimation provided below.

### Macroevolution analyses through space and time

We analyzed macroevolutionary patterns to understand how vascular plant diversity in Africa has changed over time, aiming to detect variation in diversification rates, potential mass-extinction events, and the influence of environmental and trait factors. The following methods describe the approaches we used to explore these dynamics at both the tree-wide and lineage-specific scales. Additionally, we used several different models which allowed us to cross-check estimations^140^. Since each method differs in how speciation and extinction rates are estimated, our comparisons allowed us to explore the robustness and consistency of results across different model assumptions and implementations. Using multiple modeling approaches (BAMM, ClaDS, RevBayes) allows cross-validation of diversification estimates and reduces reliance on any single method, which is important given ongoing debates about diversification-rate inference.

### Time-dependent diversification analysis

First, we employed the Bayesian Analysis of Macroevolutionary Mixtures (BAMM) v.2.5.0^141^. BAMM implements the reversible jump Metropolis Coupled Markov Chain Monte Carlo (rjMCMC) method and allows for both time-dependent speciation rates as well as discrete shifts in the rate and pattern of diversification. This method enabled us to infer lineage-specific diversification rates (i.e., tip rates), identify shifts in diversification rates, and generate tree-wide diversification rate-through-time matrices and plots e.g.,^55^. We emphasize tip-based diversification metrics because they are less sensitive to model identifiability issues compared to deep-time rate estimates. BAMM running priors were configured using ‘setBAMMpriors’ function in BAMMtools v.2.1.6^141^ based on the time tree of Africa’s vascular plant genera after pruning the outgroups. As incomplete taxon sampling can bias estimation of speciation and extinction rates, we specified the sampling fraction of each genus based on the number of species present in Africa according to POWO^6^. Accounting for incomplete sampling at the genus level is critical to avoid biased estimates of speciation and extinction rates, particularly in under sampled clades.

The BAMM analysis was run for 40,000,000 generations, sampling every 1000 generations. We assessed convergence using the R package coda v.0.19-1^142^ after discarding 10% as burn-in, ensuring effective sample size (ESS) values for likelihood and the number of shifts exceeded 200. Rates-through-time plots were also generated for speciation (λ), extinction (µ), and net diversification (r) using ‘PlotRateThroughTime’ function in BAMMtools for the entire tree as well as specific clades. To extract species-specific speciation rate from BAMM results, we used the ‘getTipRates’ function. Moreover, the tip rates estimated from the DR statistic^55,143^ were also used as a complementary estimator to assess speciation dynamics and their potential environmental/trait dependence (e.g., climate factors and life form). For clarity, we use “DR” to refer to diversification rates estimated from both BAMM and the DR statistic, unless noted otherwise. Spatial patterns of diversification across Africa were then visualized via the R package dggridR v.2.0.3^144^, and median tip rates within each grid cell were used to produce a single representative value per cell. Here our main conclusions are mostly based on tip rates which are suggested to be less affected by identifiability issues e.g.,^145^.

Second, we used the cladogenetic diversification rate shift (ClaDS) model to estimate branch-specific diversification rates. ClaDS implements a birth–death model where speciation rates are inherited at speciation events, but with shifts drawn from a probability distribution parameterized by parental rates^146^. The extinction rate also varies across branches, but the turnover (i.e., speciation divided by extinction) is constant across branches. The model is computed using data augmentation in an MCMC with three chains. To assess convergence, the Gelman statistics are computed every 200 iterations, and the chains stop when these statistics drop below 1.05. We used the ClaDS model as implemented in the Julia package PANDA v.0.0.8^147^.

Third, we used RevBayes to estimate branch-specific diversification (speciation and extinction) rates (BSDR)^148^. The BSDR model is a birth–death model that divides time into small intervals, with speciation and extinction rates allowed to change at each interval. The new rates are drawn from a lognormal distribution, approximated using discrete rate categories to ease computation. We set up the model following the RevBayes tutorial (available at: https://revbayes.github.io/tutorials/divrate/branch_specific.html). Parameters were estimated using two rjMCMC chains of 4000 iterations each, sampling every 200 iterations. We performed several analyses with different prior specifications to check result consistency. Convergence was assessed with ESS > 200 using Tracer v.1.7^149^. For all BAMM, ClaDS, and RevBayes analyses, we set the sampling fraction to 0.90, while other parameters were kept at default values, unless stated otherwise in the text. The consistency of results across these independent frameworks was used as a key criterion for interpreting robust diversification patterns.

### Contemporary environmental data

We modeled the relationships between generic richness (GR), phylogenetic endemism (PE), and DR of vascular plants across Africa using a suite of predictor variables including climate, soil, topography, aridity, and human influence. Climate data were extracted from the CHELSA v.1.2 dataset (http://chelsa-climate.org/) at a 30 arc-seconds resolution, covering monthly temperature and precipitation from 1979–2013^150^. To assess soil heterogeneity, we used the Harmonized World Soil Database (www.fao.org/soils-portal/soil-survey/soil-maps-and-databases/harmonized-world-soil-database-v12/en/), calculating the number of distinct soil groups per grid cell using the using the “Zonal Statistics as Table” tool in ArcGIS Spatial Analyst (ArcGIS v.10.8)^151^.

Topographic data were obtained from the Shuttle Radar Topography Mission (SRTM30) Digital Elevation Model (DEM) version 2 (https://www2.jpl.nasa.gov/srtm/), which provided key metrics such as average elevation, elevation variation, and topographic roughness. The aridity index was derived from the Global Aridity Index and Potential Evapotranspiration Database (Global-AI_PET_v3: https://doi.org/10.6084/m9.figshare.7504448.v5) at a resolution of 30 arc-seconds. For measuring the impact of anthropogenic factors, we extracted the global human footprint (HFP) for each grid cell from the Global Human Footprint Database (https://doi.org/10.5061/dryad.052q5). All raster datasets were projected into the same AAEAC coordinate system as the analytical grid, and values were summarized to the 100 km × 100 km grid polygons using the “Zonal Statistics as Table” tool in ArcGIS v.10.8 to avoid resampling artifacts associated with raster reprojection.

To avoid potential collinearity among the climatic variables, we first conducted a correlation analysis, selecting three predictors where the inter-correlation was below an absolute value of 0.7. Reducing multicollinearity improves model interpretability and prevents inflation of parameter estimates. These included (i) mean annual temperature, (ii) annual precipitation, and (iii) precipitation seasonality. Following this, we selected eight variables for the final analysis, which included the three previously mentioned climatic predictors, as well as (iv) number of soil types, (v) average elevation, (vi) topographic roughness, (vii) aridity index, and (viii) human footprint (table S2). Most of these variables have also been commonly considered as the most important factors determining distributions and diversity of plants and animals. Regression models and data transformations followed steps and R scripts used by Antonelli et al.^83^, with GR, PE, DR, and the eight environmental variables acting as the response and predictor variables, respectively.

### Paleoenvironment-dependent diversification analysis

Abiotic environmental factors have long been regarded as important drivers of macroevolutionary dynamics^152^. Therefore, we investigated the impact of environmental change on diversification of African flora using paleoenvironment-dependent models in RPANDA 1.9^145^. We used proxies for three environmental variables commonly linked to continental-scale macroevolutionary diversification and that capture major aspects of paleoclimatic change in Africa: paleotemperature and atmospheric CO_2_ [datasets provided in^88^], and past elevation change from Scotese and Wright^153^ (table S2). These variables were selected because they represent major axes of environmental change (temperature, atmospheric composition, and topography) known to influence macroevolutionary processes at continental scales. Because the Scotese and Wright dataset reports global paleoelevations through 0–540 Ma, we spatially filtered the coordinate grid to retain only points falling within Africa, ensuring that the resulting elevation proxy reflects environmental dynamics specific to our study region. We specifically hypothesized that variation in these three variables may have influenced the diversification of vascular plant genera in Africa. By adapting the method and scripts from Condamine et al.^140^, we developed four hierarchical models for each environmental proxy. Each model assumed exponential dependencies of speciation and extinction as follows: 1) speciation varies with the paleoenvironmental variable and no extinction; 2) speciation varies with the paleoenvironmental variable and extinction is constant; 3) speciation is constant and extinction varies with the paleoenvironmental variable; and 4) both speciation and extinction vary with the paleoenvironmental variable. In the models, the value of α (β) indicates how strongly speciation (extinction) rate changes in response to environmental variation, and the sign of α (β) indicates a positive (negative) environmental effect on the diversification rate trend. We then computed the AIC for each model fit and plotted the net diversification rate over time based on the model with the lowest AIC. Model comparison using AIC allows objective selection of the best-supported environmental driver of diversification dynamics.

### Detecting mass extinction events

To examine tree-wide variation in diversification rate and the occurrence of mass extinctions during the evolution of the African flora, we used the Compound Poisson Process on Mass-Extinction Times model (CoMET)^87^, as implemented in the R package TESS v.2.1.2^154^. This model estimates tree-wide speciation and extinction rates, as well as the probability of several tree-wide events occurring, such as shifts in speciation rate, shifts in extinction rate, or sudden extinction events (i.e., when several lineages go extinct with a prior probability). We note that detecting extinction from extant-only phylogenies is inherently challenging^85,86^; therefore, results from CoMET should be interpreted cautiously and in conjunction with other analyses. The events are estimated under the independent Compound Poisson Process. Parameters are estimated using an rjMCMC method over various episodic birth–death models. Model confidence is assessed by computing the Bayes factor (BF) between each model^87^. For the CoMET analysis, we set the priors on the number of extinction events and number of expected changes to 2. Note that the prior on the number of events does not impact the results as the models are compared using BFs^154^.

We applied the R scripts from Condamine et al.^140^ and ran the rjMCMC for 10 million steps, discarding the initial 25% as burn-in. We considered support for a rate shift or for a sudden extinction event as substantially significant for 2lnBF > 2, strongly significant for 2lnBF > 6, and decisive for 2lnBF > 10^154,155^. To check the convergence of interval-specific parameters, we used the ‘tess.plot.singlechain.diagnostics’ function of the TESS package. We further used the CRABS method^156^ to explore the congruence class induced by our tree and various extinction-rate and speciation-rate scenarios to identify whether the TESS-based diversification trend is robust. The use of congruence class exploration (CRABS) helps to assess whether inferred diversification trends are robust to alternative extinction scenarios. Using the TESS inference of global diversification dynamics, we tested five scenarios of extinction rate and two scenarios of speciation rate.

### Trait-dependent diversification analyses

To evaluate whether diversification rates are influenced by traits or habitat shifts, we tested trait-dependent diversification rates for African lineages with traits such as aridity, life form, succulence, and habitat type (classified as dry/desert, temperate and tropical regions). We applied the BiSSE (Binary State Speciation and Extinction) and MuSSE (Multi-State Speciation and Extinction) models^157^ to examine trait-dependent diversification. These analyses were conducted using the R package diversitree v.0.9-10^158^, fitting five distinct models: (1) a full model allowing all parameters to vary independently; (2) a model constraining extinction rates (μ) to be equal across states while allowing speciation rates (λ) and transition rates (q) to vary; (3) a model constraining speciation rates (λ) to be equal while allowing extinction rates (μ) and transition rates (q) to vary; (4) a model constraining extinction rates (μ) and transition rates (q) to be equal while allowing speciation rates (λ) to vary; and (5) a model constraining transition rates (q) to be equal while allowing speciation (λ) and extinction rates (μ) to vary. Model comparisons were performed using likelihood ratio tests based on a chi-square distribution and AICc (Akaike Information Criterion corrected; Akaike, 1974). In addition to life forms mentioned in the “Species distribution data” section, trait data for habitat, aridity, and succulence were compiled from online floras, including eFloras (http://www.efloras.org/) and Plants of the World Online (https://powo.science.kew.org/; accessed June 2025). Additionally, genera were classified as succulent or non-succulent following the criteria and references provided by Dimitrov et al.^101^.

Because trait-dependent diversification models can suffer from high false-positive rates, results were interpreted conservatively and compared across alternative model constraints.

### Locating diversity hotspots and conservation priorities

We detected areas of potential conservation priority (i.e., areas with high phylogenetic diversity but not currently protected) in Africa by overlaying PD, standardized effect size of phylogenetic diversity (SES-PD), and DR hotspots and CANAPE results on maps of protected areas using ArcGIS 10.8. Maps of protected areas were downloaded from the World Database on Protected Areas (WDPA, https://www.protectedplanet.net accessed June 2025); the definition of protected areas followed those of the International Union for Conservation of Nature (IUCN) and the Convention on Biological Diversity (CBD). To define hotspots, we applied a common threshold, identifying the top 5% and 30% of grid cells in each category.

To avoid conflating statistical significance with arbitrary percentile thresholds, hotspot definitions differed according to metric type. For observed PD and DR, hotspot cells were defined as the top 5% of grid cells, representing concentrated centers of evolutionary diversity and recent diversification, respectively. For SES-PD, values were interpreted using statistical significance thresholds (|z| > 1.96, α = 0.05) based on null model expectations, while CANAPE results identified statistically significant centers of neo- and paleo-endemism following Mishler et al. (2014). This approach has been widely recognized for its ability to represent a substantial proportion of terrestrial biodiversity across various taxonomic groups^50,53,159,160^. The 30% threshold was used only for exploratory visualization of broader spatial patterns and not for statistical inference or conservation designation. Only cells outside existing protected areas were highlighted as priority regions. Cells within protected areas were excluded to highlight unprotected regions of evolutionary importance, rather than to define evolutionary significance itself. This step was applied solely for spatial overlap analysis and does not affect the identification of evolutionary significance or hotspot detection.

By integrating phylogenetic diversity, phylogenetic structure, endemism, and diversification metrics, this framework captures both deep evolutionary history and recent lineage accumulation, thereby complementing traditional species richness–based conservation prioritization.

## ACKNOWLEDGMENTS

We thank Anthony Verboom (University of Cape Town) and Fabien Condamine (Institut des Sciences de l’Évolution de Montpellier) for their insightful comments and constructive suggestions that substantially improved the manuscript.

## Author contributions

Conceptualization: W.O.O. and M.S.; Methodology: W.O.O., X.W., and L.Z.; Investigation: L.B., M.K.G., R.W.G., A.A., Q.F.W., and Z.D.C.; Writing – Original Draft: W.O.O. and M.S.; Writing – Review & Editing: M.K.G., A.A., Q.F.W., Z.D.C., and M.S.; Funding Acquisition: W.O.O. and M.S.; Data Curation: X.W., L.Z., X.Z., T.Z., Y.L., and Z.L.; Supervision: A.A., Q.F.W., C.D.Z., and M.S.

## Data availability statement

All data needed to evaluate the conclusions of this paper are available in the paper or the Supplementary Materials.

## Supplementary Materials

Materials and Methods

Figs. S1 to S30

Tables S1 and S6

References

## REFERENCES AND NOTES

1. F. White, The vegetation of Africa. Unesco, Paris. (1983).

2. L. Mucina, M. C. Rutherford (eds.), The vegetation of South Africa, Lesotho and Swaziland. Strelitzia 19, South African National Biodiversity Institute, Pretoria (2006).

3. H. P. Linder, The evolution of African plant diversity. Front. Ecol. Evol. 2, 38 (2014).

4. N. Myers, R. A. Mittermeier, C. G. Mittermeier, G. A. da Fonseca, J. Kent, Biodiversity hotspots for conservation priorities. Nature 403, 853–858 (2000).

5. H. Qian, Y. Zhou, J. Zhang, Y. Jin, T. Deng, S. Cheng, A synthesis of botanical informatics for vascular plants in Africa. Ecol. Inform. 64, 101382 (2021).

6. POWO, Plants of the World Online. Facilitated by the Royal Botanic Gardens, Kew (2025); available at https://powo.science.kew.org (accessed 19 September 2024).

7. Y. Malhi, T. A. Gardner, G. R. Goldsmith, M. R. Silman, P. Zelazowski, Tropical forests in the Anthropocene. Annu. Rev. Environ. Resour. 39, 125–159 (2014).

8. L. N. Hoveka, M. van der Bank, B. S. Bezeng, T. J. Davies, Identifying biodiversity knowledge gaps for conserving South Africa’s endemic flora. Biodivers. Conserv. 29, 2803– 2819 (2020).

9. S. McLoughlin, The breakup history of Gondwana and its impact on pre-Cenozoic floristic provincialism. Aust. J. Bot. 49, 271–300 (2001).

10. T. L. P. Couvreur, G. Dauby, A. Blach-Overgaard, V. Deblauwe, S. Dessein, V. Droissart, O. J. Hardy, D. J. Harris, S. B. Janssens, A. C. Ley, B. A. Mackinder, Tectonics, climate and the diversification of the tropical African terrestrial flora and fauna. Biol. Rev. 96, 16–51 (2021).

11. H. P. Linder, H. M. de Klerk, J. Born, N. D. Burgess, J. Fjeldså, C. Rahbek, The partitioning of Africa: Statistically defined biogeographical regions in sub-Saharan Africa. J. Biogeogr. 39, 1189–1205 (2012).

12. Q. Zhang, J. F. Ye, C. T. Le, D. N. Mwithukia, R. N. Rabarijaona, W. O. Omollo, L. M. Lu, B. Liu, Z. D. Chen, New insights into the formation of biodiversity hotspots of the Kenyan flora. Divers. Distrib. 28, 2696–2711 (2022).

13. G. A. Verboom, H. P. Linder, F. Forest, V. Hoffmann, N. G. Bergh, R. M. Cowling, N. Allsopp, J. F. Colville, Cenozoic assembly of the Greater Cape flora, pp. 93–118 (2014).

14. J. N. Wan, S. W. Wang, A. R. Leitch, I. J. Leitch, J. B. Jian, Z. Y. Wu, H. P. Xin, M. Rakotoarinivo, G. E. Onjalalaina, R. W. Gituru, C. Dai, The rise of baobab trees in Madagascar. Nature 629, 1091–1099 (2024).

15. N. Werner, Z. Wang, L. Werdelin, Q. Zhang. East African uplift as a catalyst for Middle Miocene faunal transitions. Sci. Adv. 11, eadx6569 (2025).

16. Davis CC, Bell CD, Fritsch PW, Mathews S. 2002. Phylogeny of Acridocarpus-Brachylophon (Malpighiaceae): implications for Tertiary tropical floras and Afroasian biogeography. Evolution 56: 2395–2405.

17. Couvreur TLP, Chatrou LW, Sosef MSM, Richardson JE. 2008. Molecular phylogenetics reveal multiple Tertiary vicariance origins of the African rain forest trees. BMC Biology 6: 54.

18. Brée B, Helmstetter AJ, Bethune K, Ghogue J-P, Sonké B, Couvreur TLP. 2020. Diversification of African rainforest restricted clades: Piptostigmateae and Annickieae (Annonaceae). Diversity 12: 227

19. L. P. M. Dagallier, F. L. Condamine, T. L. P. Couvreur, Sequential diversification with Miocene extinction and Pliocene speciation linked to mountain uplift explains the diversity of the African rain forest clade Monodoreae (Annonaceae). Ann. Bot. 133, 677–696 (2024).

20. Plana V. 2004. Mechanisms and tempo of evolution in the African Guineo-Congolian rainforest. Philosophical Transactions of the Royal Society B: Biological Sciences 359: 1585–1594.

21. Morley RJ. 2011. Cretaceous and Tertiary climate change and the past distribution of megathermal rainforests. In: Bush M, Flenley J, Gosling W, eds. Tropical rainforest responses to climatic change. Berlin: Springer, 1–34.

22. G. A. Verboom, N. G. Bergh, S. A. Haiden, V. Hoffmann, M. N. Britton, Topography as a driver of diversification in the Cape Floristic Region of South Africa. New Phytol. 207, 368– 376 (2015).

23. H. P. Linder, G. A. Verboom, The evolution of regional species richness: the history of the southern African flora. Annu. Rev. Ecol. Evol. Syst. 46, 393–412 (2015).

24. Pan AD, Jacobs BF, Dransfield J, Baker WJ. 2006. The fossil history of palms (Arecaceae) in Africa and new records from the Late Oligocene (28–27 Mya) of north-western Ethiopia. Botanical Journal of the Linnean Society 151: 69–81.

25. Faye A, Pintaud J-C, Baker WJ, Vigouroux Y, Sonke B, Couvreur TLP. 2016. Phylogenetics and diversification history of African rattans (Calamoideae, Ancistrophyllinae). Botanical Journal of the Linnean Society 182: 256–271.

26. E. D. Currano, B. F. Jacobs, A. D. Pan, Is Africa really an “odd man out”? Evidence for diversity decline across the Oligocene–Miocene boundary. Int. J. Plant Sci. 182, 551–563 (2021).

27. Couvreur TLP. 2015. Odd man out: why are there fewer plant species in African rain forests? Plant Systematics and Evolution 301: 1299–1313.

28. Richards, P. W. (1973). Africa, the ‘odd man out’. In Tropical Forest Ecosystems of Africa and South America: A Comparative Review (eds B. J. Meggers, E. S. Ayensu and W. D. Duckworth), pp. 21–26. Smithsonian Institution Press, Washington, DC.

29. Morley RJ. 2000. Origin and evolution of tropical rain forests. Chichester: John Wiley & Sons Ltd.

30. J. A. Coetzee, J. Praglowski, Winteraceae pollen from the Miocene of the southwestern Cape (South Africa). Grana 27, 27–37 (1988).

31. S. Nilsson, J. H. Coetzee, E. Grafström, On the origin of the Sarcolaenaceae with reference to pollen morphological evidence. Grana 35, 321–334 (1996).

32. A. Antonelli, A. Zizka, D. Silvestro, R. Scharn, B. Cascales-Miñana, C. D. Bacon, An engine for global plant diversity: Highest evolutionary turnover and emigration in the American tropics. Front. Genet. 6, 130 (2015).

33. Uno KT, Polissar PJ, Jackson KE, deMenocal PB. 2016. Neogene biomarker record of vegetation change in eastern Africa. Proceedings of the National Academy of Sciences 113: 6355–6363.

34. Sepulchre, P., Ramstein, G., Fluteau, F., Schuster, M., Tiercelin, J. J. & Brunet, M. (2006). Tectonic uplift and eastern Africa aridification. Science 313, 1419–1423.

35. Veranso-Libalah, M. C., Kadereit, G., Stone, R. D. & Couvreur, T. L. P. (2018). Multiple shifts to open habitats in Melastomateae (Melastomataceae) congruent with the increase of African Neogene climatic aridity. Journal of Biogeography 45, 1420–1431.

36. C. Hoorn, F. P. Wesselingh, H. ter Steege, M. A. Bermudez, A. Mora, J. Sevink, I. Sanmartín, A. Sanchez-Meseguer, C. L. Anderson, J. P. Figueiredo, C. Jaramillo, Amazonia through time: Andean uplift, climate change, landscape evolution, and biodiversity. Science 330, 927–931 (2010).

37. L. P. Lagomarsino, F. L. Condamine, A. Antonelli, A. Mulch, C. C. Davis, The abiotic and biotic drivers of rapid diversification in Andean bellflowers (Campanulaceae). New Phytol. 210, 1430–1442 (2016).

38. J. Bentley, G. A. Verboom, N. G. Bergh, Erosive processes after tectonic uplift stimulate vicariant and adaptive speciation: Evolution in an Afrotemperate-endemic paper daisy genus. BMC Evolutionary Biology 14, 27 (2014).

39. V. Hoffmann, G. A. Verboom, F. P. Cotterill, Dated plant phylogenies resolve Neogene climate and landscape evolution in the Cape Floristic Region. PLoS ONE 10, e0137847 (2015)

40. A. Antonelli, R. J. Smith, A. L. Perrigo, A. Crottini, J. Hackel, W. Testo, H. Farooq, M. F. Torres Jiménez, N. Andela, T. Andermann, A. M. Andriamanohera, Madagascar’s extraordinary biodiversity: Evolution, distribution, and use. Science 378, eabf0869 (2022).

41. J. D. D. Vidal Junior, A. Antonelli, C. Carbutt, V. R. Clark, T. Fremout, C. Chapano, I. Chelene, D. Chuba, T. W. Gole, C. Langa, B. Loeuille, Late 21st-century climate- and land-use–driven loss of plant diversity in African mountains. Glob. Change Biol. 31, e70492 (2025).

42. T. Stévart, G. Dauby, P. P. Lowry, A. Blach-Overgaard, V. Droissart, D. J. Harris, B. A. Mackinder, G. E. Schatz, B. Sonké, M. S. Sosef, J. C. Svenning, A third of the tropical African flora is potentially threatened with extinction. Sci. Adv. 5, eaax9444 (2019).

43. Morton, O., Bousfield, C.G., Dégny Valé, P. et al. Mining triggers extensive additional deforestation in sub-Saharan Africa. Nature (2026).

44. T. Loft, N. Stevens, F. M. P. Gonçalves, I. Oliveras Menor, Extensive woody encroachment altering Angolan miombo woodlands despite cropland expansion and frequent fires. Glob. Change Biol. 30, e17171 (2024).

45. IUCN, The IUCN Red List of Threatened Species, version 2025-2 (2025); available at https://www.iucnredlist.org.

46. E. Milot, A. Béchet, V. Maris, The dimensions of evolutionary potential in biological conservation. Evol. Appl. 13, 1363–1379 (2020).

47. G. Kier, H. Kreft, T. M. Lee, W. Jetz, P. L. Ibisch, C. Nowicki, J. Mutke, W. Barthlott, A global assessment of endemism and species richness across island and mainland regions. Proc. Natl. Acad. Sci. U.S.A. 106, 9322–9327 (2009).

48. B. D. Mishler, N. Knerr, C. E. González-Orozco, A. H. Thornhill, S. W. Laffan, J. T. Miller, Phylogenetic measures of biodiversity and neo- and paleo-endemism in Australian *Acacia*. Nat. Commun. 5, 4473 (2014).

49. L. M. Lu, L. Mao, T. Yang, J. F. Ye, B. Liu, H. L. Li, M. Sun, J. T. Miller, S. Mathews, H. H. Hu, Y. T. Niu, D. X. Peng, Y. H. Chen, M. Chen, K. L. Xiang, C. T. Le, V. C. Dang, A.M. Lu, P. S. Soltis, D. E. Soltis, J. H. Li, Z. D. Chen, Evolutionary history of the angiosperm flora of China. Nature 554, 234–238 (2018).

50. W. O. Omollo, R. N. Rabarijaona, R. M. Ranaivoson, R. Mijoro, L. B. Russel, Q. Zhang, C. T. Le, J. F. Ye, A. Antonelli, B. Liu, L. M. Lu, Z. D. Chen, Spatial heterogeneity of neo- and paleo-endemism for plants in Madagascar. Curr. Biol. 34, 1271–1283 (2024).

51. D. Rosauer, S. W. Laffan, M. D. Crisp, S. C. Donnellan, L. G. Cook, Phylogenetic endemism: a new approach for identifying geographical concentrations of evolutionary history. Mol. Ecol. 18, 4061–4072 (2009).

52. L. P. Dagallier, S. B. Janssens, G. Dauby, A. Blach-Overgaard, B. A. Mackinder, V. Droissart, J. C. Svenning, Cradles and museums of generic plant diversity across tropical Africa. New Phytol. 225, 2196–2213 (2020).

53. Feng, YL., Hu, HH., Liu, B. et al. A comprehensive dated phylogeny of China’s vascula plants reveals a hidden global biodiversity hotspot. Nat Ecol Evol 10, 794–806 (2026).

54. R. A. Folk, R. L. Stubbs, M. E. Mort, N. Cellinese, J. M. Allen, P. S. Soltis, D. E. Soltis, R. P. Guralnick, Rates of niche and phenotype evolution lag behind diversification in a temperate radiation. Proc. Natl. Acad. Sci. U.S.A, 116, 10874–10882 (2019).

55. M. Sun, F. A. Folk, M. A. Gitzendanner, P. S. Soltis, Z. D. Chen, D. E. Soltis, R. P. Guralnick, Recent, accelerated diversification in rosids occurred outside the tropics. Nat. Commun. 11, 3333 (2020a).

56. F. Forest, R. Grenyer, M. Rouget, T. J. Davies, R. M. Cowling, D. P. Faith, A. Balmford, J. C. Manning, Ş. Procheş, M. Van Der Bank, G. Reeves, Preserving the evolutionary potential of floras in biodiversity hotspots. Nature 445, 757–760 (2007).

57. M. S. Sosef, G. Dauby, A. Blach-Overgaard, X. van der Burgt, L. Catarino, T. Damen, V. Deblauwe, S. Dessein, J. Dransfield, V. Droissart, Exploring the floristic diversity of tropical Africa. BMC Biol. 15, 1–23 (2017).

58. H. Qian, M. Kessler, J. Zhang, Y. Jin, D. E. Soltis, S. Qian, Y. Zhou, P. S. Soltis, Angiosperm phylogenetic diversity is lower in Africa than South America. Sci. Adv. 9, eadj1022 (2023).

59. APG IV, An update of the Angiosperm Phylogeny Group classification for the orders and families of flowering plants. Bot. J. Linn. Soc. 181, 1–20 (2016).

60. Y. Yang, D. K. Ferguson, B. Liu, K. S. Mao, L. M. Gao, S. Z. Zhang, T. Wan, K. Rushforth, Z. X. Zhang, Recent advances on phylogenomics of gymnosperms and a new classification. Plant Divers. 44, 340–350 (2022).

61. PPG I, A community-derived classification for extant lycophytes and ferns. J. Syst. Evol. 54, 563–603 (2016).

62. S. Magallón, S. Gómez-Acevedo, L. L. Sánchez-Reyes, et al., A metacalibrated time-tree documents the early rise of flowering plant phylogenetic diversity. New Phytol. 207, 437–453 (2015).

63. S. A. Smith, J. W. Brown, Constructing a broadly inclusive seed plant phylogeny. Am. J. Bot. 105, 302–314 (2018).

64. S. Ramírez-Barahona, H. Sauquet, S. Magallón, The delayed and geographically heterogeneous diversification of flowering plant families. *Nat*. Ecol. Evol. 4, 1232–1238 (2020).

65. A. R. Zuntini, T. Carruthers, O. Maurin, P. C. Bailey, K. Leempoel, G. E. Brewer, N. Epitawalage, E. Françoso, B. Gallego-Paramo, C. McGinnie, R. Negrão, Phylogenomics and the rise of the angiosperms. Nature 629, 843–850 (2024).

66. Linder HP. 2001. Plant diversity and endemism in sub-Saharan tropical Africa. Journal of Biogeography 28: 169–182.

67. Küper W, Sommer JH, Lovett JC, Mutke J, Linder HP, Beentje HJ, Van Rompaey RSAR, Chatelain C, Sosef M, Barthlott W. 2004. Africa’s hotspots of biodiversity redefined. Annals of the Missouri Botanical Garden 91: 525–535.

68. Plumptre AJ, Davenport TRB, Behangana M, Kityo R, Eilu G, Ssegawa P, Ewango C, Meirte D, Kahindo C, Herremans M et al. 2007. The biodiversity of the Albertine Rift. Biological Conservation 134: 178–194.

69. G. Bañares-de-Dios, M. J. Macía, G. M. de Carvalho, G. Arellano, L. Cayuela, Soil and climate drive floristic composition in tropical forests: A literature review. Front. Ecol. Evol. 10, 866905 (2022).

70. C. M. D. Cramer, G. A. Verboom, Quantitative evaluation of the drivers of species richness in a Mediterranean ecosystem (Cape, South Africa). Ann. Bot. 133, 801–817 (2024).

71. H. P. Linder, The radiation of the Cape flora, southern Africa. Biol. Rev. 78, 597–638 (2003).

72. M. D. Pirie, E. G. H. Oliver, A. Mugrabi de Kuppler, B. Gehrke, N. C. Le Maitre, M. Kandziora, D. U. Bellstedt, The biodiversity hotspot as evolutionary hot-bed: spectacular radiation of Erica in the Cape Floristic Region. BMC Evol. Biol. 16, 190 (2016).

73. B. Bytebier, A. Antonelli, D. U. Bellstedt, H. P. Linder, Estimating the age of fire in the Cape flora of South Africa from an orchid phylogeny. Proc. R. Soc. B 278, 188–195 (2011).

74. J. Murienne, L. R. Benavides, L. Prendini, G. Hormiga, G. Giribet, Forest refugia in Western and Central Africa as ‘museums’ of Mesozoic biodiversity. Biol. Lett. 9, 20120932 (2013).

75. Fjeldså J, Lovett JC. 1997. Geographical patterns of old and young species in African forest biota: the significance of specific montane areas as evolutionary centres. Biodiversity & Conservation 6: 325–346.

76. J. C. Brito, R. Godinho, F. Martínez-Freiría, J. M. Pleguezuelos, H. Rebelo, X. Santos, C. G. Vale, G. Velo-Antón, Z. Boratyński, S. B. Carvalho, S. Ferreira, D. V. Gonçalves, T. L. Silva, P. Tarroso, J. C. Campos, J. V. Leite, J. Nogueira, F. Álvares, N. Sillero, A. Sow, S. Fahd, P.-A. Crochet, S. Carranza. Unravelling biodiversity, evolution and threats to conservation in the Sahara–Sahel. Biol. Rev. 89, 215–231 (2014).

77. . J. Wiens, M. J. Donoghue, The origins of the latitudinal diversity gradient: Revisiting the tropical conservatism hypothesis. J. Biogeogr. 52, e15172 (2025).

78. G. A. Verboom, J. K. Archibald, F. T. Bakker, D. U. Bellstedt, F. Conrad, L. L. Dreyer, F. Forest, C. Galley, P. Goldblatt, J. F. Henning, K. Mummenhoff, Origin and diversification of the Greater Cape flora: ancient species repository, hot-bed of recent radiation, or both? Mol. Phylogenet. Evol. 51, 44–53 (2009).

79. C. O. Webb, D. D. Ackerly, M. A. McPeek, M. J. Donoghue, Phylogenies and community ecology. Annu. Rev. Ecol. Evol. Syst. 33, 475–505 (2002).

80. W. L. Eiserhardt, L. E. S. F. Hansen, T. L. P. Couvreur, J. Dransfield, P. De Lima Ferreira, M. Rakotoarinivo, S. Bellot, W. J. Baker, Explaining extreme differences in species richness among co-occurring palm clades in Madagascar. J. Linn. Soc. Evol. Biol. 3, 1 (2024).

81. W. Wang, K. J. Wurdack, and C. C. Davis, Rosid radiation and the rapid rise of angiosperm-dominated forests. Proc. Natl. Acad. Sci. USA 106, 3857–3862 (2009).

82. Eiserhardt, W.L., Couvreur, T.L.P. & Baker, W.J. (2017). Plant phylogeny as a window on the evolution of hyperdiversity in the tropical rainforest biome. New Phytologist 214: 1408– 1422.

83. A. Antonelli, W. D. Kissling, S. G. Flantua, M. A. Bermúdez, A. Mulch, A. N. Muellner-Riehl, J. Fjeldså, Geological and climatic influences on mountain biodiversity. Nat. Geosci. 11, 718–725 (2018).

84. A. B. Leslie, J. Beaulieu, G. Holman, C. S. Campbell, W. Mei, L. R. Raubeson, S. Mathews, An overview of extant conifer evolution from the perspective of the fossil record. Am. J. Bot. 105, 1531–1544 (2018).

85. D. L. Rabosky, Extinction rates should not be estimated from molecular phylogenies. Evolution 64, 1816–1824 (2010).

86. D. L. Rabosky, Challenges in the estimation of extinction from molecular phylogenies: A response to Beaulieu and O’Meara. Evolution 70, 218–228 (2016).

87. M. R. May, S. Höhna, B. R. Moore, A Bayesian approach for detecting the impact of mass-extinction events on molecular phylogenies when rates of lineage diversification may vary. Methods Ecol. Evol. 7, 947–959 (2016).

88. F. L. Condamine, D. Silvestro, E. B. Koppelhus, A. Antonelli, The rise of angiosperms pushed conifers to decline during global cooling. Proc. Natl. Acad. Sci. U.S.A. 117, 28867–28875 (2020).

89. C. D. Bell, D. E. Soltis, P. S. Soltis, The age and diversification of the angiosperms revisited. Am. J. Bot. 97, 1296–1303 (2010).

90. D. Silvestro, C. D. Bacon, W. Ding, Q. Zhang, P. C. Donoghue, A. Antonelli, Y. Xing, Fossil data support a pre-Cretaceous origin of flowering plants. *Nat*. Ecol. Evol. 5, 449–457 (2021).

91. H. T. Li, T. S. Yi, L. M. Gao, P. F. Ma, T. Zhang, J. B. Yang, M. A. Gitzendanner, P. W. Fritsch, J. Cai, Y. Luo, H. Wang, Origin of angiosperms and the puzzle of the Jurassic gap. Nat. Plants 5, 461–470 (2019).

92. M. Coiro, J. A. Doyle, J. Hilton, How deep is the conflict between molecular and fossil evidence on the age of angiosperms? New Phytol. 223, 83–99 (2019).

93. F. L. Condamine, J. Romieu, G. Guinot, Climate cooling and clade competition likely drove the decline of lamniform sharks. Proc. Natl. Acad. Sci. U.S.A. 116, 20584–20590 (2019).

94. H. Schneider, E. Schuettpelz, K. M. Pryer, R. Cranfill, S. Magallón, R. Lupia, Ferns diversified in the shadow of angiosperms. Nature 428, 553–557 (2004).

95. G. Wu, Q. Ye, H. Liu, H. Schneider, M. Sundue, J. Song, H. Wang, Z. Qiu, Shaded habitats drive higher rates of fern diversification. J. Ecol. 113, 1200–1208 (2025).

96. X. Y. Du, J. M. Lu, L. B. Zhang, J. Wen, L. Y. Kuo, C. M. Mynssen, H. Schneider, D. Z. Li, Simultaneous diversification of Polypodiales and angiosperms in the Mesozoic. Cladistics 37, 518–539 (2021).

97. S. Lehtonen, D. Silvestro, D. N. Karger, C. Scotese, H. Tuomisto, M. Kessler, C. Peña, N. Wahlberg, A. Antonelli, Environmentally driven extinction and opportunistic origination explain fern diversification patterns. Sci. Rep. 7, 4831 (2017).

98. B. F. Jacobs, Palaeobotanical studies from tropical Africa: Relevance to the evolution of forest, woodland and savannah biomes. Philos. Trans. R. Soc. B Biol. Sci. 359, 1573–1583 (2004).

99. D. Silvestro, A. Antonelli, N. Salamin, T. B. Quental, The role of clade competition in the diversification of North American canids. Proc. Natl. Acad. Sci. U.S.A 112, 8684–8689 (2015).

100. J. Igea, A. J. Tanentzap, Angiosperm speciation cools down in the tropics. Ecol. Lett. 23, 692–700 (2020).

101. D. Dimitrov, X. Xu, X. Su, N. Shrestha, Y. Liu, J. D. Kennedy, L. Lyu, D. Nogués-Bravo, J. Rosindell, Y. Yang, J. Fjeldså, Diversification of flowering plants in space and time. Nat. Commun. 14, 760 (2023).

102. Barnabas H. Daru et al., Biogeographic processes underlying global patterns of plant diversity. Science392,845–849(2026).

103. Y. Bouchenak-Khelladi, O. Maurin, J. Hurter, M. van der Bank, The evolutionary history and biogeography of Mimosoideae (Leguminosae): an emphasis on African acacias. Mol. Phylogenet. Evol. 57, 495–508 (2010)

104. T. Särkinen, R. T. Pennington, M. Lavin, M. F. Simon, C. E. Hughes, Evolutionary islands in the Andes: persistence and isolation explain high endemism in Andean dry tropical forests. J. Biogeogr. 39, 884–900 (2012).

105. J. L. Silva, A. F. Souza, A. Caliman, E. L. Voigt, J. E. Lichston, Weak whole-plant trait coordination in a seasonally dry South American stressful environment. Ecol. Evol. 8, 4–12 (2018).

106. L. Ségalen, J. A. Lee-Thorp, T. Cerling, Timing of C4 grass expansion across sub-Saharan Africa. J. Hum. Evol. 53, 549–559 (2007).

107. P. J. Polissar, C. Rose, K. T. Uno, S. R. Phelps, P. deMenocal, Synchronous rise of African C4 ecosystems 10 million years ago in the absence of aridification. Nat. Geosci. 12, 657–660 (2019).

108. G. J. Kergoat, F. L. Condamine, E. F. Toussaint, C. Capdevielle-Dulac, A. L. Clamens, J. Barbut, P. Z. Goldstein, B. Le Ru, Opposite macroevolutionary responses to environmental changes in grasses and insects during the Neogene grassland expansion. Nat. Commun. 9, 5089 (2018).

109. M. Arakaki, P. A. Christin, R. Nyffeler, A. Lendel, U. Eggli, R. M. Ogburn, E. J. Edwards, Contemporaneous and recent radiations of the world’s major succulent lineages. Proc. Natl. Acad. Sci. U.S.A. 108, 8379–8384 (2011).

110. S. Buerki, D. S. Devey, M. W. Callmander, P. B. Phillipson, F. Forest, Spatio-temporal history of the endemic genera of Madagascar. Bot. J. Linn. Soc. 171, 304–329 (2013).

111. C. Klak, G. Reeves, T. Hedderson, Unmatched tempo of evolution in Southern African semi-desert ice plants. Nature 427, 63–65 (2004).

112. L. M. Valente, A. W. Britton, M. P. Powell, A. S. Papadopulos, P. M. Burgoyne, V. Savolainen, Correlates of hyperdiversity in southern African ice plants (Aizoaceae). Bot. J. Linn. Soc. 174, 110–129 (2014).

113. S. A. Smith, M. J. Donoghue, Rates of molecular evolution are linked to life history in flowering plants. Science, 322, 86–89 (2008).

114. F. C. Boucher, G. A. Verboom, S. Musker, A. G. Ellis, Plant size: a key determinant of diversification? New Phytol. 216, 24–31 (2017).

115. A. Rice, P. Šmarda, M. Novosolov, M. Drori, L. Glick, N. Sabath, S. Meiri, J. Belmaker, I. Mayrose, The global biogeography of polyploid plants. Nat. Ecol. Evol. 3, 265–273 (2019).

116. A. G. Ellis, A. E. Weis, S. G. Brandon, Evolutionary radiation of ‘stone plants’ in the genus *Argyroderma* (Aizoaceae): unraveling the effects of landscape, habitat, and flowering time. Evolution 60, 39–55 (2006).

117. Weil, S.-S., S. Lavergne, F. C. Boucher, W. L. Allen, and L. Gallien. 2025. “Can Macroevolution Inform Contemporary Extinction Risk?.” Ecology Letters28, no. 7: e70171

118. H. Morlon, E. Lewitus, F. L. Condamine, M. Manceau, J. Clavel, J. Drury, RPANDA: an R package for macroevolutionary analyses on phylogenetic trees. Methods Ecol. Evol. 7, 589– 597 (2016).

119. E. B. Knox, J. D. Palmer, Chloroplast DNA variation and the recent radiation of the giant senecios (Asteraceae) on the tall mountains of eastern Africa. Proc. Natl. Acad. Sci. U.S.A. 92, 10349–10353 (1995).

120. A. Assefa, D. Ehrich, P. Taberlet, S. Nemomissa, C. Brochmann, Pleistocene colonization of afro-alpine “sky islands” by the arctic–alpine *Arabis alpina*. Heredity 99, 133–142 (2007).

121. B. Gehrke, C. Brauchler, K. Romoleroux, M. Lundberg, G. Heubl, T. Eriksson, Molecular phylogenetics of *Alchemilla*, *Aphanes* and *Lachemilla* (Rosaceae) inferred from plastid and nuclear intron and spacer DNA sequences, with comments on generic classification. Mol. Phylogenet. Evol. 47, 1030–1044 (2008).

122. I. Sanmartín, C. L. Anderson, M. Alarcón, F. Ronquist, J. J. Aldasoro, Bayesian island biogeography in a continental setting: the Rand Flora case. Biol. Lett. 6, 703–707 (2010).

123. S. M. Durant, N. Pettorelli, S. Bashir, R. Woodroffe, T. Wacher, P. De Ornellas, C. Ransom, T. Abáigar, M. Abdelgadir, H. El Alqamy, M. Beddiaf, Forgotten biodiversity in desert ecosystems. Science 336, 1379–1380 (2012).

124. H. Ralimanana, A. L. Perrigo, R. J. Smith, J. S. Borrell, S. Faurby, M. T. Rajaonah, T. Randriamboavonjy, M. S. Vorontsova, R. S. Cooke, L. N. Phelps, F. Sayol, Madagascar’s extraordinary biodiversity: threats and opportunities. Science 378, eadf1466 (2022).

125. M. Sun, R. A. Folk, M. A. Gitzendanner, P. S. Soltis, Z.-D. Chen, D. E. Soltis, R. P. Guralnick, Estimating rates and patterns of diversification with incomplete sampling: a case study in the rosids. Am. J. Bot. 107, 895–909 (2020).

126. M. J. Brown, B. E. Walker, N. Black, R. H. Govaerts, I. Ondo, R. Turner, E. Nic Lughadha, rWCVP: a companion R package for the World Checklist of Vascular Plants. New Phytol. 240, 1355–1365 (2023).

127. A. Zizka, D. Silvestro, T. Andermann, J. Azevedo, C. Duarte Ritter, D. Edler, H. Farooq, A. Herdean, M. Ariza, R. Scharn, CoordinateCleaner: standardized cleaning of occurrence records from biological collection databases. Methods Ecol. Evol. 10, 744–751 (2019).

128. A. E. Zanne, D. C. Tank, W. K. Cornwell, J. M. Eastman, S. A. Smith, R. G. FitzJohn, D. J. McGlinn, B. C. O’Meara, A. T. Moles, P. B. Reich, D. L. Royer, Three keys to the radiation of angiosperms into freezing environments. Nature 506, 89–92 (2014).

129. J. Kattge, S. Díaz, S. Lavorel, I. C. Prentice, P. Leadley, G. Bönisch, E. Garnier, M. Westoby, P. B. Reich, I. J. Wright, J. H. C. Cornelissen, TRY – a global database of plant traits. Glob. Change Biol. 17, 2905–2935 (2013).

130. J. Allaire, RStudio: Integrated Development Environment for R (Boston, MA, 2012).

131. D. J. Winter, rentrez: an R package for the NCBI eUtils API. R J. 9, 520–526 (2017).

132. K. Katoh, D. M. Standley, MAFFT multiple sequence alignment software version 7: improvements in performance and usability. Mol. Biol. Evol. 30, 772–780 (2013).

133. T. A. Hall, BioEdit: a user-friendly biological sequence alignment editor and analysis program for Windows 95/98/NT. Nucleic Acids Symp. Ser. 41, 95–98 (1999).

134. A. Stamatakis, RAxML version 8: a tool for phylogenetic analysis and post-analysis of large phylogenies. Bioinformatics 30, 1312–1313 (2014).

135. A. Rambaut, A. Drummond, FigTree, version 1.3.1 [computer program] (2009).

136. S. A. Smith, B. C. O’Meara, treePL: divergence time estimation using penalized likelihood for large phylogenies. Bioinformatics 28, 2689–2690 (2012).

137. Laffan, S. W., Lubarsky, E., & Rosauer, D. F. (2010). Biodiverse: a tool for the spatial analysis of biological and related diversity. Ecography, 33(4), 643–647.

138. Faith, D. P. (1992). Conservation evaluation and phylogenetic diversity. Biological Conservation, 61(1), 1–10.

139. Kembel, S. W., Cowan, P. D., Helmus, M. R., Cornwell, W. K., Morlon, H., Ackerly, D. D., Blomberg, S. P., & Webb, C. O. (2010). Picante: R tools for integrating phylogenies and ecology. Bioinformatics, 26(11), 1463–1464.

140. F. L. Condamine, J. Rolland, S. Höhna, F. A. H. Sperling, I. Sanmartín, Testing the role of the Red Queen and Court Jester as drivers of the macroevolution of Apollo butterflies. Syst. Biol. 67, 940–964 (2018).

141. D. L. Rabosky, M. Grundler, C. Anderson, P. Title, J. J. Shi, J. W. Brown, H. Huang, J. G. Larson, BAMM tools: an R package for the analysis of evolutionary dynamics on phylogenetic trees. Methods Ecol. Evol. 5, 701–707 (2014).

142. M. Plummer, N. Best, K. Cowles, K. Vines, CODA: convergence diagnosis and output analysis for MCMC. R News 6, 7–11 (2006).

143. W. Jetz, G. H. Thomas, J. B. Joy, K. Hartmann, A. O. Mooers, The global diversity of birds in space and time. Nature 491, 444–448 (2012).

144. R. Barnes, dggridR: discrete global grids for R, R package version 0.1.12 (2017). 10.32614/CRAN.package.dggridR

145. H. Morlon, S. Robin, F. Hartig, Studying speciation and extinction dynamics from phylogenies: addressing identifiability issues. Trends Ecol. Evol. 37, 497–506 (2022).

146. O. Maliet, F. Hartig, H. Morlon, A model with many small shifts for estimating species-specific diversification rates. Nat. Ecol. Evol. 3, 1086–1092 (2019).

147. O. Maliet, H. Morlon, Fast and accurate estimation of species-specific diversification rates using data augmentation. Syst. Biol. 71, 353–366 (2022).

148. S. Höhna, W. A. Freyman, Z. Nolen, J. P. Huelsenbeck, M. R. May, B. R. Moore, A Bayesian approach for estimating branch-specific speciation and extinction rates. bioRxiv 555805 (2019).

149. A. Rambaut, A. J. Drummond, D. Xie, G. Baele, M. A. Suchard, Posterior summarization in Bayesian phylogenetics using Tracer 1.7. Syst. Biol. 67, 901–904 (2018).

150. D. N. Karger, N. E. Zimmermann, Climatologies at high resolution for the Earth’s land surface areas: CHELSA V1.2 technical specification (2021); available online.

151. Esri. (2020). ArcGIS Desktop: Release 10.8. Environmental Systems Research Institute, Redlands, CA. https://www.esri.com/en-us/arcgis/products/arcgis-desktop/resources.

152. A. D. Barnosky, Distinguishing the effects of the Red Queen and Court Jester on Miocene mammal evolution in the Northern Rocky Mountains. J. Vertebr. Paleontol. 21, 172–185 (2001).

153. C. R. Scotese, N. Wright, PALEOMAP Paleodigital Elevation Models (PaleoDEMS) for the Phanerozoic. PALEOMAP Project (2018).

154. S. Höhna, M. R. May, B. R. Moore, Phylogeny simulation and diversification rate analysis with TESS. (2015).

155. R. E. Kass, A. E. Raftery, Bayes Factors. J. Am. Stat. Assoc. 90, 773–795 (1995).

156. S. Höhna, B. T. Kopperud, A. F. Magee, CRABS: Congruent rate analyses in birth–death scenarios. Methods Ecol. Evol. 13, 2709–2718 (2022).

157. W. P. Maddison, P. E. Midford, S. P. Otto, Estimating a binary character’s effect on speciation and extinction. Syst. Biol. 56, 701–710 (2007).

158. R. G. FitzJohn, Diversitree: Comparative phylogenetic analyses of diversification in R. Methods Ecol. Evol. 3, 1084–1092 (2012).

159. B. H. Daru, M. van der Bank, T. J. Davies, Spatial incongruence among hotspots and complementary areas of tree diversity in southern Africa. Divers. Distrib. 21, 769–780 (2015).

160. M. Tietje, A. Antonelli, F. Forest, R. Govaerts, S. A. Smith, M. Sun, W. J. Baker, W. L. Eiserhardt, Global hotspots of plant phylogenetic diversity. New Phytol. 240, 1636–1646 (2023)

161. C. E González-Orozco, & M. Parra-Quijano Comparing species and evolutionary diversity metrics to inform conservation. Divers. Distrib. 29, 224–231. (2023).

