## Supplementary figures and Tables for "The diversity, evolution, and conservation of African plants"

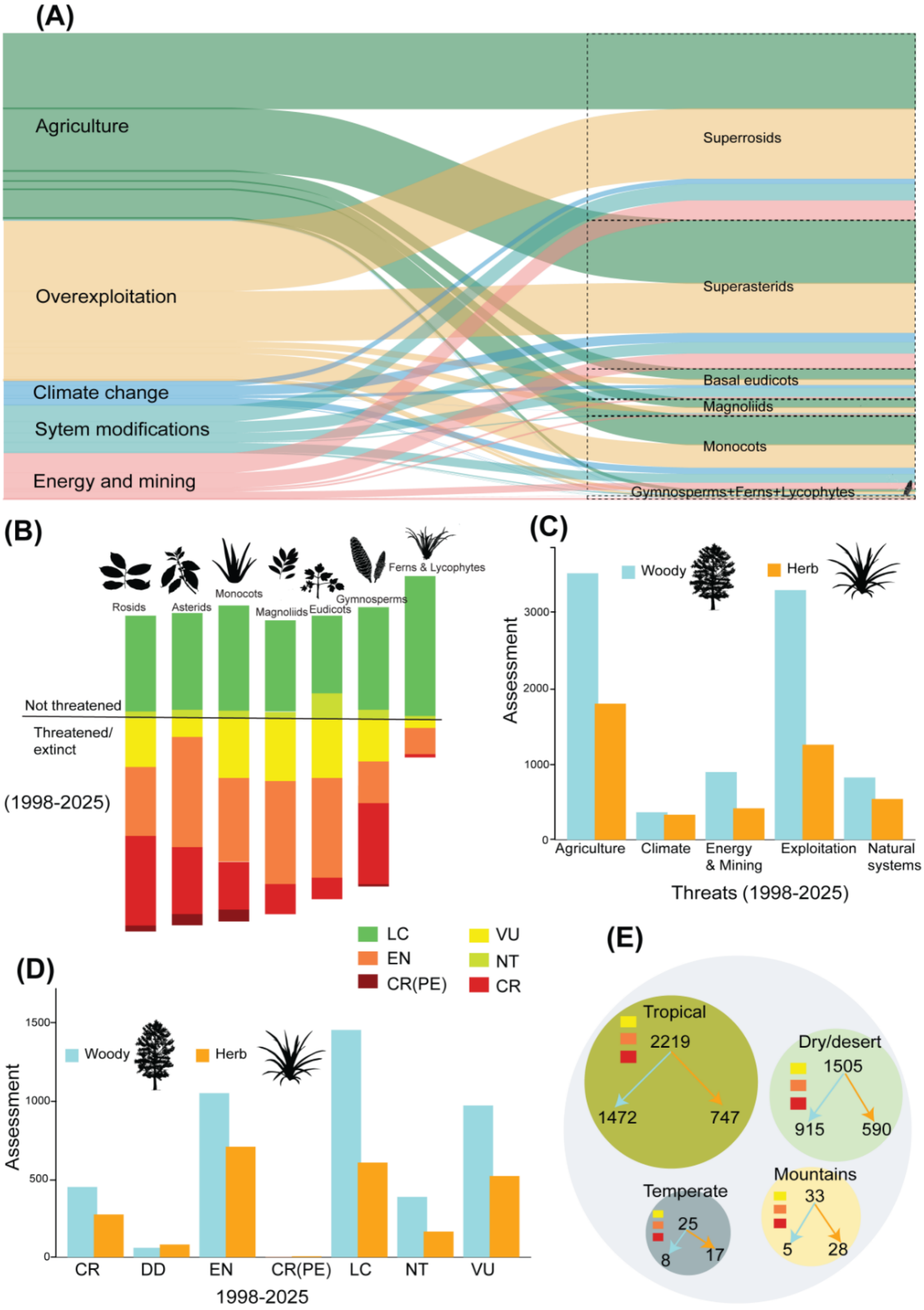


**Fig. S2.** (A) **Threats and conservation status of African vascular plants**. Alluvial plot showing threats defined by the IUCN and their associations with major groups of vascular plants (angiosperms, gymnosperms, ferns, and lycophytes). The widths of boxes and connecting flows reflect the number of species affected by each threat. The color scheme is consistent across panels. Some threat classes were renamed for brevity and clarity; for example, the IUCN category "biological resource use" is referred to as "overexploitation" here and in the main text, following the terminology of the Intergovernmental Science-Policy Platform on Biodiversity and Ecosystem Services (IPBES). (**B**) IUCN Red List assessment categories for major plant groups in Africa. Assessment categories and colors follow IUCN Red List standards. Category distributions (shown using bars) include magnoliids (N = 252), gymnosperms (N = 86), rosids (N = 2,598), monocots (N = 1,074), asterids (N = 2,111), eudicots (N = 366), and ferns and lycophytes (N = 46). (**C–D**) Distribution of threats by life form (C) and number of species per IUCN Red List category in each life form (D). (**E**) Number of Critically Endangered (CR), Endangered (EN), and Vulnerable (VU) species across habitat types. Arrows within each habitat indicate the number of woody and herbaceous species, with color coding consistent with panels (C) and (D). Abbreviations: LC = Least Concern; NT = Near Threatened; CR (PE) = Critically Endangered (Possibly Extinct).

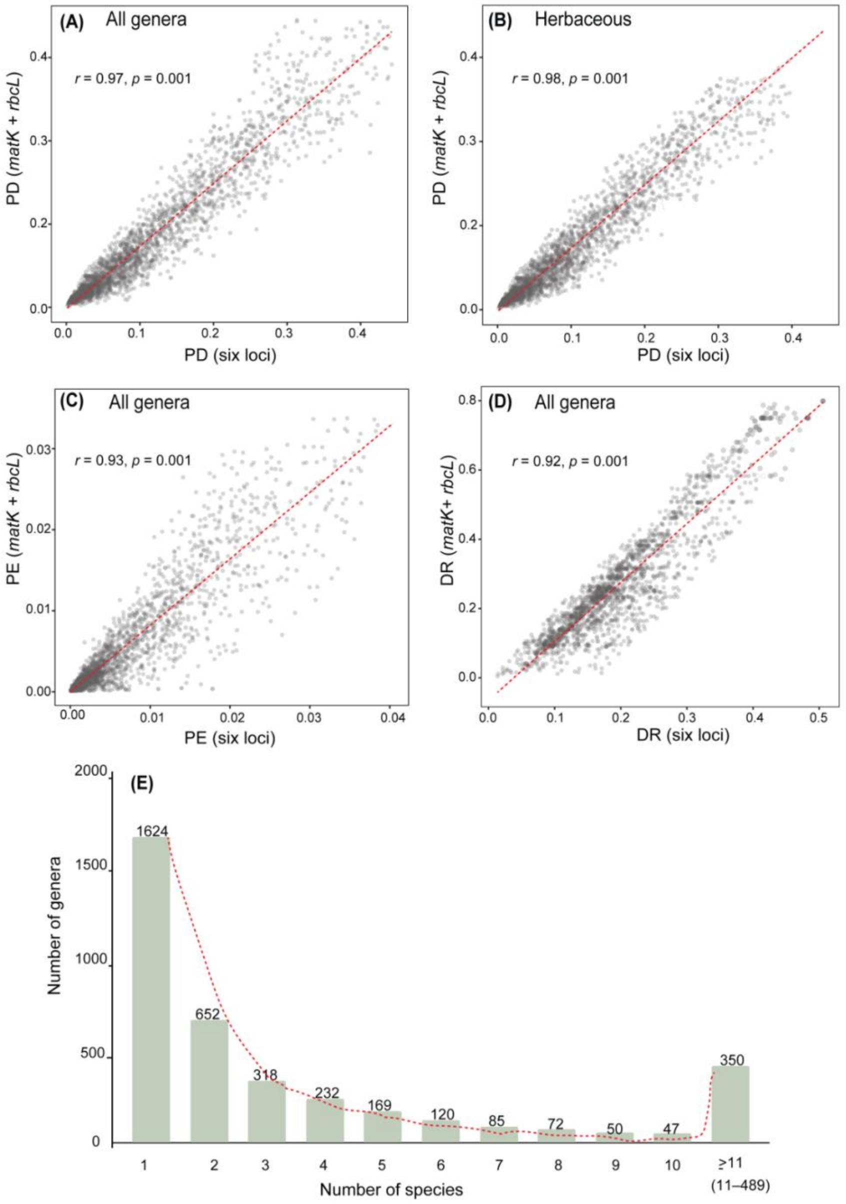


**Fig. S3.** Correlation between patterns of phylogenetic diversity and phylogenetic endemism derived from the two phylogenetic datasets: the *matK* + *rbcL* tree and the six-locus tree (*atpB*, *ndhF*, *matK*, *rbcL*, *trnL-trnF*, and ITS). Panels show phylogenetic diversity for (A) all genera, (B) herbaceous genera, and (C) woody genera, and phylogenetic endemism for (D) all genera. (E) Frequency distribution for the number of congeneric species used in our molecular tree construction. In total, sequences for ca. 44% of genera were from only one species, and ca. 9% of genera had 11 or more species.

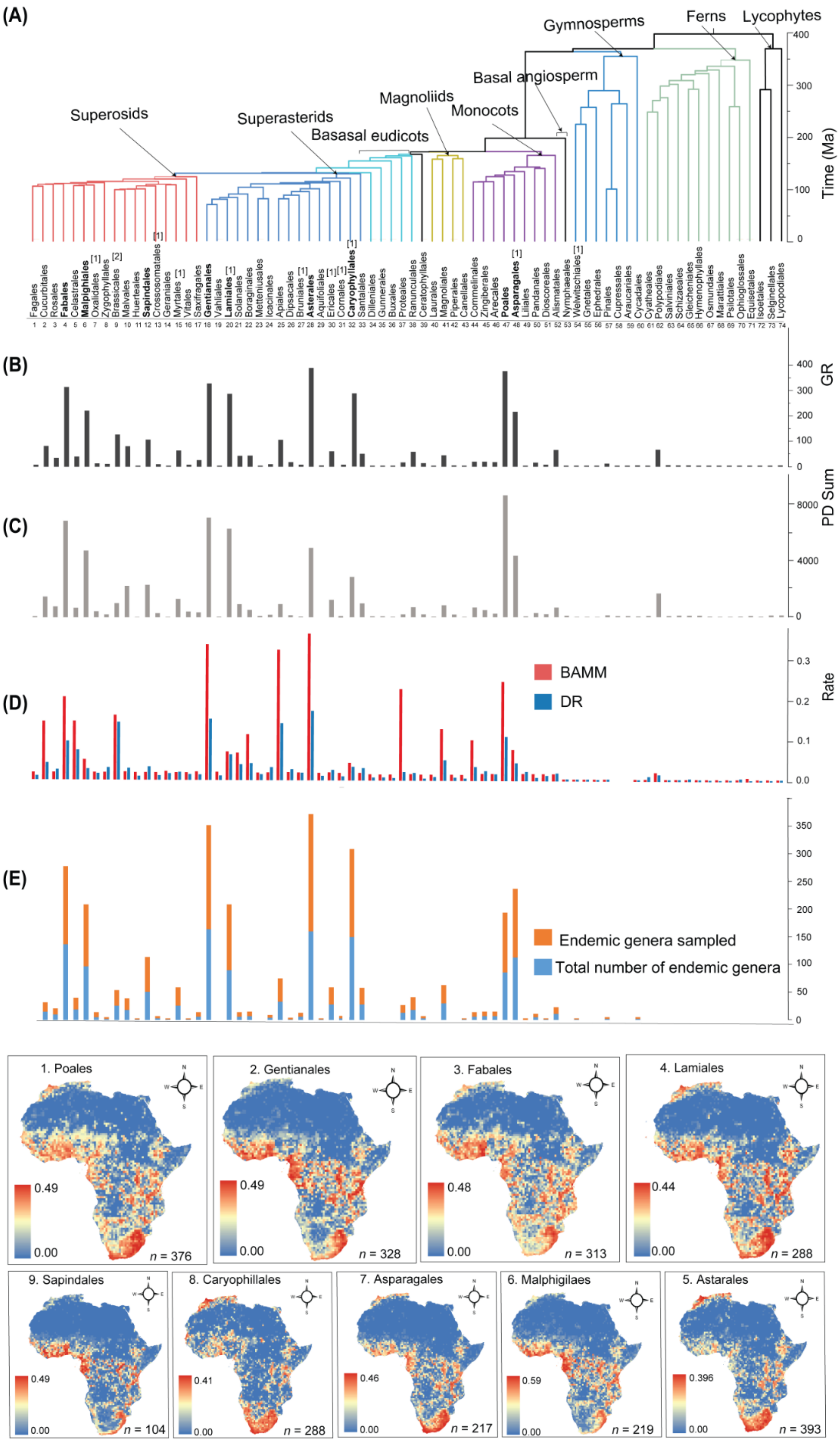

(A)
Gymnosperms
Ferns
Lycophytes
Basal angiosperm
Magnoliids
Monocots
Superosids
Superasterids
Basasal eudicots
Time (Ma)
400
300
200
100
0
Fagales
Cucurbitales
Rosales
Fabales
Celastrales
Malpighiales
Oxalidales
Zygophyllales
Brassicales
Malvales
Huerteales
Sapindales
Crossosomatales
Geraniales
Myrtales
Vitales
Saxifragales
Gentianales
Vahliales
Lamiales
Solanales
Boraginales
Metteniusales
Icacinales
Apiales
Dipsacales
Bruniales
Asterales
Aquifoliales
Ericales
Cornales
Caryophyllales
Santalales
Dilleniales
Gunnerales
Buxales
Proteales
Ranunculales
Ceratophyllales
Laurales
Magnoliales
Piperales
Canellales
Commelinales
Zingiberales
Arecales
Poales
Asparagales
Liliales
Pandanales
Dioscoreales
Alismatales
Nymphaeales
Welwitschiales
Gnetales
Ephedrales
Pinales
Cupressales
Araucariales
Cycadales
Cyatheales
Polypodiales
Salviniales
Schizaeales
Gleicheniales
Hymenophyllales
Osmundales
Marattiales
Psilotales
Ophioglossales
Equisetales
Isoetales
Selginellales
Lycopodiales
(B)
GR
(C)
PD Sum
8000
4000
(D)
BAMM
DR
Rate
0.3
0.2
0.1
0.0
(E)
Endemic genera sampled
Total number of endemic genera
350
300
250
200
150
100
50
1. Poales
0.49
0.00
n = 376
2. Gentianales
0.49
0.00
n = 328
3. Fabales
0.48
0.00
n = 313
4. Lamiales
0.44
0.00
n = 288
9. Sapindales
0.49
0.00
n = 104
8. Caryophillales
0.41
0.00
n = 288
7. Asparagales
0.46
0.00
n = 217
6. Malphigilaes
0.59
0.00
n = 219
5. Astarales
0.396
0.00
n = 393

**Fig. S4.** (A) Ordinal time-tree with the major clades indicated. Top nine orders with high phylogenetic diversity and generic richness are highlighted in bold. Numbers indicated in parentheses indicate number of endemic families, and the numbers shown below each order correspond to Figure S6A. (B-C), Proportion of phylogenetic diversity (grey bars) (B), and generic richness (black bars) (C) for genera in each order. (D) Order-level diversification signals derived from aggregated genus-level diversification rates (median BAMM tip rates, red bars; median DR statistic, blue bars). (E) Sampled endemic genera (orange bars) and total endemic genera (light blue bars) in each order. Maps labelled 1-9 indicate the phylogenetic diversity of the top nine orders (bold) in Africa, with number of genera (n) indicated.

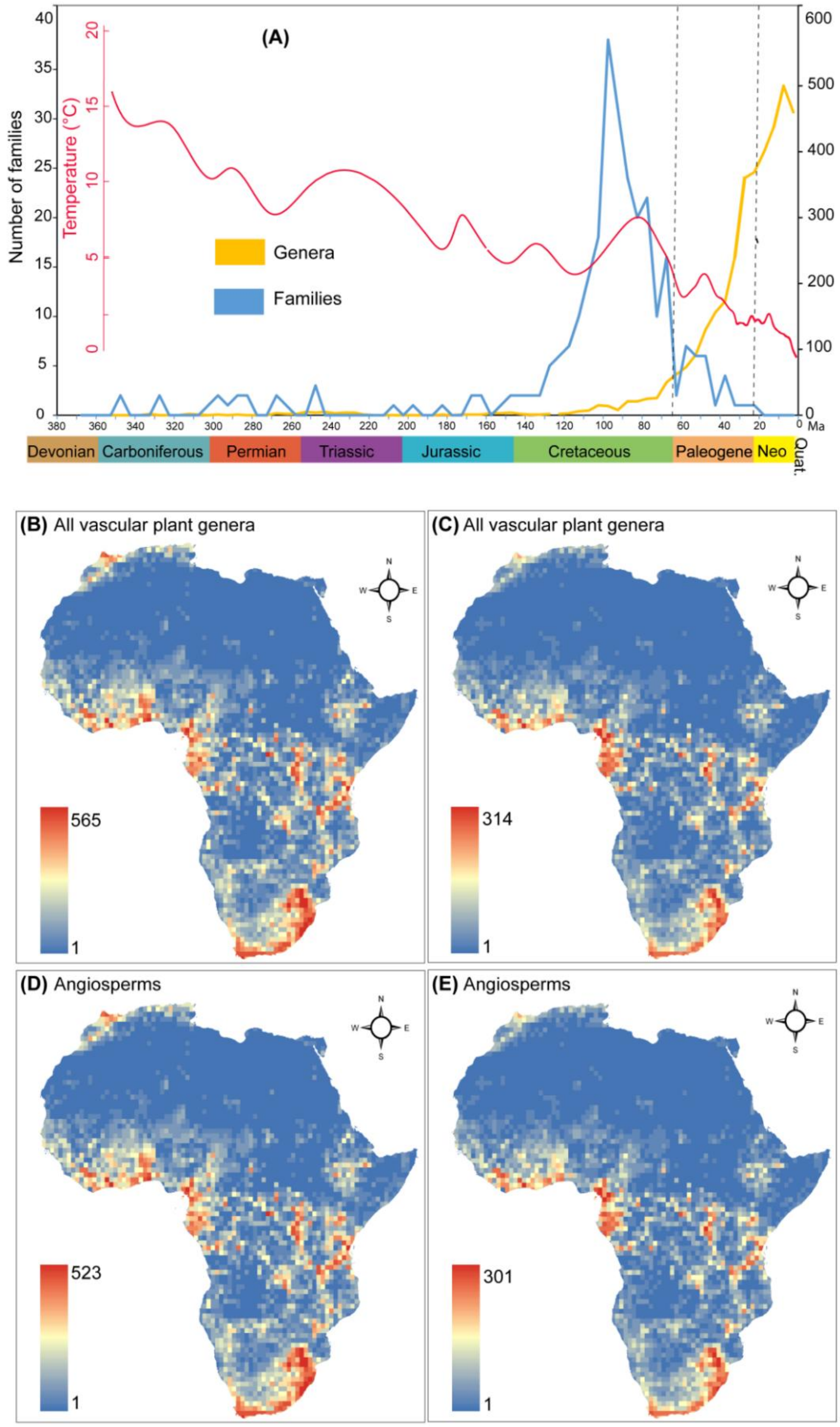


**Fig. S5. Geographic and temporal patterns of lineage origins for vascular plant genera in Africa.** (A) Temporal distribution of lineage origins showing the number of genera and families that originated during each five-million-year interval in Africa. (B–E) Geographic patterns of richness for vascular plant genera that originated before the Miocene (>23 Ma) (B) and after the Miocene (<23 Ma) (C) and (D–E) for angiosperm genera that originated before (>23 Ma) (C) and after (<23 Ma) (D) the Miocene. Color scales represent the number of genera per grid cell.

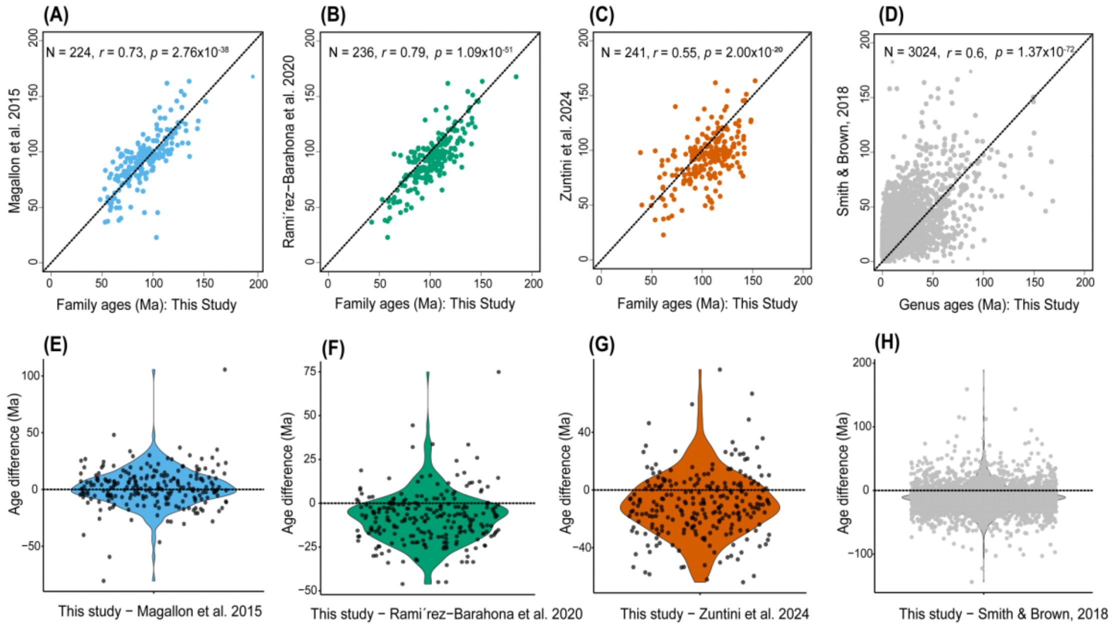


**Fig. S6. Comparisons of node age estimates for vascular plant genera in Africa.** Node ages were estimated using treePL in this study and compared with divergence times reported in recent publications. (A–D) Correlations of node ages for families and genera sampled in this study: (A–C) correlations of family ages between this study and (A) Magallón et al. (2015), (B) Ramírez-Barahona et al. (2020), and (C) Zuntini et al. (2024); (D) correlation of genus ages between this study and Smith and Brown (2018). (E–H) Differences in ages between our study and recent publications for families and genera, respectively.

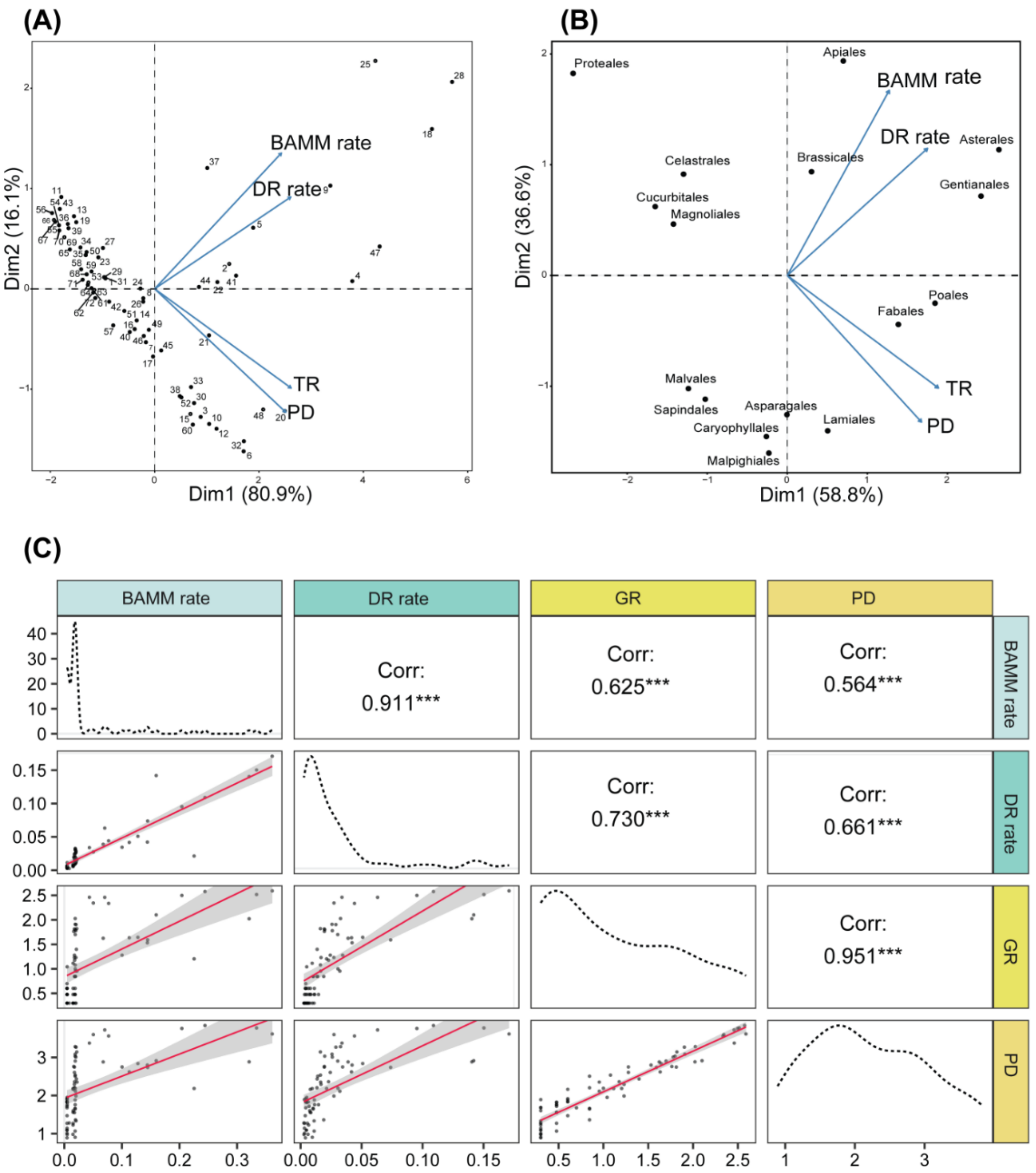


**Fig. S7. Covariation among diversification, phylogenetic, and richness metrics at the order level.** (A) Principal component analysis (PCA) of plant orders illustrating multivariate relationships among phylogenetic diversity (PD), generic richness (GR), and diversification rates estimated from DR statistic and BAMM. Points represent individual orders, with numbers corresponding to those shown below each order in Figure S3A. Arrows indicate the direction and relative contribution of each variable to the ordination. Percent variance explained by each axis is shown. (B) PCA biplot highlighting the positions of major plant orders in relation to diversification and phylogenetic metrics, illustrating contrasts among lineages with high diversification rates, phylogenetic diversity, and richness. (C) Pairwise relationships among BAMM rates, DRstatistic, GR, and PD at the order level. Diagonal panels show variable distributions; off-diagonal panels show scatterplots with fitted linear regressions. Upper panels report Pearson correlation coefficients (Corr.) and significance levels (*** $p < 0.001$).

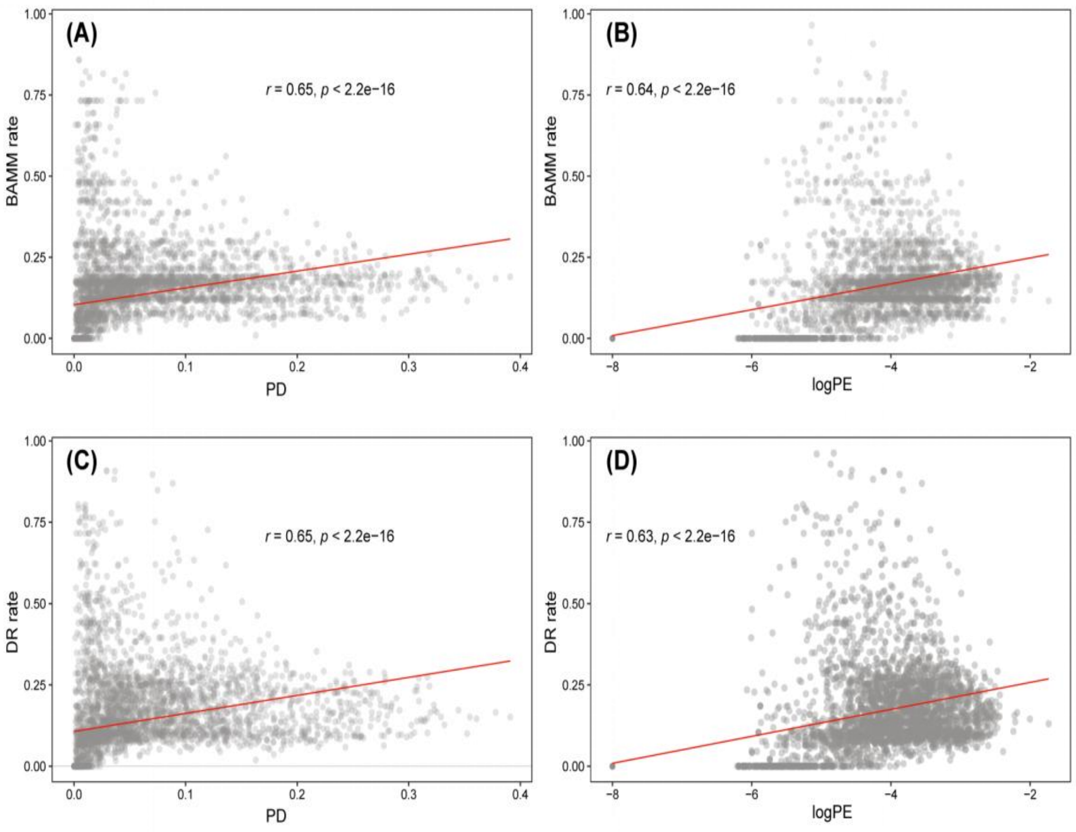


**Fig. S8. Relationships between phylogenetic diversity, phylogenetic endemism, and diversification rates across Africa.** Scatterplots show correlations per grid cell between phylogenetic diversity (PD), phylogenetic endemism (PE), and diversification rate estimates for African vascular plant genera. (A) Relationship between PD and BAMM-estimated diversification rate. (B) Relationship between log-transformed PE and BAMM-estimated diversification rate. (C) Relationship between PD and diversification rate (DR statistic). (D) Relationship between log-transformed PE and DR. Reported values show Pearson correlation coefficients (r) and associated p-values.

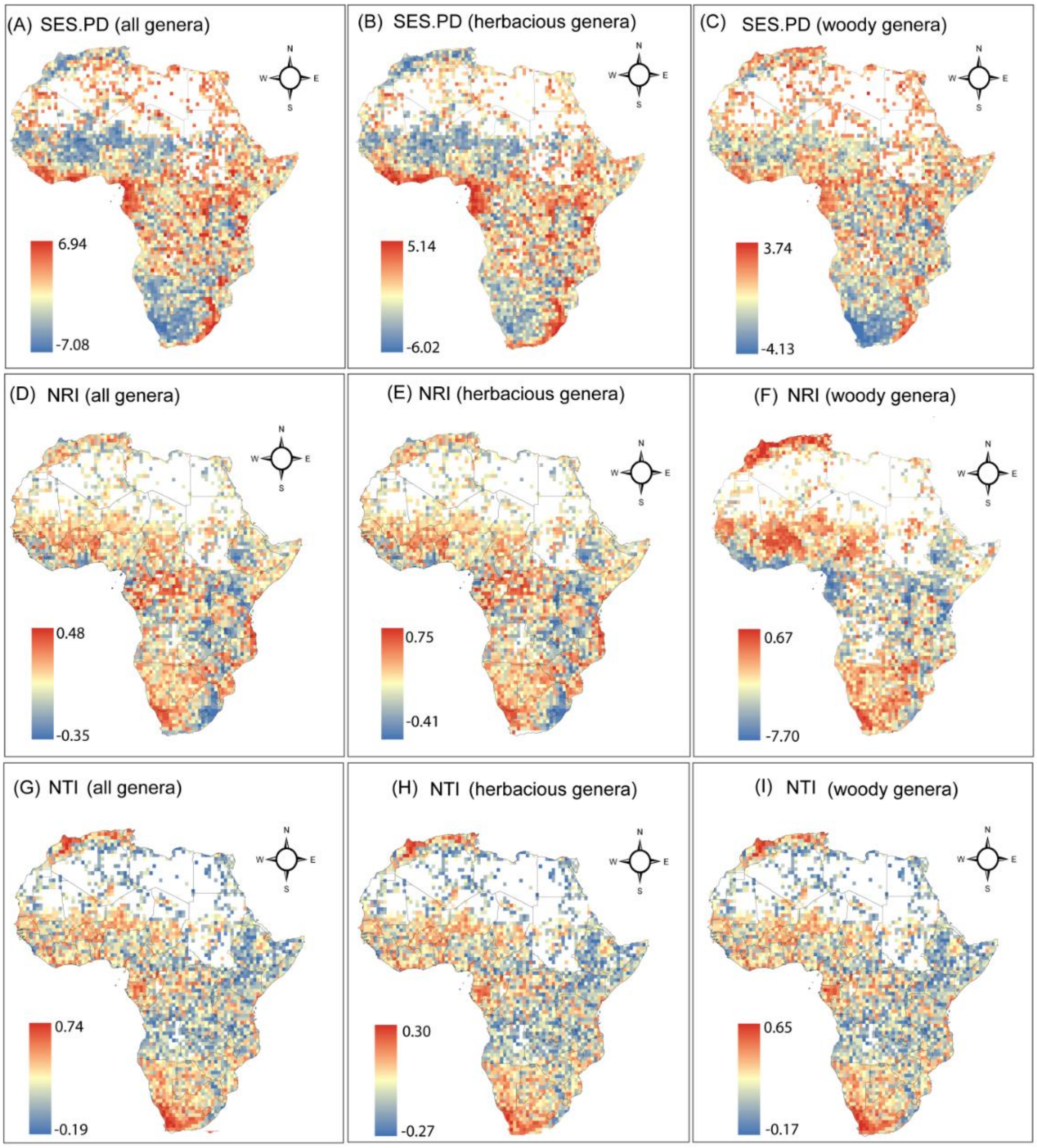


**Fig. S9. Patterns of phylogenetic structure for vascular plant genera in Africa.** (A–C) SES-PD for all genera (A), herbaceous genera (B), and woody genera (C). (D–F) NRI for all genera (D), herbaceous genera (E), and woody genera (F). (G–I) NTI for all genera (G), herbaceous genera (H), and woody genera (I). The analyses include 3,719 vascular plant genera (herbaceous genera, n = 2,101; woody genera, n = 1,325; genera with both woody and herbaceous species, n = 293). Maps are displayed in the WGS84 coordinate system.

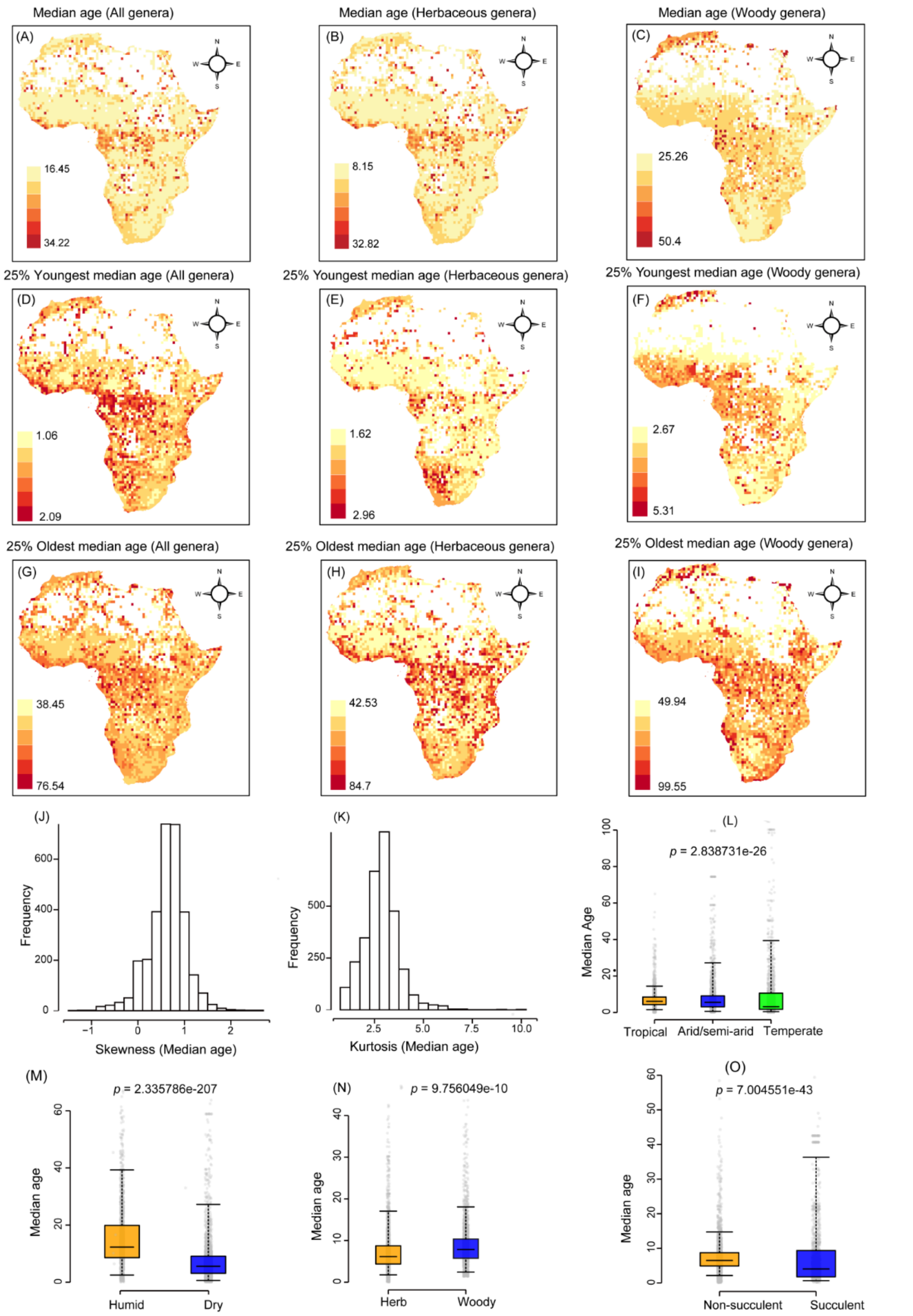


**Fig. S10. Geographic patterns of median ages for vascular plant genera in Africa.** (**A–I**) Median ages for all genera, herbaceous genera, and woody genera (from left to right), based on all sampled genera (A–C), the youngest 25% of genera (D–F), and the oldest 25% of genera (G–I) in each grid cell. Maps are displayed in the WGS84 coordinate system. (**J–K**) Histograms and distributions of (J) skewness and (K) kurtosis of median ages across grid cells for vascular plant genera in Africa. The ranges of skewness and kurtosis (computed as the fourth standardized moment) are shown for median genus ages across all genera. (**L–O**) Boxplots of median genus age by (L) habitat (tropical = orange; non-tropical = blue; temperate = green), (M) aridity (humid = orange; dry = blue), (N) life form (herbaceous = orange; woody = blue), and (O) succulence (non-succulent = orange; succulent = blue). Each boxplot shows the median (solid line), interquartile range (box = 25%–75%), and whiskers (5%–95% intervals). Analyses include 3,719 vascular plant genera (herbaceous genera, n = 2,101; woody genera, n = 1,325; genera with both woody and herbaceous species, n = 293).

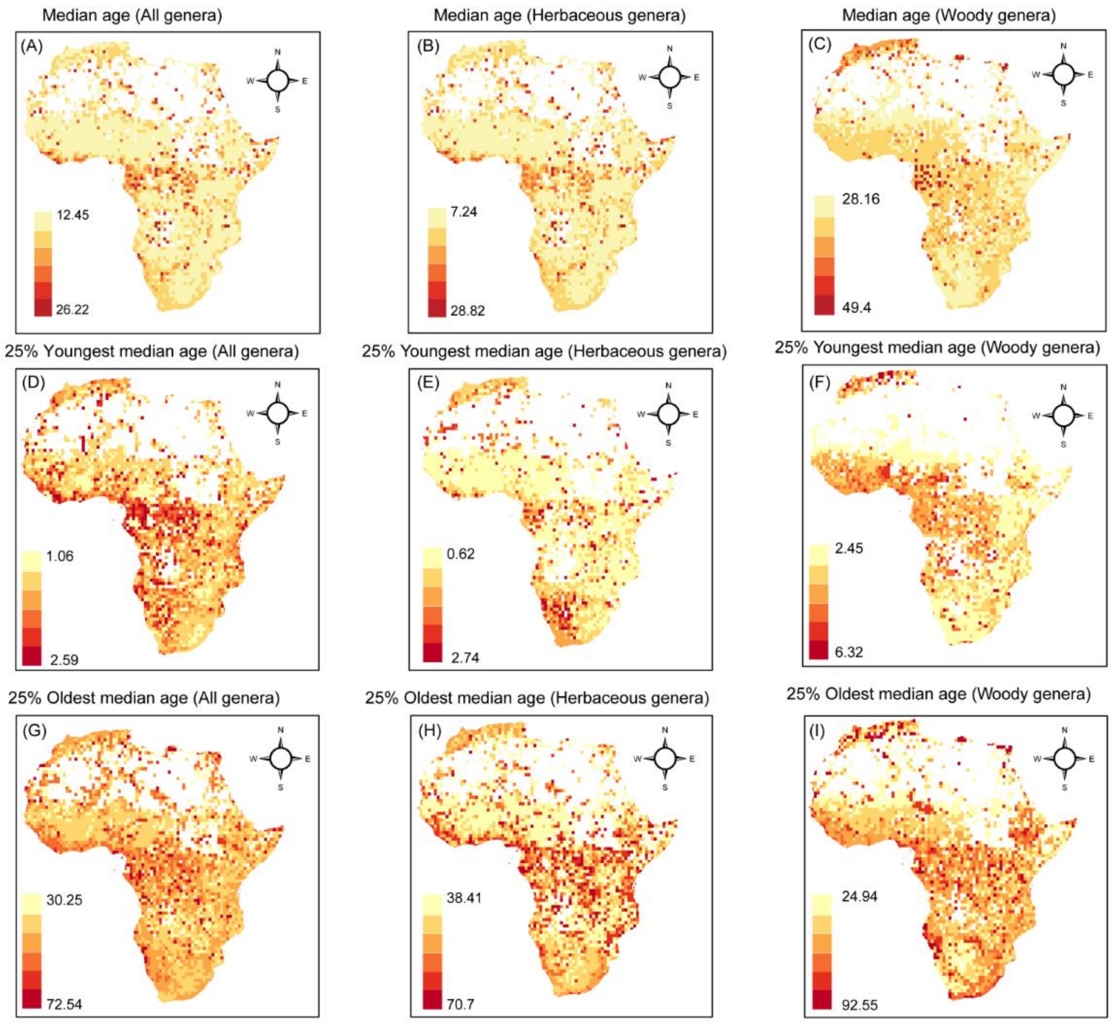


**Fig. S11.** Geographic patterns of median ages for angiosperm plant genera in Africa. (**A–I**) Median ages for all genera, herbaceous genera, and woody genera (from left to right), based on all sampled genera (A–C), the youngest 25% of genera (D–F), and the oldest 25% of genera (G–I) in each grid cell. Maps are displayed in the WGS84 coordinate system. Analyses include 3,599 angiosperm plant genera (herbaceous genera, n = 2,160; woody genera, n = 1,285; genera with both woody and herbaceous species, n = 154).

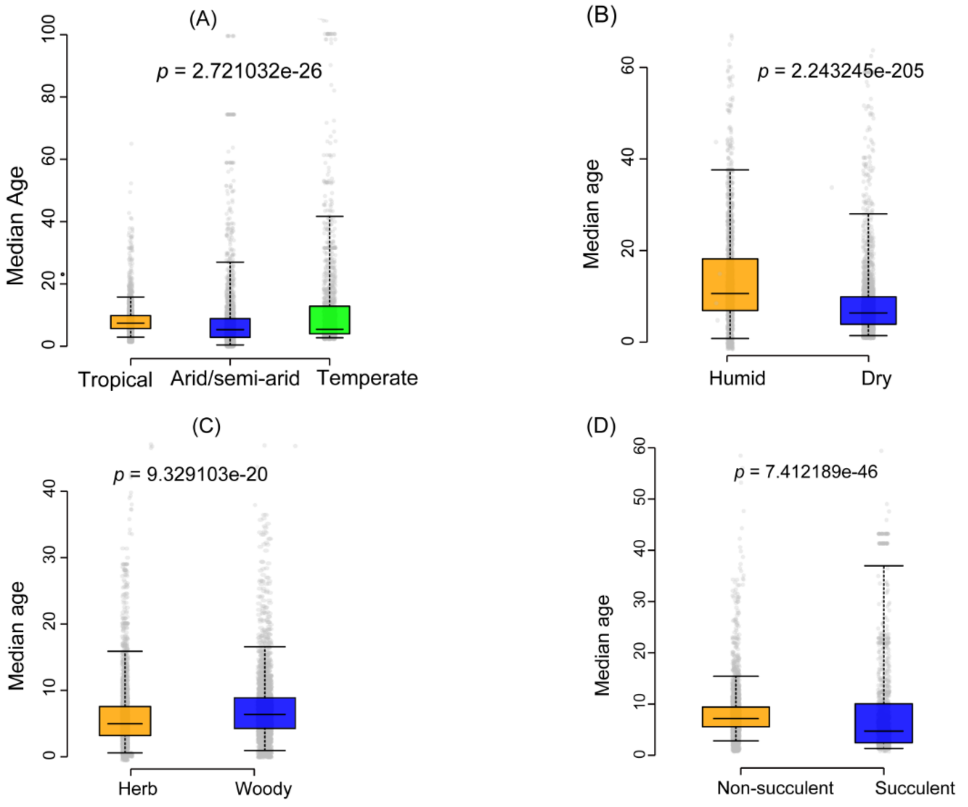


**Fig. S12.** (**A–D**) Boxplots of median genus age by (A) habitat (tropical = orange; non-tropical = blue; temperate = green), (B) aridity (humid = orange; dry = blue), (C) life form (herbaceous = orange; woody = blue), and (D) succulence (non-succulent = orange; succulent = blue). Each boxplot shows the median (solid line), interquartile range (box = 25%–75%), and whiskers (5%–95% intervals). Analyses include 3,599 angiosperm plant genera (herbaceous genera, n = 2,160; woody genera, n = 1,285; genera with both woody and herbaceous species, n = 154).

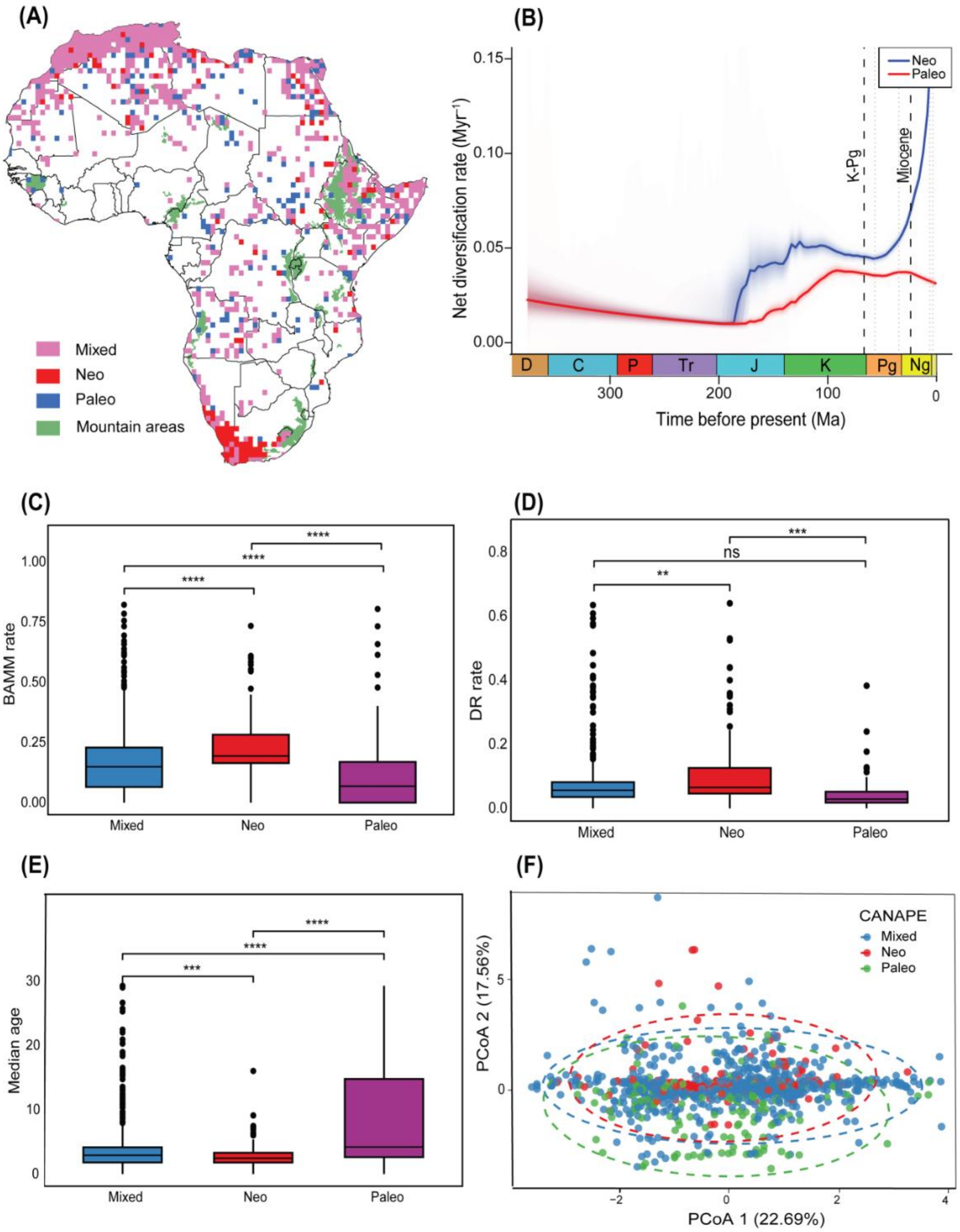


**Fig. S13.** Patterns of phylogenetic endemism and diversification across Africa. (A) Map showing results from the categorical analysis of neo- and paleo-endemism for vascular plant genera across Africa. Grid cells identified as centers of neo-endemism are shown in red, paleo-endemism in blue, and mixed endemism in purple, with mountain areas highlighted in green (data from GMBA Mountain Inventory v1.2; Körner et al., 2017). (B) Net diversification rates through time for neo-endemic (blue) and paleo-endemic (red) lineages. Shaded areas represent 95% confidence intervals. Vertical dashed lines indicate the Cretaceous–Paleogene boundary (K–Pg, 66 Ma) and the onset of the Miocene (23 Ma). Geological period abbreviations: D, Devonian; C, Carboniferous; P, Permian; Tr, Triassic; J, Jurassic; K, Cretaceous; Pg, Paleogene; Ng, Neogene. (C) Principal Coordinates Analysis (PCoA) of phylogenetic endemism assemblages based on compositional dissimilarity among grid cells. Points are colored according to CANAPE category (Mixed = purple, Neo = red, Paleo = blue), with 95% confidence ellipses for each group. Percent variation explained by each axis is indicated in parentheses.

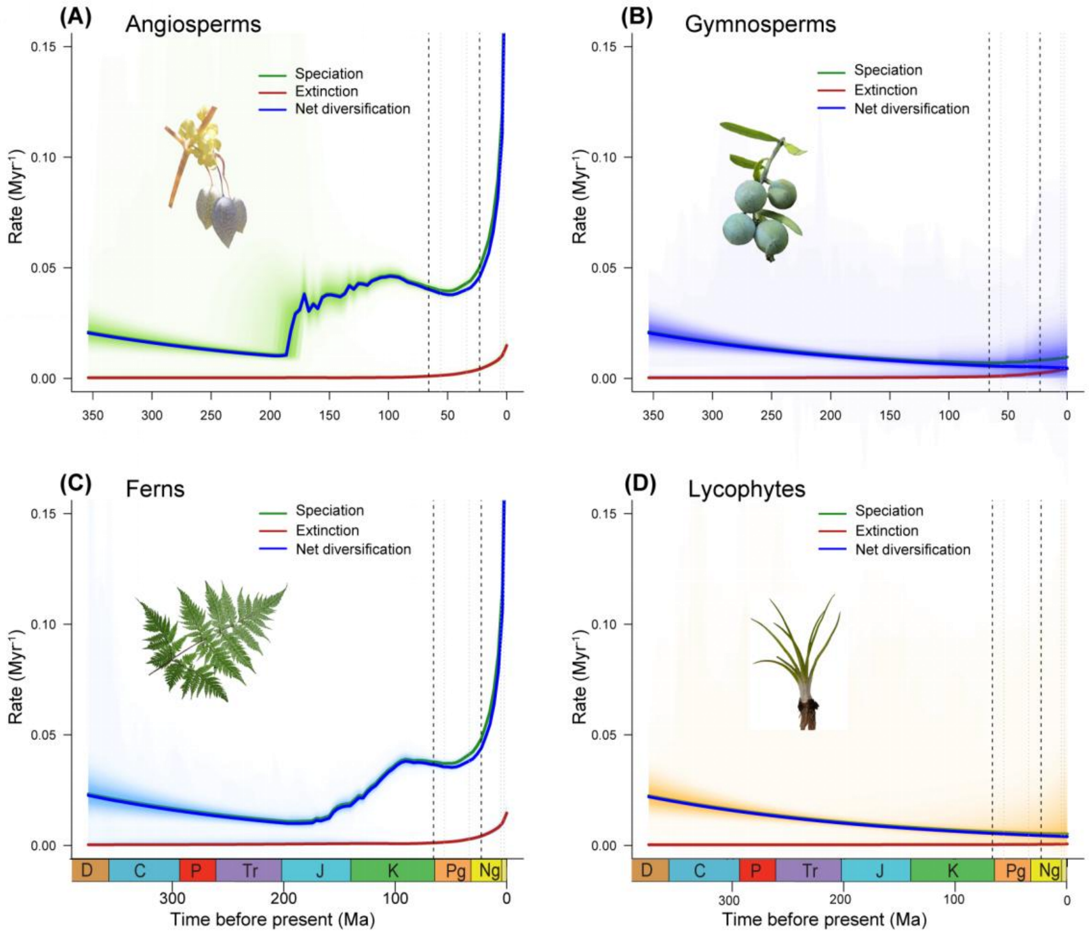


**Fig. S14. Rate-through-time (RTT) plots of vascular plant genera diversification.**
(**A–D**) RTT plots showing speciation, extinction, and net diversification rates for each major vascular plant group: angiosperms (A), gymnosperms (B), ferns (C), and lycophytes (D). Shaded areas represent 95% Bayesian credibility intervals. Vertical dashed lines indicate the Cretaceous–Paleogene boundary (K–Pg, 66 Ma) and the onset of the Miocene (23 Ma). Geological period abbreviations: D, Devonian; C, Carboniferous; P, Permian; Tr, Triassic; J, Jurassic; K, Cretaceous; Pg, Paleogene; Ng, Neogene. Representative genera are shown in each plot: (A) *Cysticapnos*, (B) *Afrocarpus*, (C) *Pteris*, and (D) *Isoetes*. Photo credits: Bing Liu and Inaturalist.

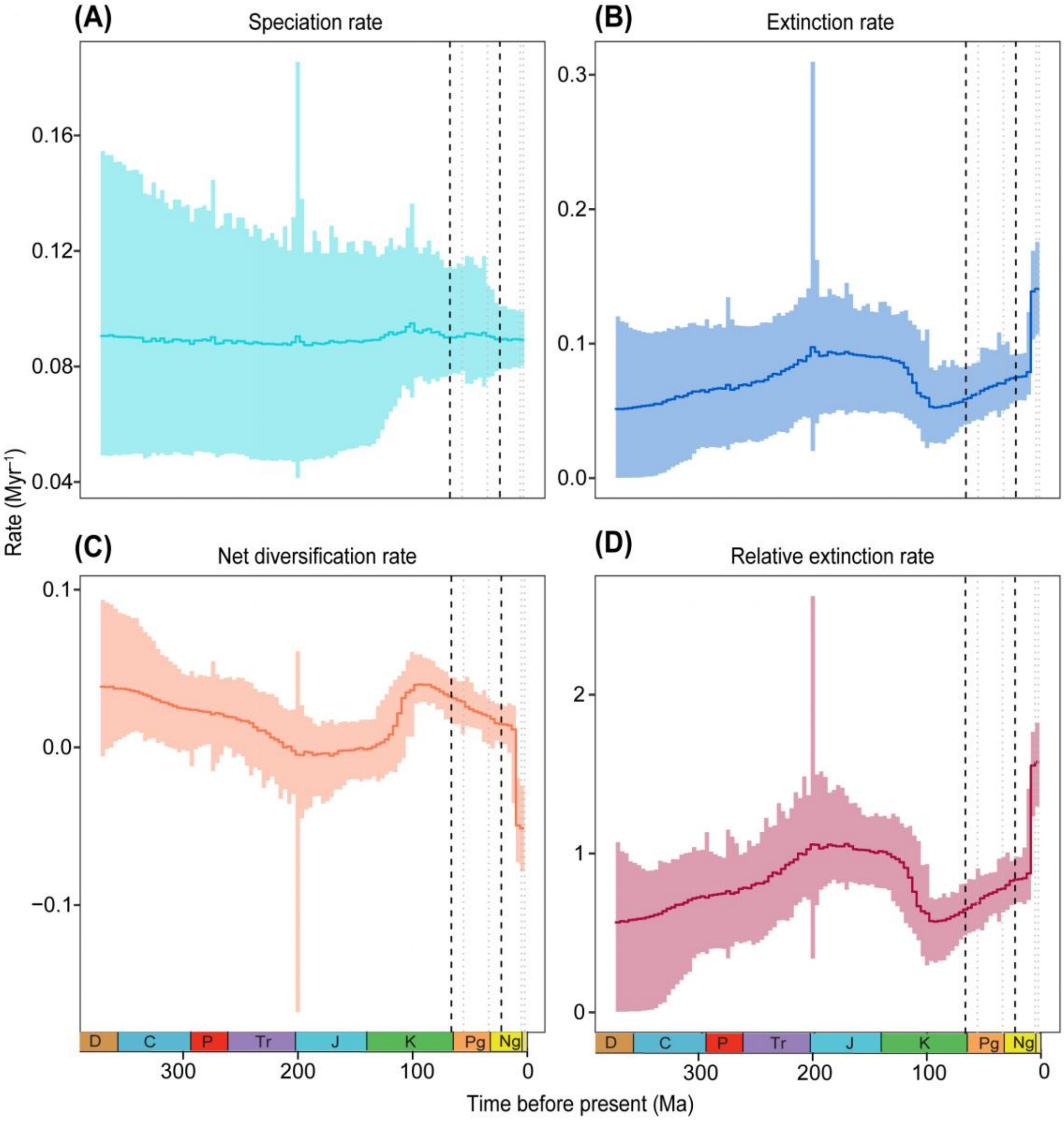


**Fig. S15.** Speciation, extinction, and net diversification rate for vascular plants in Africa based on the episodic birth-death model in RevBayes. Solid lines represent the mean estimate and colored areas around the 95% confidence interval. Vertical dashed lines indicate the Cretaceous–Paleogene boundary (K–Pg, 66 Ma) and the onset of the Miocene (23 Ma). Geological period abbreviations: D, Devonian; C, Carboniferous; P, Permian; Tr, Triassic; J, Jurassic; K, Cretaceous; Pg, Paleogene; Ng, Neogene.

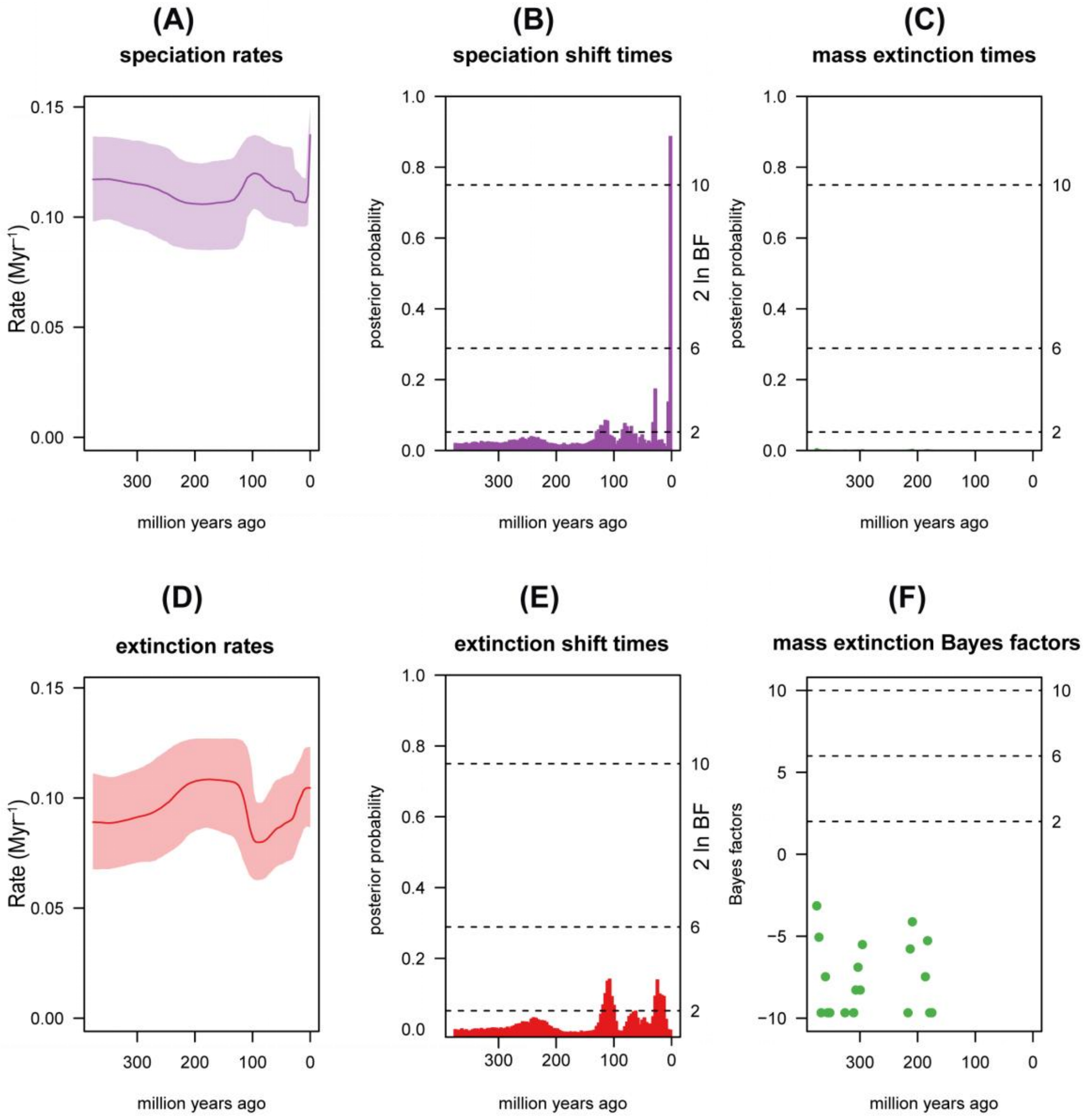


**Fig. S16.** Estimating rates of (and identifying shifts in) lineage diversification through time from TESS analysis, showing speciation shifts have higher posterior probability during the Neogene (ca. 23-2.58 Ma). Left: Plots of the posterior mean and 95% credible interval for the speciation and extinction rate (upper and lower panels, respectively). Right: Identifying temporal shifts in the speciation and extinction rate (upper and lower panels, respectively) using Bayes factors (BF) estimated by rjMCMC. Each bar indicates the posterior probability of at least one rate shift within that interval. Bars that exceed the specified significance threshold (2ln BF > 6) indicate significant rate shifts.

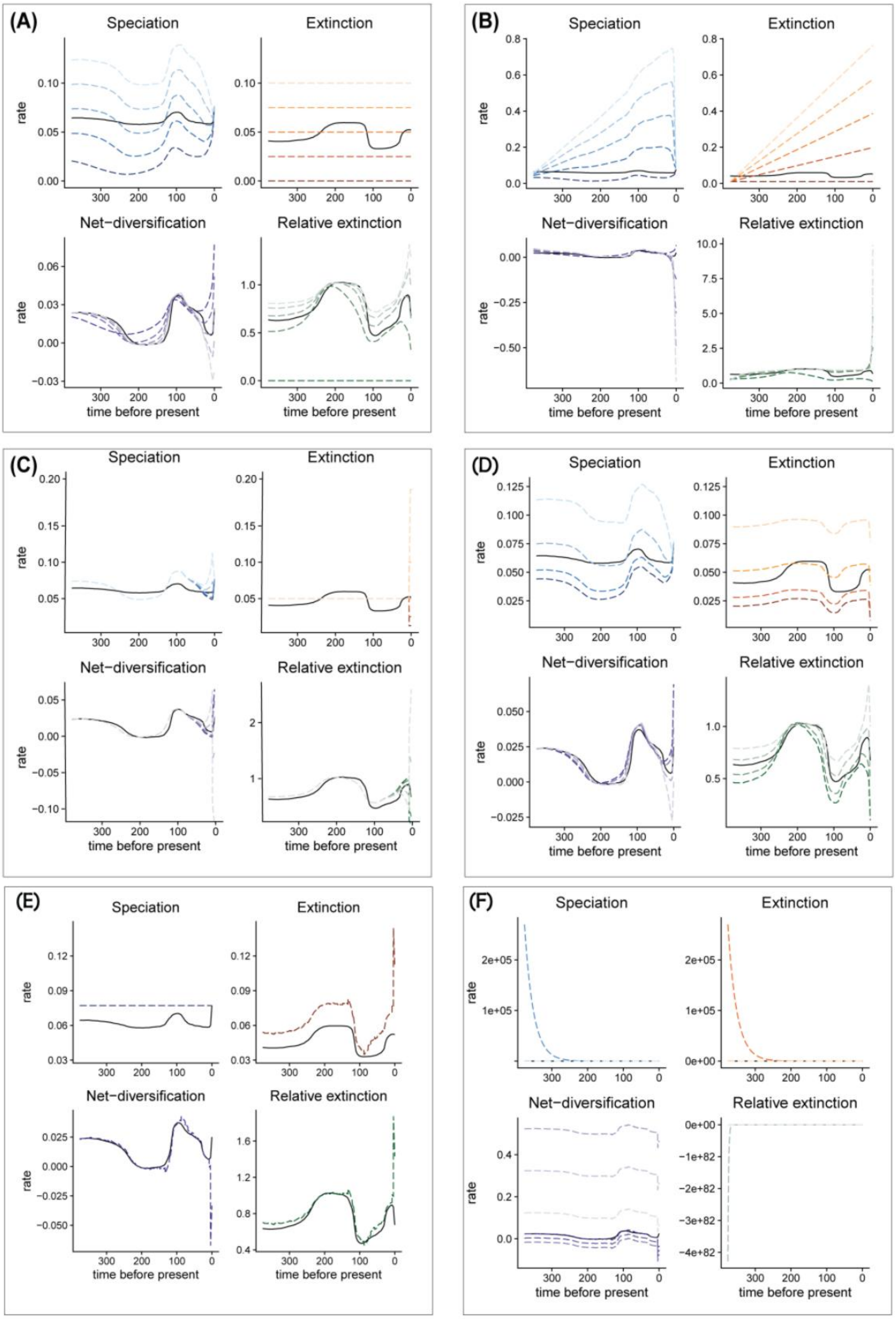


**Fig. S17 A. Scenario with constant extinction rates.**
Diversification rate functions within the congruence class assuming different constant extinction rate functions. The dashed lines show the different alternatives that were explored here, and the solid lines represent reference model. The overall pattern of speciation rates as well as net-diversification rates are robust to alternative constant extinction rates within the congruence class.

Only the relative extinction rate changes, which is not surprising since we explored different extinction rates.

**S17B. Scenario with linearly increasing extinction rates.**
Diversification rate function for different linear extinction rates within the same congruence class. The dashed lines show the different alternatives that we explored here, and the solid lines represent our reference model. We observe that the speciation and net diversification rates are robust to different linear extinction rates within the congruence. As expected, the relative extinction rate changes because of different extinction rates.

**S17C. Scenario with two-epoch extinction rates.**
Diversification rate functions for two-epoch extinction rate functions within the congruence class. The solid line shows the reference model, and the dashed lines show the alternative models. We observe no noticeable effect on the overall trend of the derived speciation rates.

**S16D. Scenario with reversed trend extinction rates.**
Diversification rate functions for a reversed trend extinction rate function. The solid line shows the reference model, and the dashed lines show the alternative model. We observe no noticeable effect on the overall trend of the derived speciation rates.

**S17E. Scenario with constant speciation rates.**
Diversification rate functions for a constant speciation rate within the congruence class. The solid line shows our reference model, and the different dashed line shows the alternative model. We observe that for the constant speciation rate function we compute an extinction rate that was negative multiple times during the clade's evolutionary history.

**S17F. Scenario with exponential speciation rates.**
Diversification rate functions for exponentially increasing/decreasing speciation rate functions within the congruence class. The solid line shows the reference model, and the dashed lines show the alternative models. We observe that for all speciation rate functions that the derived extinction rate functions are negative towards the present and/or at some other point in time

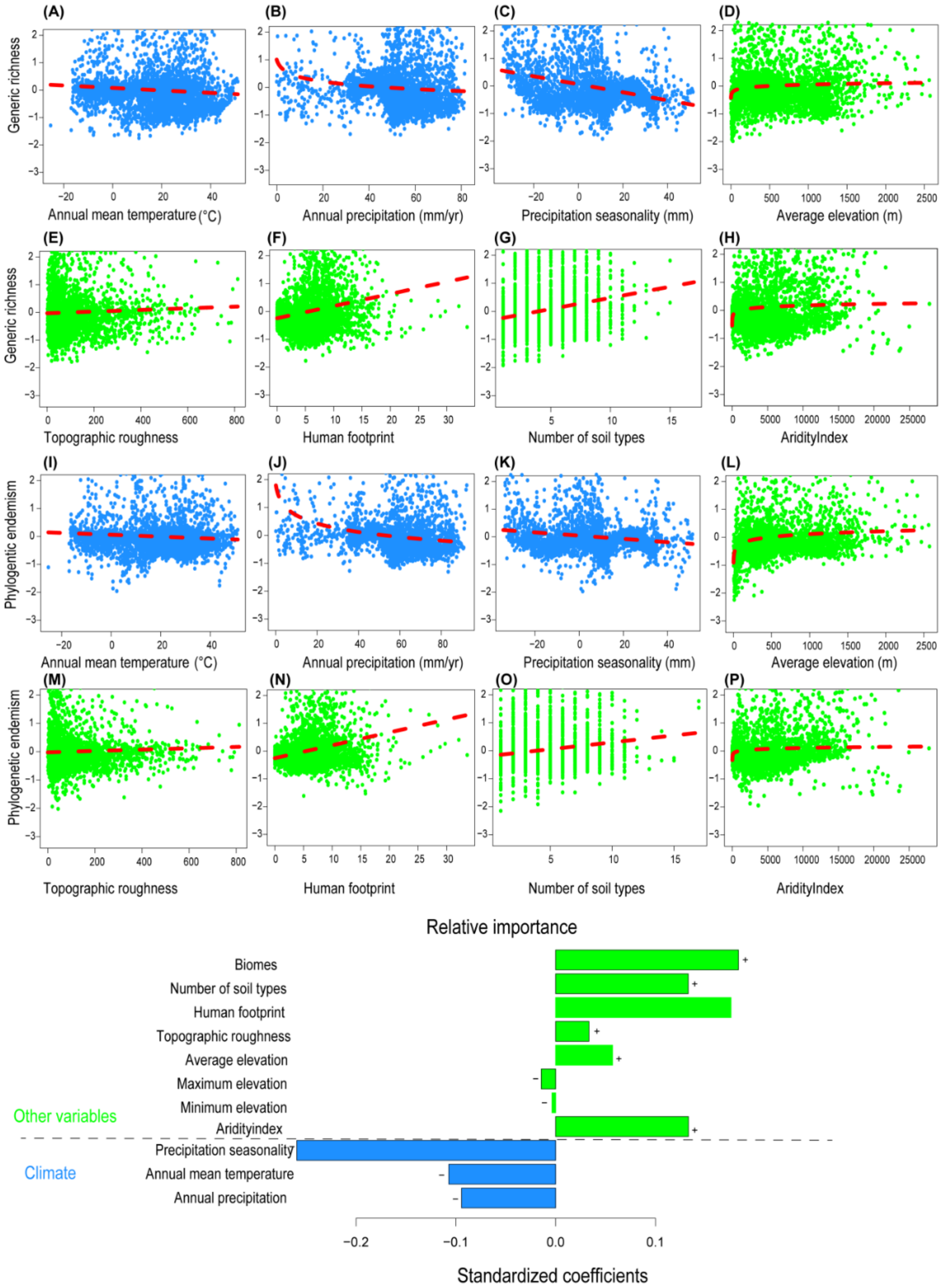


**Fig. S18. Determinants of biodiversity and endemism patterns in Africa**. (**A–H**) Relationships between environmental and anthropogenic predictor variables (aridity index, climate, elevation, human footprint, and soil) and generic richness of vascular plant genera across Africa. (**I–P**) Relationships between the same predictor variables and phylogenetic endemism of vascular plant genera. Partial residual plots illustrate the relationship between each response variable and its corresponding predictor while statistically controlling for all other predictors in the modelBars in the relative importance plots indicate the standardized coefficients of predictor variables, with the direction of effect denoted by (+) for positive and (–) for negative associations.

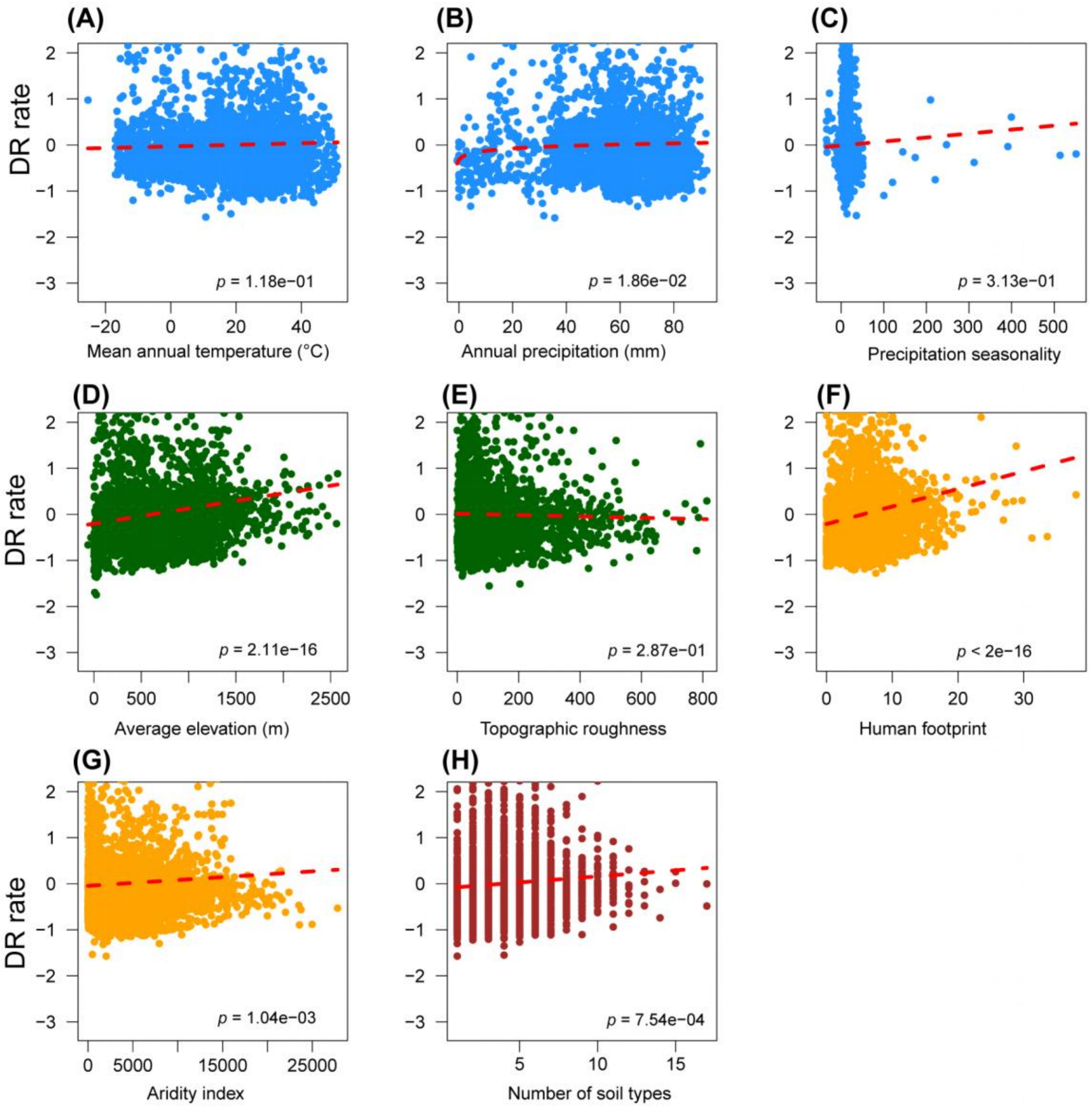


**Fig. S19. Environmental and anthropogenic determinants of diversification rates across Africa**. (**A–H**) Relationships between environmental and anthropogenic predictor variables (aridity index, climate, elevation, human footprint, and soil) and tip rates from DR statistic of vascular plant genera across Africa. Partial residual plots illustrate the relationship between diversification rate and each predictor variable while statistically controlling for the effects of all other predictors in the model.

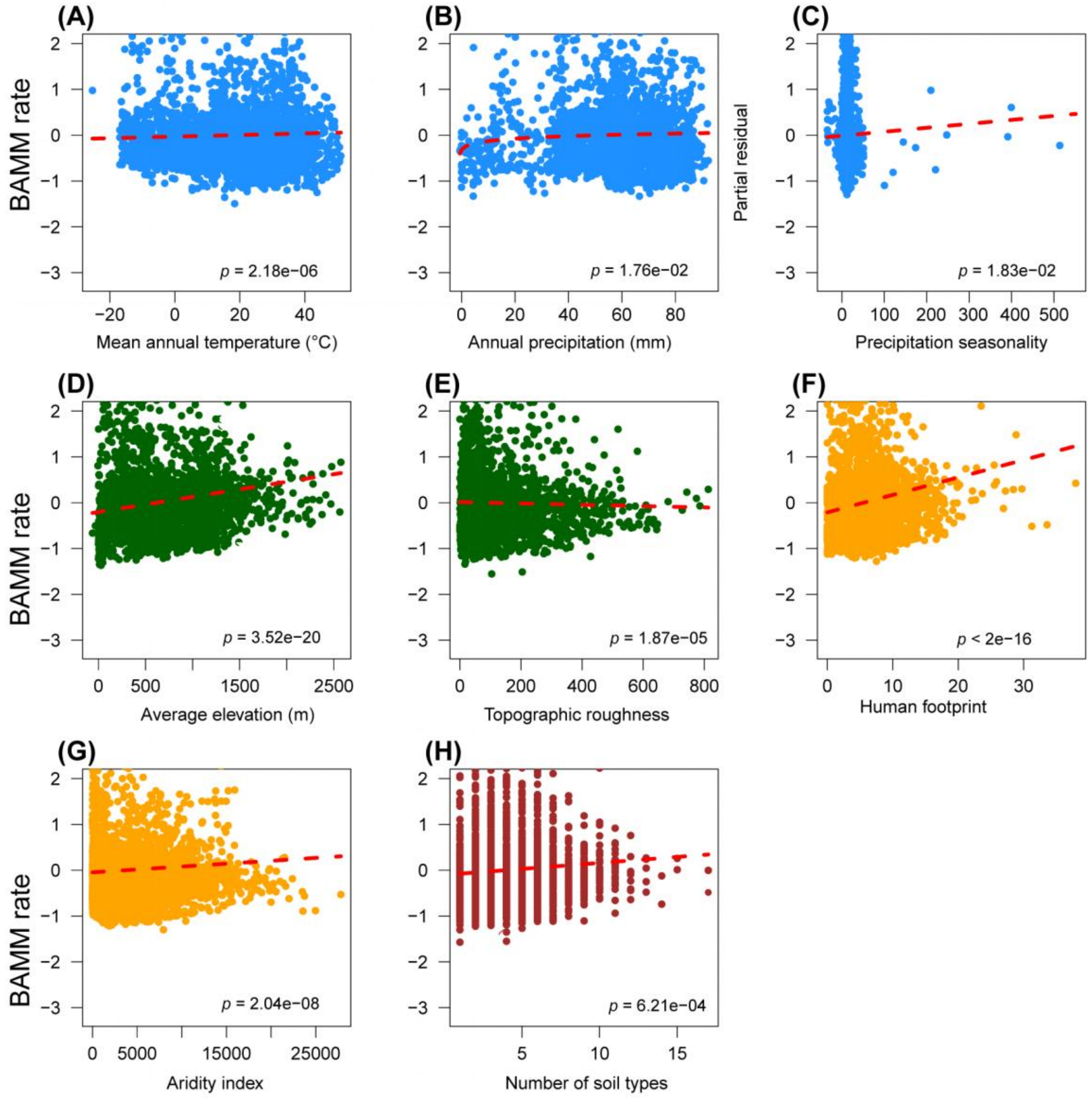


**Fig. S20. Environmental and anthropogenic determinants of diversification rates across Africa**. (**A–H**) Relationships between environmental and anthropogenic predictor variables (aridity index, climate, elevation, human footprint, and soil) and tip rates from BAMM of vascular plant genera across Africa. Partial residual plots illustrate the relationship between diversification rate and each predictor variable while statistically controlling for the effects of all other predictors in the model.

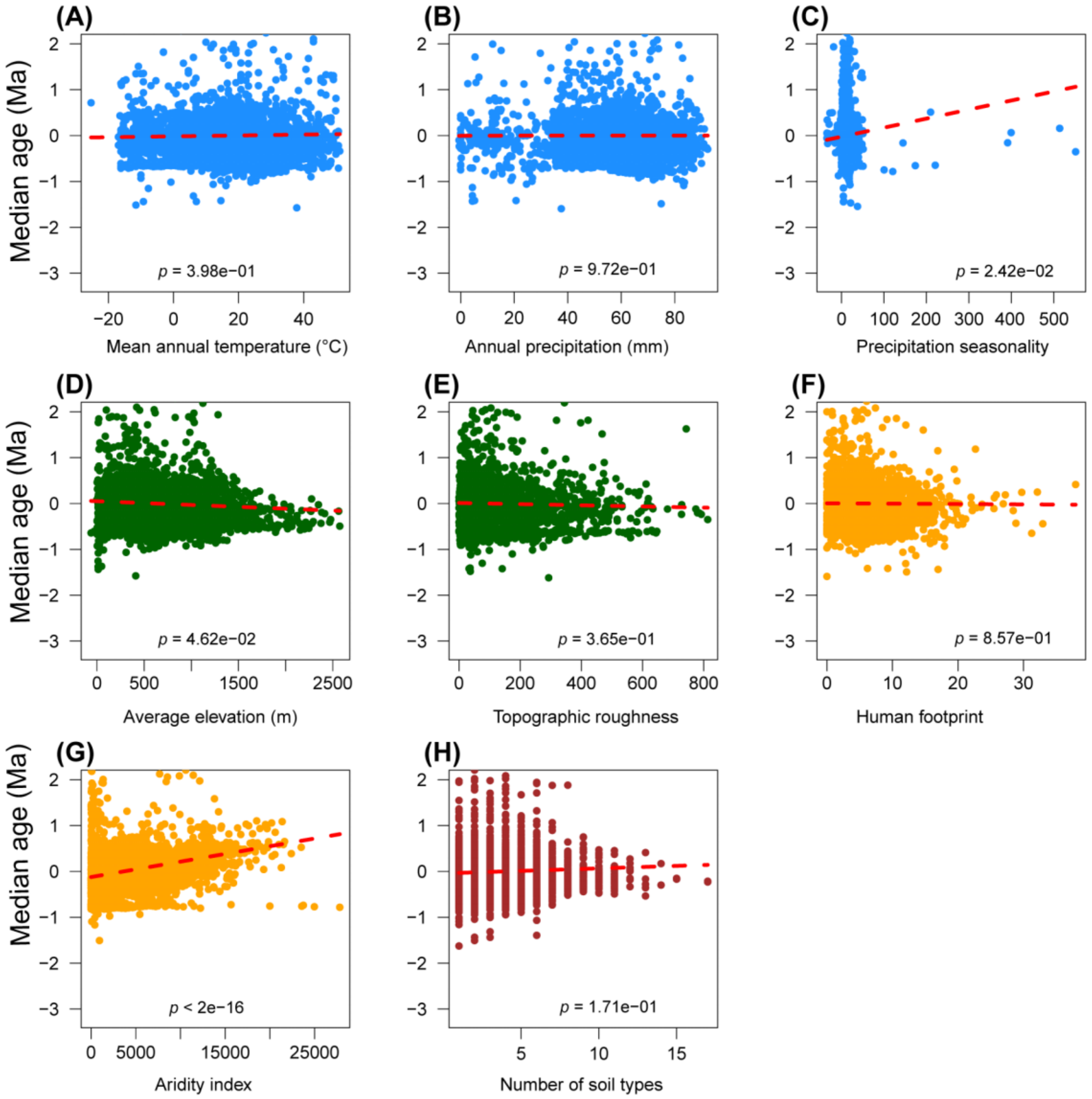


**Fig. S21. Environmental and anthropogenic determinants of median lineage age across Africa.** **(A–H)** Relationships between environmental and anthropogenic predictor variables (aridity index, climate, elevation, human footprint, and soil) and the median lineage age of vascular plant genera across Africa. Partial residual plots illustrate the relationship between median lineage age and each predictor variable while statistically controlling for the effects of all other predictors in the model**.**

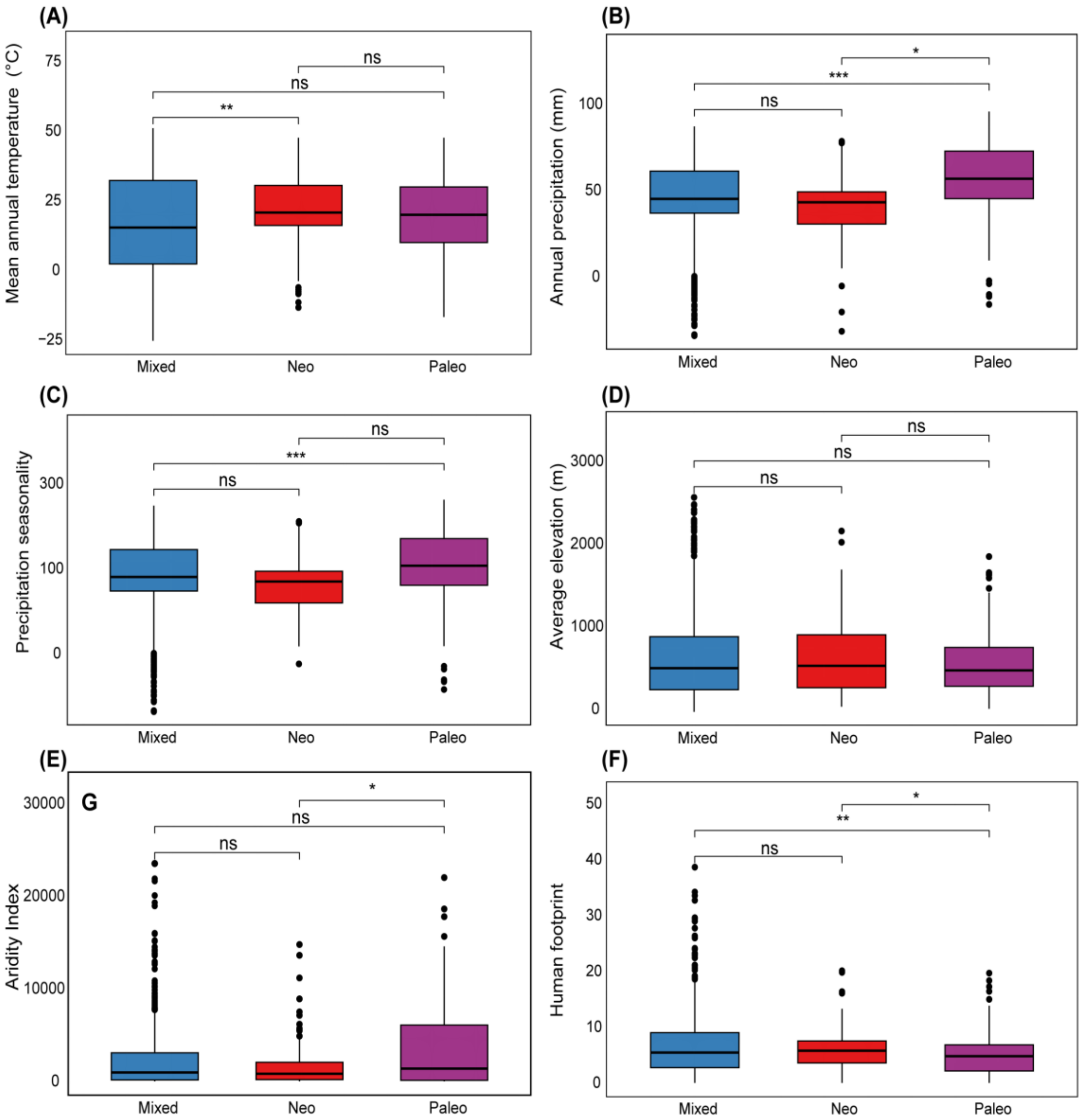


**Fig. S22. Environmental, anthropogenic, and evolutionary correlates of centers of endemism in Africa.** (**A–F**) Boxplots showing the relationship between eight predictor variables and centers of neo-(red), paleo-(blue), and mixed (purple) endemism based on CANAPE analysis. Variables include mean annual temperature (A), annual precipitation (B), precipitation seasonality (C), average elevation (D), aridity index (E), and human footprint (F). Asterisks indicate statistical significance (*$p < 0.05$; **$p < 0.01$; ***$p < 0.001$; ns = not significant) based on Kruskal–Wallis tests followed by pairwise Wilcoxontests.

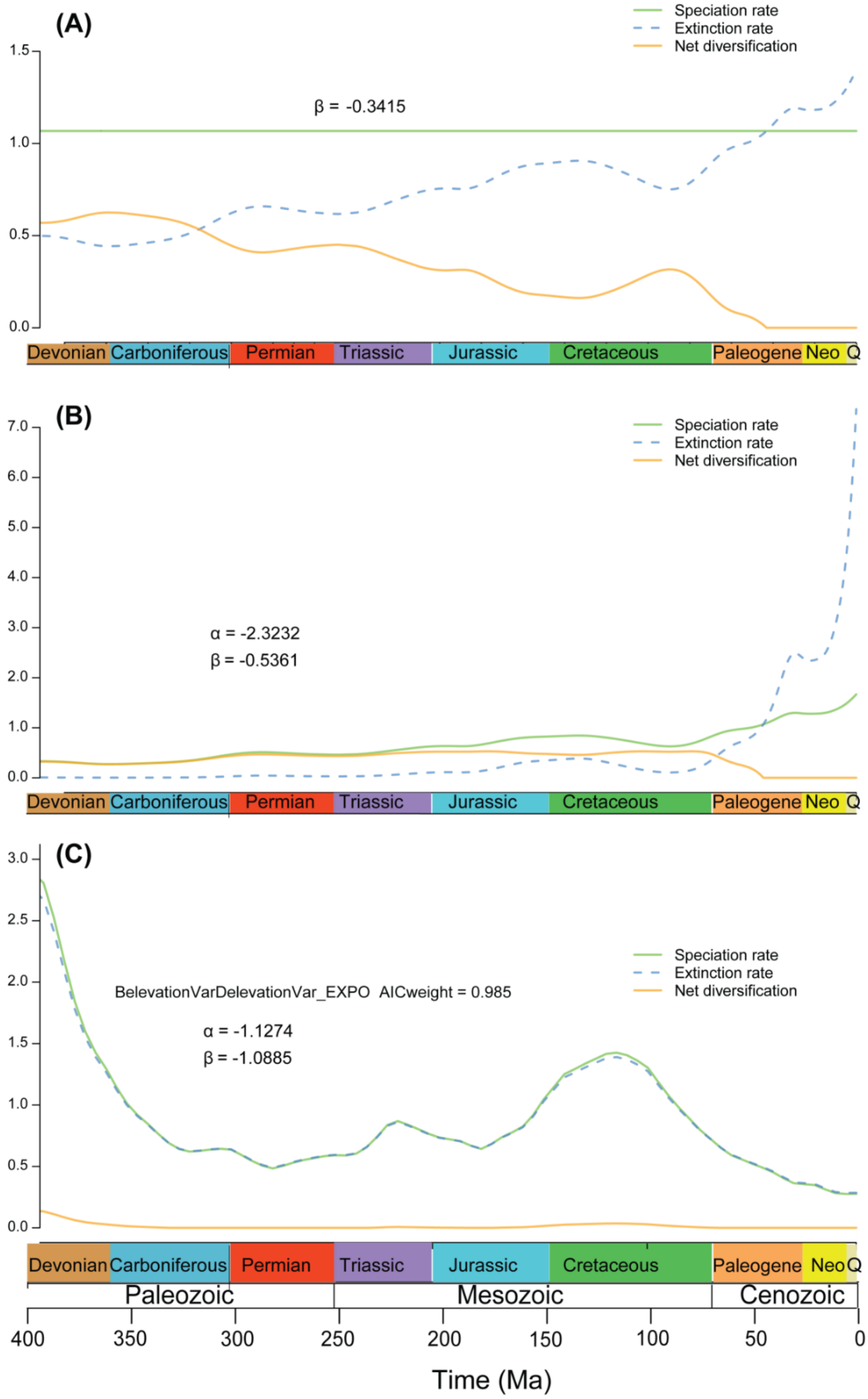


**Fig. S23. Paleoenvironment-dependent diversification dynamics inferred using RPANDA.** (A) Temperature-dependent model, (B) atmospheric $CO_2$–dependent model, and (C) elevation-dependent model. Time-dependent trajectories of speciation rate (solid green lines), extinction rate (blue dashed lines), and net diversification rate (orange lines) estimated for African vascular plant genera under different environmental predictors. Model names and corresponding Akaike weights (AICw) are shown within each panel

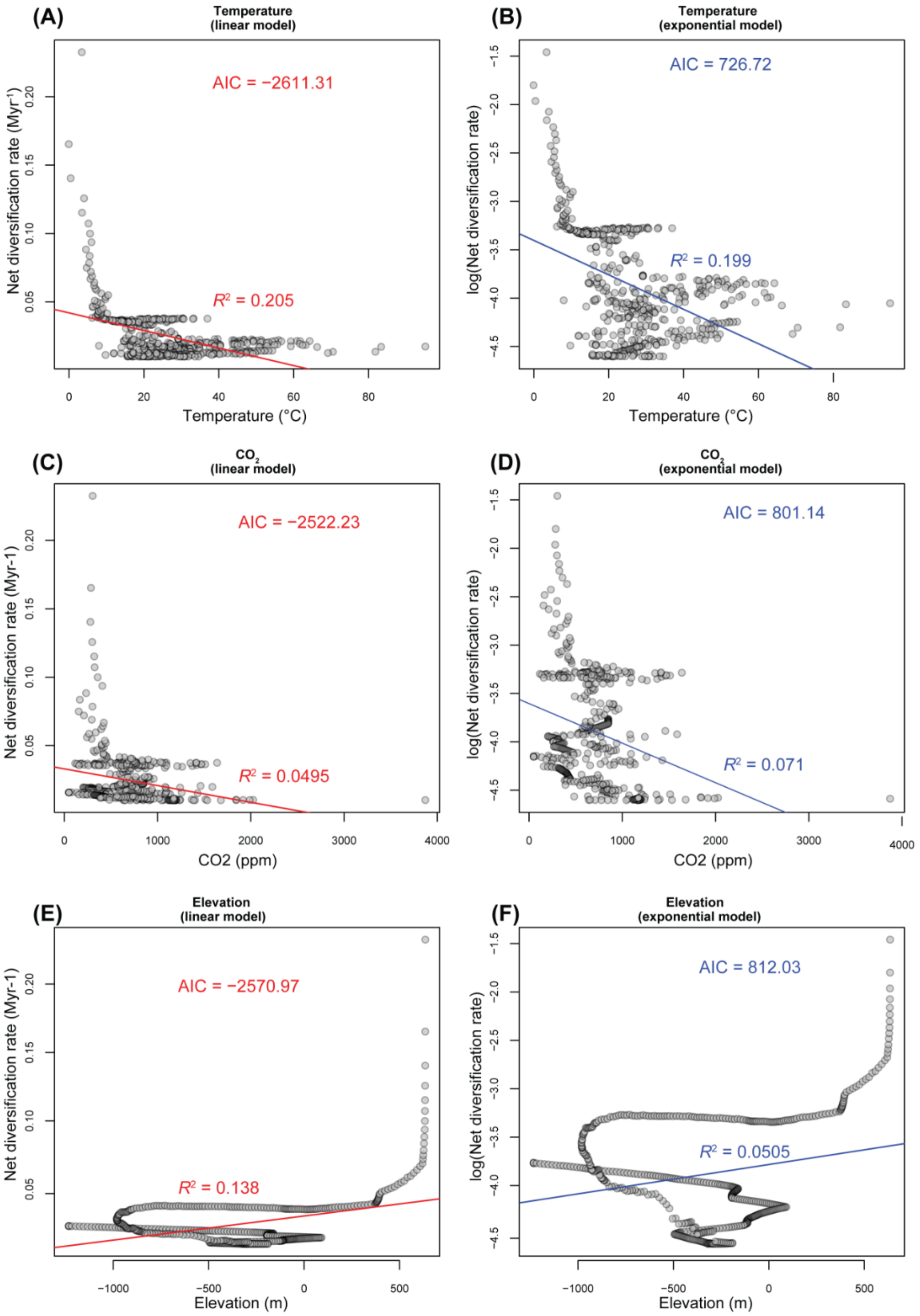


**Fig. S24.** Correlation between BAMM tree-wide net diversification rates and historical global temperature, atmopsheric $CO_2$, and paleo-elevation based on linear and exponential best-fit models. Panels A, C, and E represent the best linear model with AIC and $R^2$ as indicated; similarly, and panels B, D, and F represent the best exponential model with AIC and $R^2$ as indicated.

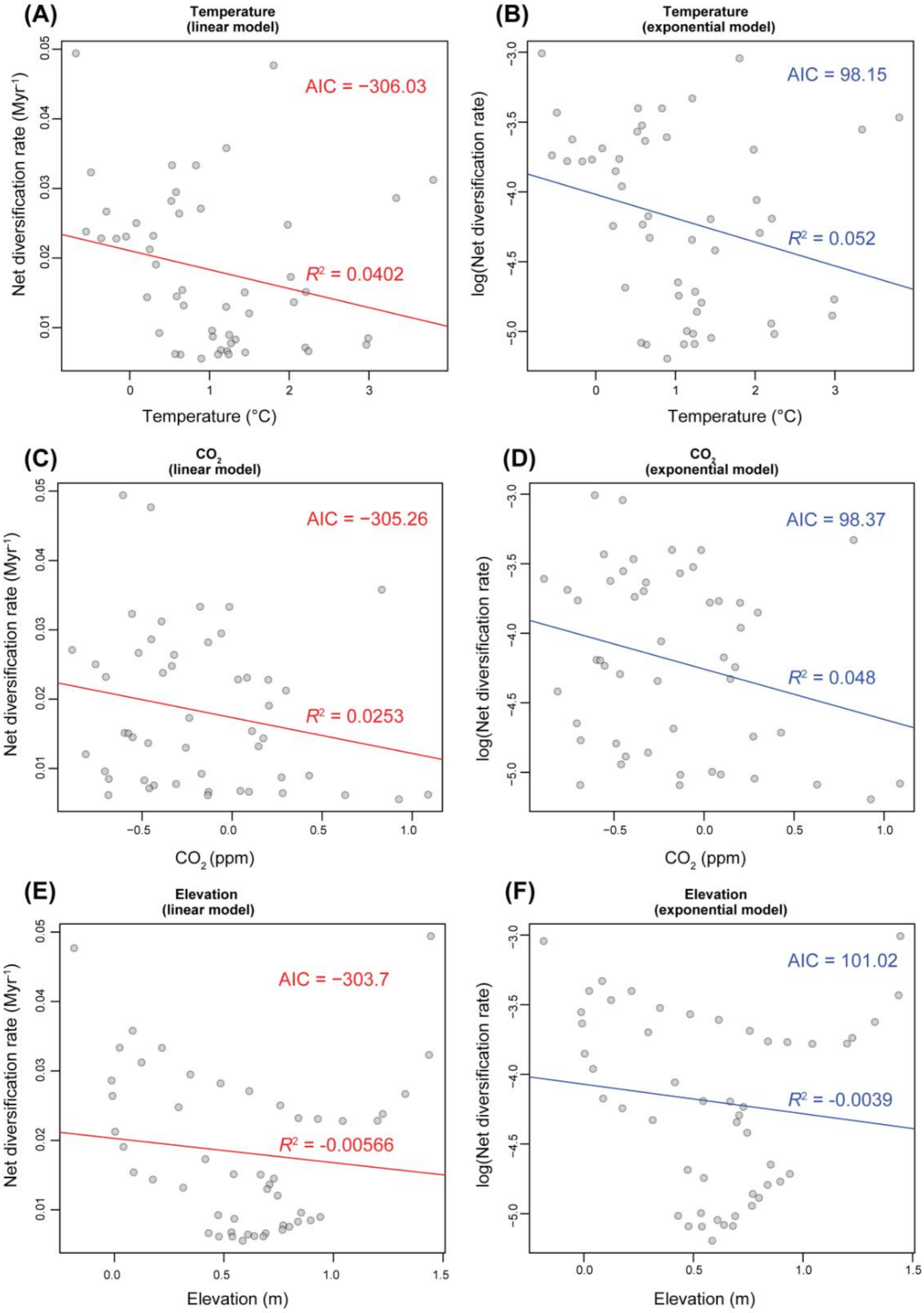


**Fig. S25.** Correlation between ClaDS tree-wide net diversification rates and historical global temperature, atmopsheric $CO_2$, and paleo-elevation based on linear and exponential best-fit models. Panels A, C, and E represent the best linear model with AIC and $R^2$ as indicated; similarly, and panels B, D, and F represent the best exponential model with AIC and $R^2$ as indicated.

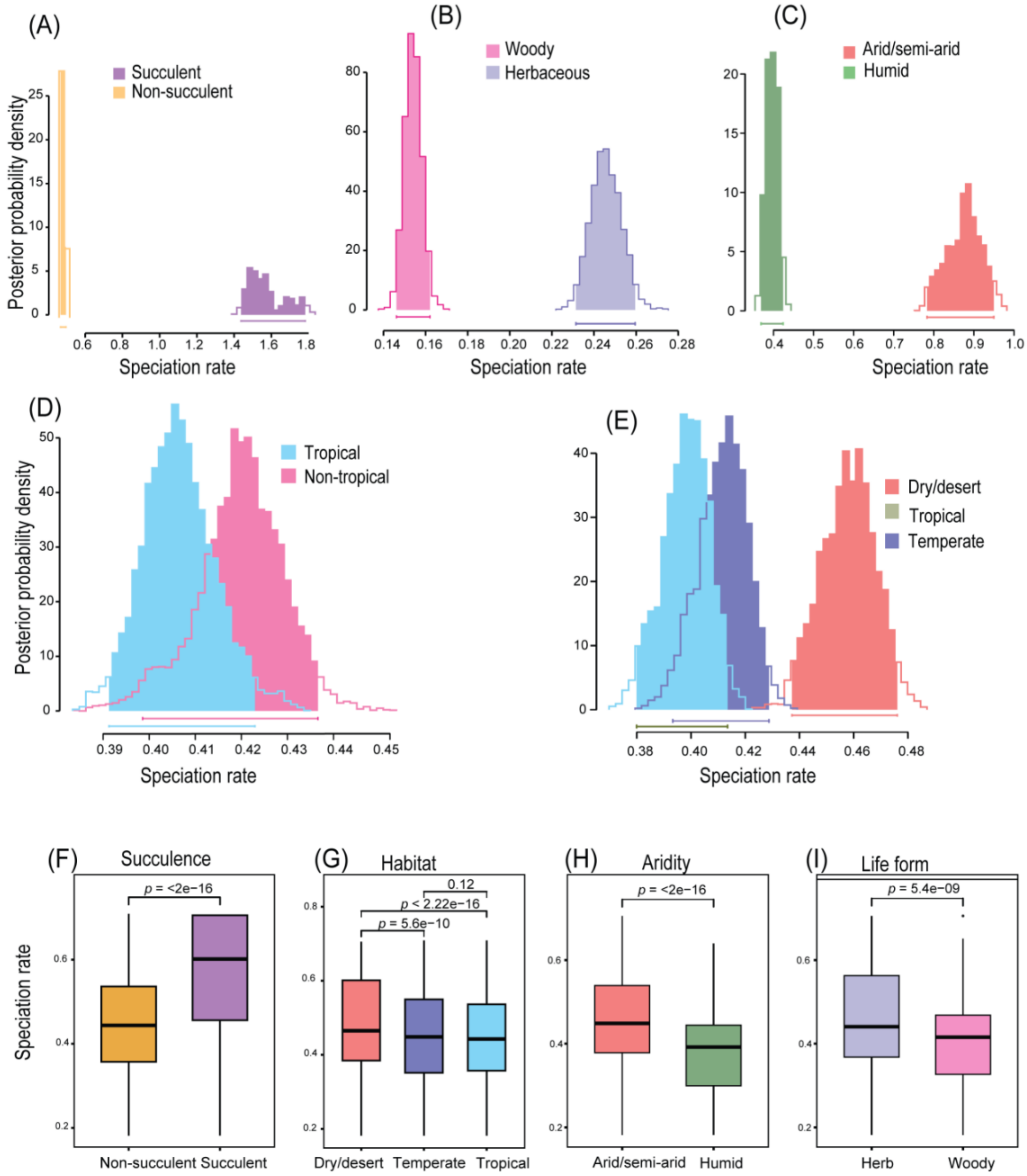


**Fig. S26. Biotic and abiotic drivers of speciation for vascular plant genera in Africa.** **(A–E)** Speciation analyses using binary state speciation and extinction (BiSSE; A–D) and multistate speciation and extinction (MuSSE; E) models showing the effects of two abiotic and two biotic traits on speciation rate: succulence (A), life form (B), aridity (C), and habitat (D, E). (**F–I**) Diversification estimated by BAMM across four different trait categroies of African vascular plant lineages.

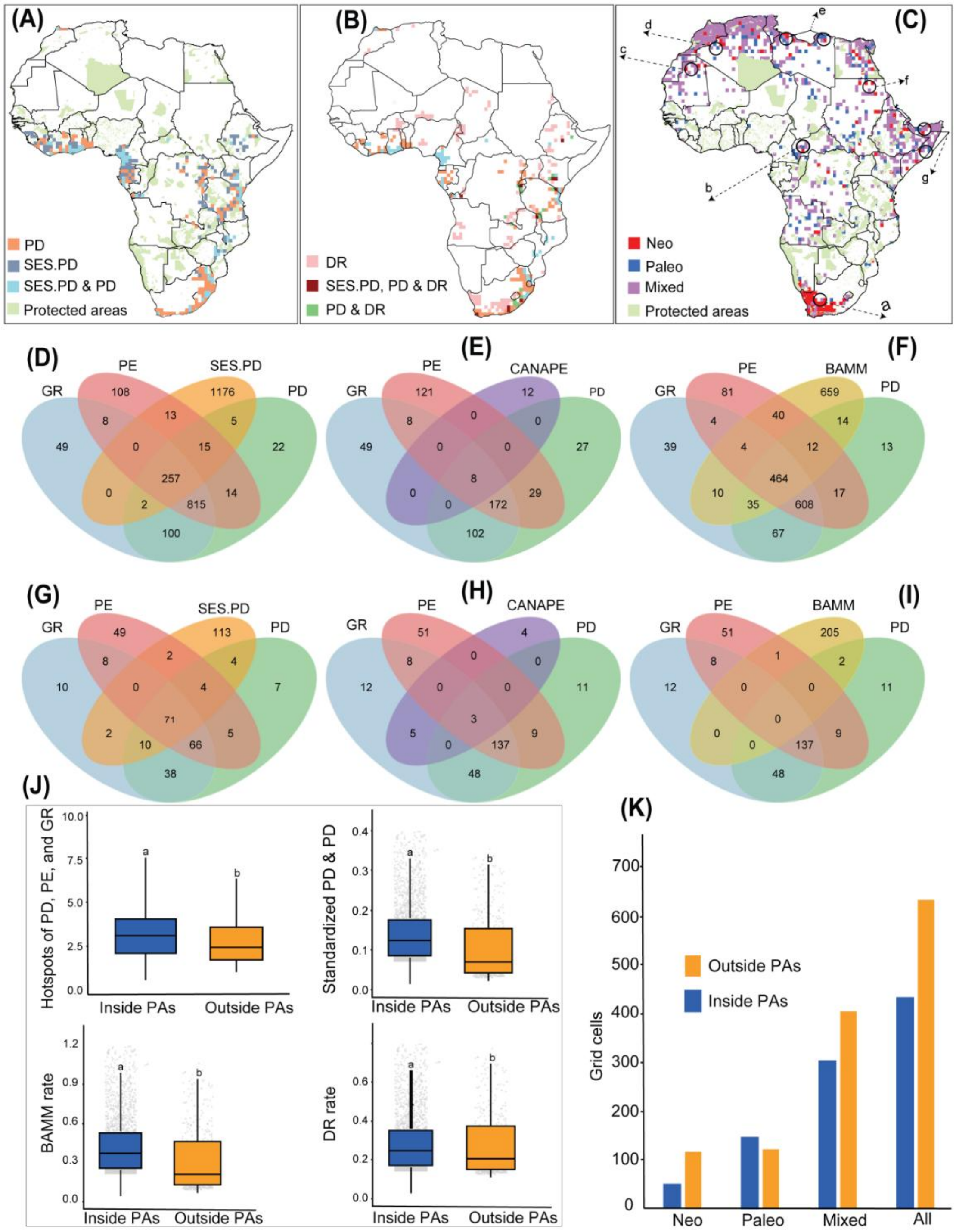


**Fig. 27. Conservation priorities for vascular plant genera in Africa.** (A-C) Grid cells representing the top 5% of phylogenetic diversity (PD, brown) and standardized PD (SES.PD, blue), with overlapping cells in gray. Protected areas are highlighted in light green. (B) Grid cells representing top 5% of diversification rate (BAMM and DR statistic, pink), all three indices (SES.PD, PD, and DR; deep red), and overlap between PD and DR (green). (C) Conservation priorities identified using CANAPE analysis: neo-endemism (red), paleo-endemism (blue), and mixed endemism (purple). Letters (a–g) indicate selected priority areas. Maps of protected areas are adapted from the World Database on Protected Areas (WDPA; https://www.protectedplanet.net/, accessed June 2025). (D–I), Venn diagrams showing overlap and uniqueness of grid cells across the 30% criterion (D–F) and 5% criterion (G–I) hotspots for GR, PD, PE, SES.PD, CANAPE, and BAMM rate. (J–K) Boxplots

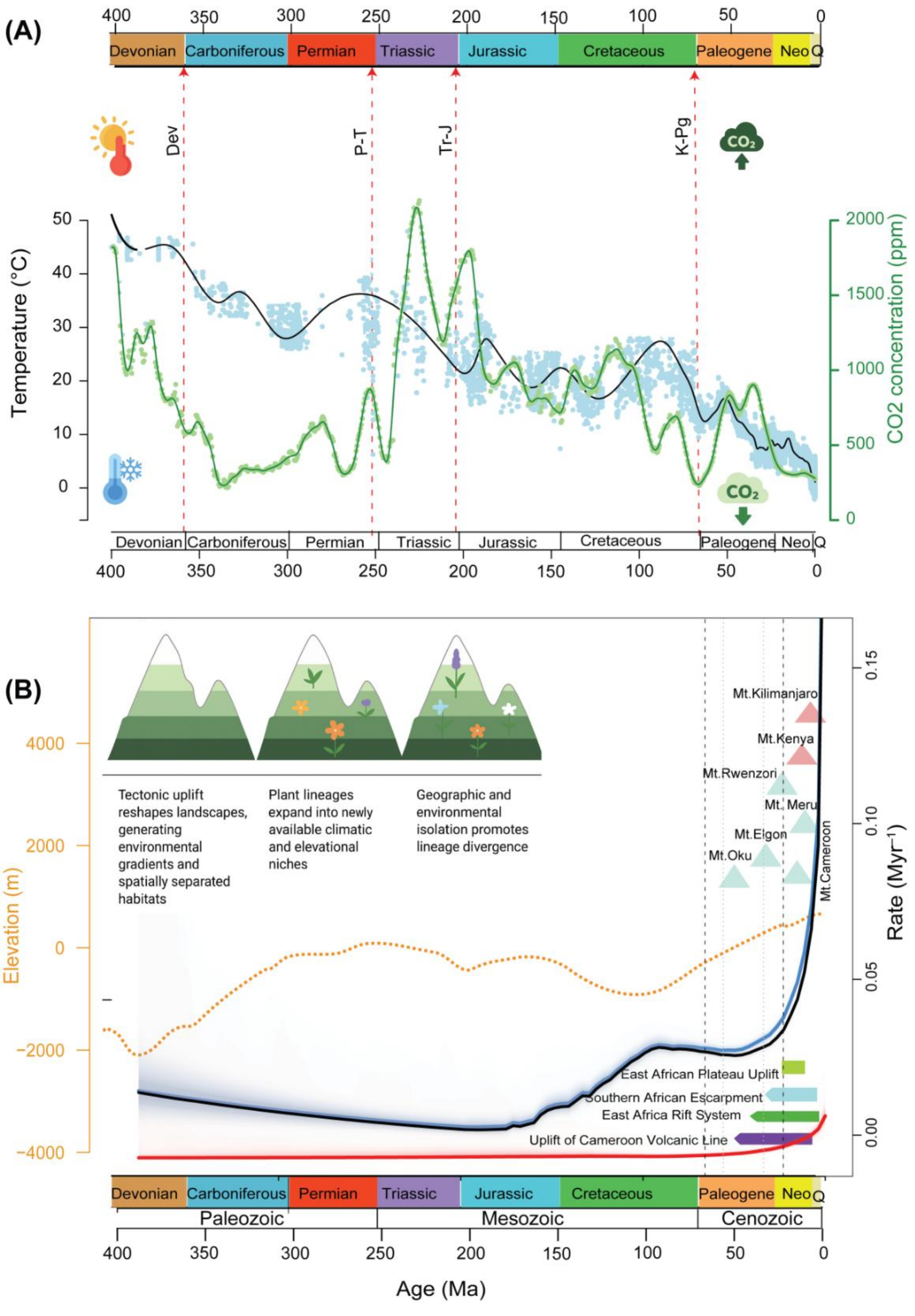


**Fig. 28. Climate, tectonics, and mountain uplift as drivers of evolutionary opportunity through time.** (A) Global temperature (black line with blue uncertainty envelope) and atmospheric $CO_2$ concentration (green line) over the last ~400 million years, shown alongside major geological periods. Vertical dashed red lines indicate key mass extinction events (Devonian, Permian–Triassic [P–T], Triassic–Jurassic [Tr–J], and Cretaceous–Paleogene [K–Pg]). (B) Empirical curves (bottom) showing generalized elevation change (orange dashed line) and inferred net diversification rate through time (adopted from Fig. 4). Colored bars indicate major

uplift phases in Africa, including the Cameroon Volcanic Line, East African Rift System, Southern African Escarpment uplift, and East African Plateau uplift. Gray triangles mark the emergence of major East African mountains (e.g., Mt. Elgon, Mt. Meru, Mt. Kenya, Mt. Kilimanjaro), highlighting the close temporal association between mountain building and increased diversification toward the present.

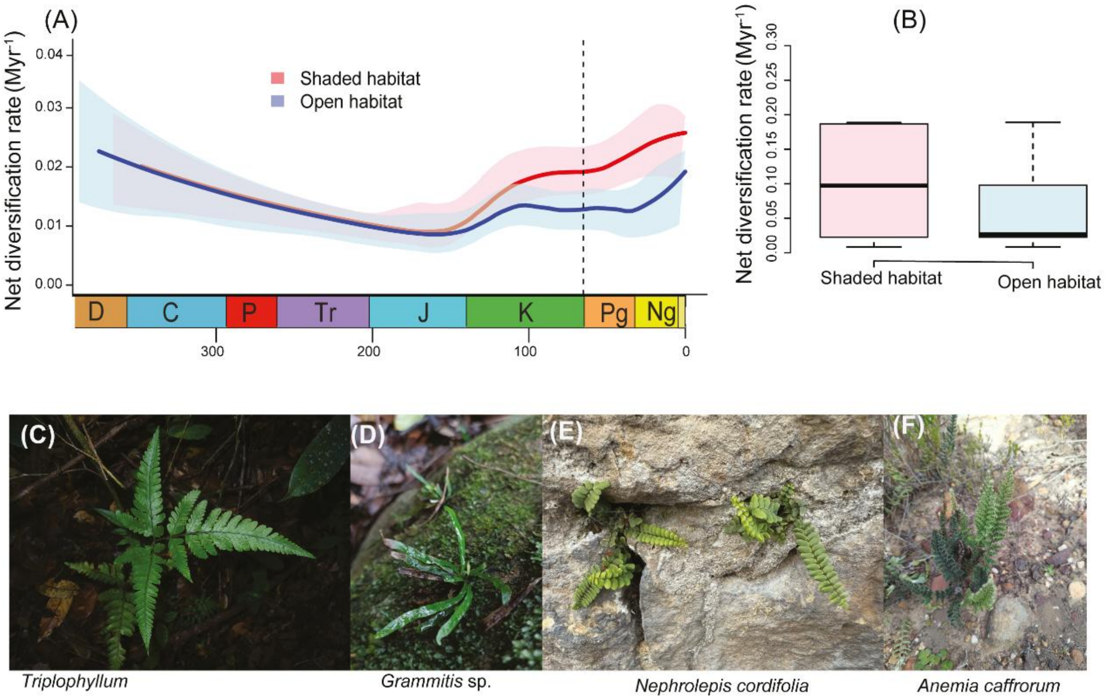


**Fig. S29. Diversification dynamics of ferns in shaded and open habitats.** (A) Clade-wide net diversification rates of ferns adapted to shaded (red) and open (blue) habitats through time, as estimated by Bayesian Analysis of Macroevolutionary Mixtures (BAMM). The vertical dashed line at 65.5 Ma marks the approximate establishment of angiosperm-dominated forest canopies (based on fossil evidence; Jacobs, 2004; Johnson & Ellis, 2002), when the confidence intervals of diversification rates for shaded- and open-adapted lineages began to diverge. (B) Boxplots showing diversification rates for ferns from shaded and open habitats inferred using BAMM. Habitat classifications were derived from Wu et al. (2025). (C–F) Representative examples of fern species from different habitats: (C–D) shaded habitats and (E–F) open habitats. Representative species are indicated in each panel. Photo credit: iNaturalist.

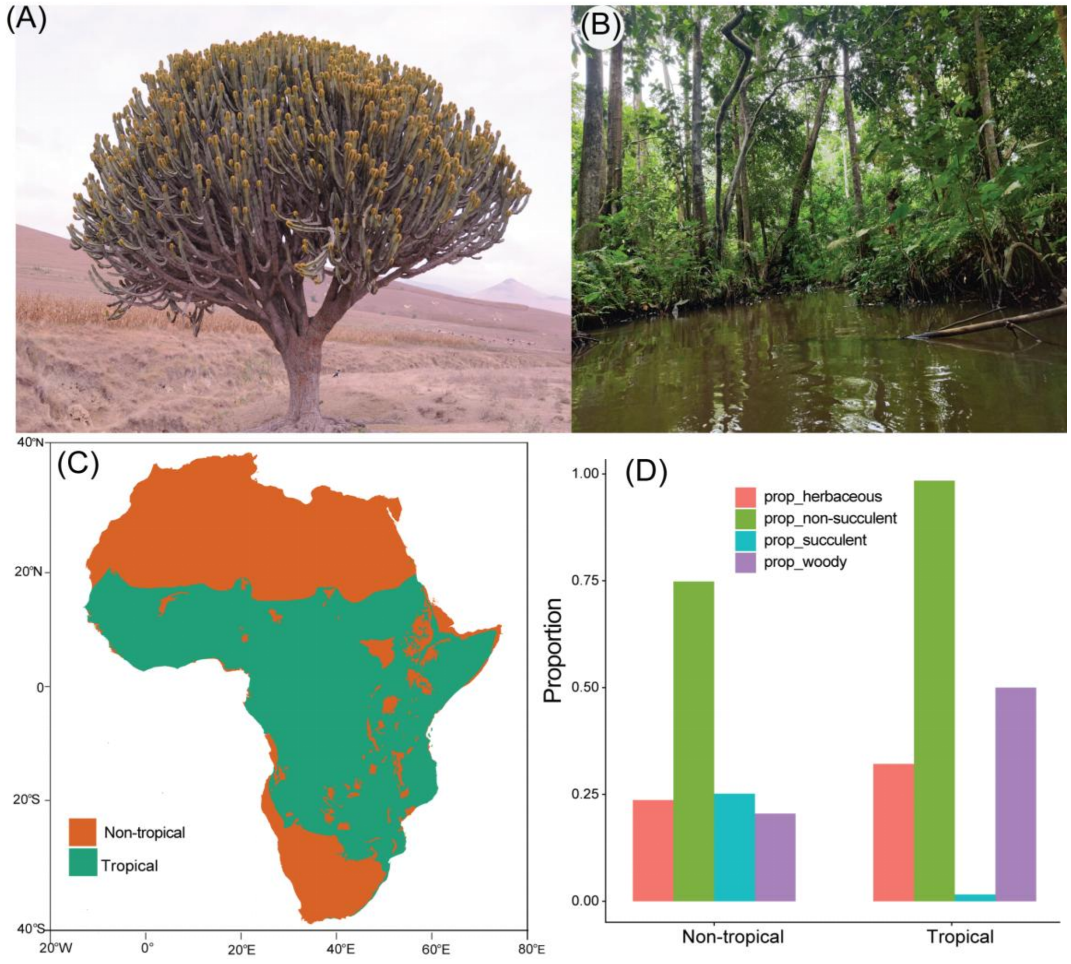


**Fig. S30. Contrasting vegetation structure and growth-form composition between tropical and non-tropical Africa.** (A-B) Representative vegetations of non-tropical Africa, illustrating open, arid to semi-arid systems (A), and tropical Africa, showing humid lowland forest characterized by closed canopies and woody plant dominance (B). (C) Map of Africa showing the spatial delineation of tropical (green) and non-tropical (orange) regions used in the analyses. (D) Proportional representation of major growth forms—herbaceous, woody, succulent, and within tropical and non-tropical regions.

### 1. Supplementary Tables

**Table S1**. Calibrations used for divergence time estimates of the vascular plants of Africa (MRCA: most recent common ancestor, Ma: million years ago, CG: crown group, SG: stem group).

| **Clade** | **MRCA** | min/ max | Age (Ma) | **References** |
|---|---|---|---|---|
| CG Angiosperms | Nymphaeaceae_*Nymphaea_nouchali_var_caerulea,* Lamiaceae_*Clinopodium_alpinum* | min | 136 | Ref.[1–4] |
| | | max | 209 | Ref.[5] |
| Vacsular plants | Lycopodiaceae_*Huperzia_filiformis,* Lamiaceae_*Clinopodium_alpinum* | min | 423 | Ref.[4] |
| SG Gymnosperms | Zamiaceae_*Encephalartos_equatorialis,* Lamiaceae_*Clinopodium_alpinum* | min | 350 | Ref.[5] |
| CG Solanaceae | Solanaceae_*Nicotiana_alata*, Solanaceae_*Solanum_pronum* | min | 52.2 | Ref.[6] |
| CG Convolvulaceae | Convolvulaceae_*Ipomoea_coccinea,* Convolvulaceae_*Distimake_dissectus* | min | 47.8 | Ref.[7] |
| CG Sapotaceae | Sapotaceae_*Gluema_ivorensis,* Sapotaceae_*Spiniluma_oxyacantha* | min | 56 | Ref.[8] |
| SG Olacaceae | Olacaceae_*Olax_pentandra,* Balanophoraceae_*Mystropetalon_thomii* | min | 66 | Ref.[9] |
| SG Polygonaceae | Polygonaceae_*Rumex_spinosus,* Plumbaginaceae_*Plumbago_auriculata* | min | 66 | Ref.[10] |
| SG Rhamnaceae | Rhamnaceae_*Ziziphus_pubescens,* Barbeyaceae_*Barbeya_oleoides* | min | 72.1 | Ref.[11] |
| CG Rhamnaceae | Rhamnaceae_*Rhamnus_prinoides,* Rhamnaceae_*Noltea_africana* | min | 48.6 | Ref.[12,13] |
| CG Moraceae | Moraceae_*Milicia_regia,* Moraceae_*Ficus_mucuso* | min | 47.8 | Ref.[14] |
| CG Rhizophoraceae | Rhizophoraceae_*Rhizophora_mangle,* Rhizophoraceae_*Cassipourea_lescotiana* | min | 33.9 | Ref.[15] |
| CG Chrysobalanaceae | Chrysobalanaceae_*Chrysobalanus_icaco,* Chrysobalanaceae_*Parinari_excelsa* | min | 48.5 | Ref.[16] |
| CG Fagaceae | Fagaceae_*Quercus_canariensis,* Fagaceae_*Quercus_ilex* | min | 37.8 | Ref.[17] |
| CG Polygalaceae | Polygalaceae_*Muraltia_pageae,* Polygalaceae_*Polygala_hispida* | min | 56 | Ref.[18] |
| SG Cucurbitaceae | Cucurbitaceae_*Siraitia_africana,* Begoniaceae_*Begonia_dregei* | min | 47.8 | Ref.[19] |

| | | | | |
|---|---|---|---|---|
| CG Anisophylleaceae | Anisophylleaceae_*Poga_oleosa,* Anisophylleaceae_*Anisophyllea_laurina* | min | 11.63 | Ref.[20] |
| SG Celastraceae | Celastraceae_*Salacia_loloensis,* Lepidobotryaceae_*Lepidobotrys_staudtii* | min | 46.3 | Ref.[21] |
| CG Celastraceae | Celastraceae_*Parnassia_palustris,*Celastraceae_*Crossopetalum_mossambicense* | min | 37.8 | Ref.[22] |
| CG Simaroubaceae | Simaroubaceae_*Harrisonia_abyssinica,* Simaroubaceae_*Brucea_tenuifolia* | min | 41.2 | Ref.[23] |
| CG Meliaceae | Meliaceae_*Melia_azedarach,* Meliaceae_*Khaya_senegalensis* | min | 47.8 | Ref.[24] |
| SG Malvaceae | Malvaceae_*Hermannia_cuneifolia,* Thymelaeaceae_*Lachnaea_elsieae* | min | 72.1 | Ref.[25] |
| CG Vitaceae | Vitaceae_*Leea_guineensis,* Vitaceae_*Cissus_trothae* | min | 61.6 | Ref.[26] |
| SG Melastomataceae | Melastomataceae_*Lijndenia_procteri,* Penaeaceae_*Penaea_cneorum* | min | 56 | Ref.[27] |
| CG Lythraceae | Lythraceae_*Pemphis_acidula,* Lythraceae_*Ammannia_schinzii* | min | 72.1 | Ref.[28] |
| SG Combretaceae | Combretaceae_*Terminalia_ivorensis,* Onagraceae_*Ludwigia_erecta* | min | 86.3 | Ref.[29] |
| CG Francoaceae | Francoaceae_*Greyia_flanaganii,* Francoaceae_*Melianthus_comosus* | min | 15.97 | Ref.[30] |
| SG Haloragaceae | Haloragaceae_*Laurembergia_repens,* Crassulaceae_*Crassula_cordata* | min | 72.1 | Ref.[31] |
| SG Menispermaceae | Menispermaceae_*Tinospora_tenera,* Ranunculaceae_*Nigella_arvensis* | min | 89.8 | Ref.[32] |
| CG Proteaceae | Proteaceae_*Brabejum_stellatifolium,* Proteaceae_*Aulax_umbellata* | min | 72.1 | Ref.[33] |
| SG Asteraceae | Asteraceae_*Serratula_tinctoria,* Goodeniaceae_*Scaevola_taccada* | min | 47.6 | Ref.[13,34] |
| SG Zingiberaceae | Zingiberaceae_*Aframomum_thonneri,* Costaceae_*Costus_afer* | min | 66 | Ref.[35] |
| CG Typhaceae | Typhaceae_*Typha_domingensis,* Typhaceae_*Sparganium_erectum* | min | 51.66 | Ref.[36] |
| CG Restionaceae | Restionaceae_*Hypodiscus_neesii,* Flagellariaceae_*Flagellaria_indica* | min | 27.7 | Ref.[37] |
| CG Cyperaceae | Cyperaceae_*Chrysitrix_dodii,* Cyperaceae_*Scleria_foliosa* | min | 47 | Ref.[38] |
| SG Posidoniaceae | Posidoniaceae_*Posidonia_oceanica,* Cymodoceaceae_*Cymodocea_nodosa* | min | 66 | Ref.[39] |
| CG Hydrocharitaceae | Hydrocharitaceae_*Najas_marina,* Hydrocharitaceae_*Ottelia_fischeri* | min | 55.9 | Ref.[40] |

| | | | | |
|---|---|---|---|---|
| CG Hernandiaceae | Hernandiaceae_*Gyrocarpus_angustifolius,* Hernandiaceae_*Hernandia_beninensis* | min | 41.2 | Ref.[10] |
| SG *Ceratophyllum* | Ceratophyllaceae_*Ceratophyllum_demersum,* Aristolochiaceae_*Aristolochia_albida* | min | 127.2 | Ref.[41] |
| SG Cabombaceae | Cabombaceae_*Brasenia_schreberi,* Nymphaeaceae_*Nuphar_lutea* | min | 100.5 | Ref.[42] |
| SG Fagales | Myricaceae_*Morella_faya,* Begoniaceae_*Begonia_dregei* | min | 96 | Ref.[43] |
| CG Fabales | Surianaceae_*Suriana_maritima*, Fabaceae_*Prioria_msoo* | min | 59.9 | Ref.[44,45] |
| CG Sapindales | Nitrariaceae_*Nitraria_retusa,* Rutaceae_*Zanthoxylum_humile* | min | 65 | Ref.[45.46] |
| CG Myrtales | Combretaceae_*Combretum_nelsonii,* Myrtaceae_*Syzygium_pondoense* | min | 88.2 | Ref.[29,45] |
| CG Saxifragales | Hamamelidaceae_*Trichocladus_ellipticus,* Grossulariaceae_*Ribes_alpinum* | min | 89.3 | Ref.[47] |
| CG Ericales | Balsaminaceae_*Impatiens_nana*, Primulaceae_*Samolus_porosus* | min | 91.2 | Ref.[48] |
| CG Cornales | Curtisiaceae_*Curtisia_dentata,* Hydrostachyaceae_*Hydrostachys_insignis* | min | 86.3 | Ref.[49] |
| CG Arecales | Arecaceae_*Raphia_vinifera,* Arecaceae_*Borassus_aethiopum* | min | 65 | Ref.[50] |
| CG Pandanales | Pandanaceae_*Pandanus_rabaiensis,* Velloziaceae_*Xerophyta_spekei* | min | 55.8 | Ref.[51] |
| SG Magnoliales | Myristicaceae_*Coelocaryon_oxycarpum,* Lauraceae_*Cryptocarya_woodii* | min | 37.2 | Ref.[52] |
| CG Icacinaceae | Icacinaceae_*Cassinopsis_tinifolia,* Icacinaceae_*Iodes_liberica* | min | 47.8 | Ref.[53] |
| CG Menyanthaceae | Menyanthaceae_*Menyanthes_trifoliata,* Menyanthaceae_*Villarsia_capensis* | min | 5.33 | Ref.[54] |
| SG Cunoniaceae | Cunoniaceae_*Cunonia_capensis,* Connaraceae_*Connarus_rostratus* | min | 80.7 | Ref.[55] |
| SG Ctenolophonaceae | Ctenolophonaceae_*Ctenolophon_englerianus,* Humiriaceae_*Sacoglottis_gabonensis* | min | 66 | Ref.[56] |
| SG Clusiaceae | Clusiaceae_*Garcinia_kola,* Podostemaceae_*Tristicha_trifaria* | min | 86.3 | Ref.[57] |
| CG Brassicaceae | Brassicaceae_*Aethionema_thomasianum,* Brassicaceae_*Ammosperma_cinereum* | min | 23.03 | Ref.[10,58,59] |
| SG Brassicales | Brassicaceae_*Brassica_nigra,* Neuradaceae_*Neuradopsis_austroafricana* | min | 89.3 | Ref.[13,60] |
| CG Zingiberales | Strelitziaceae_*Strelitzia_alba,* Musaceae_*Ensete_homblei* | min | 83.5 | Ref.[45,61] |

| | | | | |
|---|---|---|---|---|
| CG Rhamnaceae | Rhamnaceae_Ziziphus_rivularis, Rhamnaceae_Rhamnus_pumila | min | 48.6 | Ref.[12,13] |
| CG Burseraceae | Burseraceae_Commiphora_edulis, Burseraceae_Boswellia_rivae | min | 48.6 | Ref. [13,24,62] |
| CG Bignoniaceae | Bignoniaceae_Podranea_ricasoliana, Bignoniaceae_Rhigozum_obovatum | min | 38.8 | Ref.[13,63] |
| CG Gentianaceae | Gentianaceae_Faroa_axillaris, Gentianaceae_Exochaenium_debile | min | 33.9 | Ref.[13] |
| SG Annonaceae | Annonaceae_Meiocarpidium_oliverianum, Myristicaceae_Brochoneura_usambarensis | min | 87.5 | Ref.[13,64] |
| SG Bignoniaceae | Bignoniaceae_Kigelia_africana, Verbenaceae_Lantana_rugosa | min | 49.4 | Ref.[45,65] |
| CG Droseraceae | Droseraceae_Drosera_natalensis, Droseraceae_Drosophyllum_lusitanicum | min | 23 | Ref.[66–69] |
| SG Dipterocarpaceae | Dipterocarpaceae_Monotes_glaber, Cistaceae_Fumana_fontqueri | min | 56 | Ref.[69,70] |
| SG Ulmaceae | Ulmaceae_Ulmus_minor, Cannabaceae_Celtis_gomphophylla | max | 88.5 | Ref.[13] |
| CG Primulaceae | Primulaceae_Embelia_schimperi, Primulaceae_Maesa_lanceolata | min | 66 | Ref.[70,71] |
| SG Monocots | Araceae_Anchomanes_difformis, Nymphaeaceae_Nymphaea_thermarum | min | 113 | Ref.[13,72,73] |
| CG Caryophyllaceae | Caryophyllaceae_Telephium_imperati, Caryophyllaceae_Silene_berthelotiana | min | 33.9 | Ref.[13,74] |

**Table S2**. **Environmental, anthropogenic, and paleoenvironmental variables used in this study.** This table summarizes the factors included in our analyses, along with the rationale for their selection. Modern environmental variables (e.g., temperature, precipitation, elevation) capture current ecological conditions affecting plant diversity, while anthropogenic variables (e.g., human footprint) reflect human impacts on biodiversity. Paleoenvironmental proxies (paleotemperature, atmospheric $CO_2$, paleo-elevation) were included to investigate historical drivers of diversification and macroevolutionary patterns over geological time. Units, sources, and references are provided for each variable.

| Variable | Source | Biological importance |
|---|---|---|
| Mean annual temperature (°C) | CHELSA v.1.2 | Influences physiological processes, growth rates, and survival; affects species distributions and community composition. |
| Annual precipitation (mm) | CHELSA v.1.2 | Determines water availability; shapes vegetation types and drives patterns of richness and endemism. |

| | | |
|---|---|---|
| Precipitation seasonality | CHELSA v.1.2 | Seasonal variation in rainfall influences plant phenology, life history strategies, and the resilience of species to drought or flooding. |
| Average elevation (m) | SRTM30) | Influences climate, temperature, and precipitation; promotes environmental heterogeneity, endemism, and local diversification. |
| Number of soil types | SoilGrids1km89[1 km] | Soil diversity drives niche differentiation, supporting higher species richness and phylogenetic diversity. |
| Topographic roughness | SRTM30 | Creates microhabitats and ecological niches, facilitating coexistence and local adaptation. |
| Aridity index | Global-AI_PET_v3: https://doi.org/10.6084/m9.figshare.7504448.v5 | Quantifies water stress; explains patterns of plant adaptation, richness, and endemism in arid vs humid regions. |
| Human footprint | Global Human foot print https://doi.org/10.5061/dryad.052q5 | Measures human impact (urbanization, agriculture, infrastructure) on natural habitats. High human footprint can lead to habitat loss, fragmentation, and species declines. |
| Paleo temperature (°C) | Condamine et al., 2020 | Historical temperatures shape habitat suitability, range shifts, and the tempo of evolution and diversification. |
| Paleo mean elevation (m) | Scotese and Wright, 2018 | Historical topography likely affects habitat availability, speciation opportunities, and extinction risk over evolutionary timescales. |
| Paleo atmospheric $CO_2$ (ppm) | Condamine et al., 2020 | $CO_2$ concentrations influence plant growth rates, photosynthesis, and global vegetation structure, affecting diversification dynamics. |

**Table S3. Results of paleoenvironment-dependent diversification models for African vascular plant genera estimated with RPANDA 1.9.** Rates of speciation (λ) and extinction (μ) were modeled as functions of paleotemperature (Temp), atmospheric $CO_2$ ($CO_2$), and past elevation (Elev). For each environmental variable, four hierarchical models were fitted, allowing speciation and extinction rates to be constant (CST), vary through time (TimeVar), or vary as functions of the environmental variable (TempVar, $CO_2$Var, ElevVar). Model fit was evaluated using the corrected Akaike Information Criterion (AICc), the difference in AICc relative to the best-fitting model (ΔAIC), and Akaike weight (ω). The model with the lowest AICc was considered the best fit. Parameter estimates include $\lambda_0$ (baseline speciation rate), $\mu_0$ (baseline extinction rate), α (strength of the effect of the environmental variable on diversification), and β (baseline rate when there is no environmental effect). The best and second best model are highlighted in bold.

| **Models** | **NP** | **LogL** | **AICc** | **λ0** | **α** | **μ0** | **β** | **Akaike_w** |
|---|---|---|---|---|---|---|---|---|
| BCSTDCST | 2 | -15755.015 | 31514.033 | 0.063 | - | 0.045 | - | 0 |
| BCO2VarDCST_EXPO | 3 | -15748.553 | 31503.113 | 0.07 | 0.0903 | -0.055 | - | 0 |
| BCSTDCO2Var_EXPO | 3 | -15746.399 | 31498.805 | 0.067 | - | 0.045 | -0.1822 | 0 |
| BCO2VarDCO2Var_EXPO | 4 | -15731.909 | 31471.83 | 0.054 | -0.5361 | 0.019 | -2.3232 | 0 |
| BCO2VarDCST_LIN | 3 | -15748.215 | 31502.436 | 0.071 | 0.0071 | 0.056 | - | 0 |
| BCSTDCO2Var_LIN | 3 | -15747.734 | 31501.474 | 0.065 | - | 0.049 | -0.0071 | 0 |
| BCO2VarDCO2Var_LIN | 4 | -15747.692 | 31503.395 | 0.067 | 0.0015 | 0.051 | -0.0057 | 0 |
| BTempVarDCST_EXPO | 3 | -15754.506 | 31515.02 | 0.07 | 0.0646 | 0.048 | - | 0 |
| BCSTDTempVar_EXPO | 3 | -15752.233 | 31510.473 | 0.067 | - | 0.05 | -0.3415 | 0 |
| BTempVarDTempVar_EXPO | 4 | -15752.04 | 31512.092 | 0.064 | -0.0488 | 0.043 | -0.4284 | 0 |

| | | | | | | | | |
|---|---|---|---|---|---|---|---|---|
| BTempVarDCST_LIN | 3 | --15754.473 | 31514.954 | 0.066 | 0.0045 | 0.048 | - | 0 |
| BCSTDTempVar_LIN | 3 | -15753.022 | 31512.05 | 0.065 | - | 0.045 | -0.0116 | 0 |
| BTempVarDTempVar_LIN | 4 | -15753.021 | 31514.052 | 0.065 | -3.00E-04 | 0.045 | -0.0119 | 0 |
| BElevVarDCST_EXPO | 3 | -15728.129 | 31462.266 | 0.099 | -0.3901 | 0.065 | - | 0 |
| BCSTDElevVar_EXPO | 3 | -15738.859 | 31483.725 | 0.069 | - | 0.037 | 0.5621 | 0 |
| **BElevVarDElevVar_EXPO** | **4** | **-15699.692** | **31407.396** | **0.343** | **-1.1274** | **0.317** | **-1.0885** | **0.985** |
| BElevVarDCST_LIN | 3 | -15727.144 | 31460.294 | 0.101 | -0.0339 | 0.067 | - | 0 |
| BCSTDElevVar_LIN | 3 | -15740.736 | 31487.478 | 0.068 | - | 0.037 | 0.0244 | 0 |
| **BElevVarDElevVar_LIN** | **4** | **-15703.897** | **31415.806** | **0.341** | **-0.1846** | **0.303** | **-0.1524** | **0.015** |
| BTimeVarDCST_EXPO | 3 | -15754.924 | 31515.854 | 0.062 | -1.00E-04 | 0.044 | - | 0 |
| BCSTDTimeVar_EXPO | 3 | -15754.903 | 31515.813 | 0.062 | - | 0.044 | 2e-04 | 0 |
| BTimeVarDTimeVar_EXPO | 4 | -15753.268 | 31514.546 | 0.063 | 0.0037 | 0.049 | 0.0044 | 0 |
| BTimeVarDCST_LIN | 3 | -15754.96 | 31515.926 | 0.062 | 0 | 0.044 | - | 0 |
| BCSTDTimeVar_LIN | 3 | -15754.96 | 31515.927 | 0.062 | - | 0.044 | 0 | 0 |
| BTimeVarDTimeVar_LIN | 4 | -15754.968 | 31517.947 | 0.062 | 0 | 0.045 | 0 | 0 |

**Table S4. Results of paleoenvironment-dependent diversification models for African Proteaceae (*Protea* clade) estimated using RPANDA 1.9.** Rates of speciation ($\lambda$) and extinction ($\mu$) were modeled as functions of paleotemperature (Temp), atmospheric $CO_2$ ($CO_2$), and past elevation (Elev). For each environmental variable, four hierarchical models were fitted, allowing speciation and extinction rates to be constant (CST), vary through time (TimeVar), or vary as functions of the environmental variable (TempVar, $CO_2$Var, ElevVar). Model fit was evaluated using the corrected Akaike Information Criterion (AICc), the difference in AICc relative to the best-fitting model ($\Delta$AIC), and Akaike weight ($\omega$). The model with the lowest AICc was considered the best fit. Parameter estimates include $\lambda_0$ (baseline speciation rate), $\mu_0$ (baseline extinction rate), $\alpha$ (strength of the effect of the environmental variable on diversification), and $\beta$ (baseline rate when there is no environmental effect). The best and second best model are highlighted in bold.

| **Models** | **Parameters** | **logL** | **AICc** | **Lambda** | **Alpha** | **Mu** | **Beta** | **Akaike_w** |
|---|---|---|---|---|---|---|---|---|
| BCSTDCST | 2 | -116.228 | 236.832 | 0.092 | - | 0 | - | 0.021 |
| BCO2VarDCST_EXPO | 3 | -115.959 | 238.691 | 0.189 | 1.4166 | 0 | - | 0.008 |
| BCSTDCO2Var_EXPO | 3 | -116.234 | 239.242 | 0.092 | - | 0 | 0.0151 | 0.006 |
| BCO2VarDCO2Var_EXPO | 4 | -115.959 | 241.251 | 0.189 | 1.4164 | 0 | 0.0861 | 0.002 |
| BCO2VarDCST_LIN | 3 | -115.733 | 238.241 | 0.226 | 0.2604 | 0 | - | 0.011 |
| BCSTDCO2Var_LIN | 3 | -116.234 | 239.243 | 0.092 | - | 0 | 0 | 0.006 |
| BCO2VarDCO2Var_LIN | 4 | -115.705 | 240.743 | 0.286 | 0.3627 | -0.167 | -0.3217 | 0.003 |
| BTempVarDCST_EXPO | 3 | -114.281 | 235.336 | 0.286 | 2.1294 | 0 | - | 0.045 |
| BCSTDTempVar_EXPO | 3 | -116.234 | 239.242 | 0.092 | - | 0 | 0.0151 | 0.006 |
| BTempVarDTempVar_EXPO | 4 | -114.281 | 237.896 | 0.286 | 2.1279 | 0 | 0.1225 | 0.013 |
| BTempVarDCST_LIN | 3 | -112.32 | 231.415 | 0.418 | 0.4831 | 0.099 | - | 0.322 |

| | | | | | | | | |
|---|---|---|---|---|---|---|---|---|
| BCSTDTempVar_LIN | 3 | -116.234 | 239.242 | 0.092 | - | 0 | 0 | 0.006 |
| **BTempVarDTempVar_LIN** | **4** | **-111.351** | **232.036** | **0.454** | **0.5389** | **-0.315** | **-0.3815** | **0.236** |
| BTimeVarDCST_EXPO | 3 | -115.875 | 238.524 | 1.202 | -1.8507 | 0 | - | 0.009 |
| BCSTDTimeVar_EXPO | 3 | -116.234 | 239.242 | 0.092 | - | 0 | 0.0061 | 0.006 |
| BTimeVarDTimeVar_EXPO | 4 | -115.875 | 241.083 | 1.202 | -1.8504 | 0 | 0.0988 | 0.003 |
| BTimeVarDCST_LIN | 3 | -115.857 | 238.488 | 0.357 | -0.1903 | 0 | - | 0.009 |
| BCSTDTimeVar_LIN | 3 | -116.234 | 239.242 | 0.092 | | 0 | 0 | 0.006 |
| BTimeVarDTimeVar_LIN | 4 | -115.835 | 241.003 | 0.744 | -0.4591 | -0.714 | 0.4936 | 0.003 |
| BElevVarDCST_EXPO | 3 | -115.355 | 237.484 | 0.069 | 0.0296 | 0 | - | 0.015 |
| BCST_DElevVar_EXPO | 3 | -116.228 | 239.231 | 0.092 | - | 0 | 0.0139 | 0.006 |
| BElevVar_DElevVar_EXPO | 4 | -115.355 | 240.043 | 0.069 | 0.0296 | 0 | 0.0367 | 0.004 |
| BElevVarDCST_LIN | 3 | -115.073 | 236.92 | 0.055 | 0.004 | 0 | - | 0.021 |
| BCST_DElevVar_LIN | 3 | -116.234 | 239.242 | 0.092 | - | 0 | 0 | 0.006 |
| **BElevVar_DElevVar_LIN** | **4** | **-111.407** | **232.147** | **0.035** | **0.0414** | **0.022** | **0.0359** | **0.223** |

**Table S5. Results of paleoenvironment-dependent diversification models for the African rain forest clade Monodoreae (Annonaceae) estimated using RPANDA 1.9.** Speciation (λ) and extinction (μ) rates were modeled as functions of paleotemperature (Temp), atmospheric $CO_2$ concentration ($CO_2$), and past elevation (Elev). The Monodoreae phylogeny corresponds to the clade previously analyzed by Dagallier et al. (*17*) and is used here as a case study to evaluate paleoenvironment-dependent diversification patterns under a unified modeling framework. For each environmental variable, four hierarchical models were fitted, allowing diversification rates to be constant (CST), vary through time (TimeVar), or vary as functions of the focal environmental variable (TempVar, $CO_2$Var, ElevVar). Model fit was assessed using the corrected Akaike Information Criterion (AICc), ΔAIC relative to the best-fitting model, and Akaike weight (ω).

Parameter estimates include $\lambda_0$ (baseline speciation rate), $\mu_0$ (baseline extinction rate), $\alpha$ (strength of environmental dependence), and $\beta$ (baseline rate in the absence of environmental effects). The best- and second-best-supported models are highlighted in bold.

| **Models** | **Parameters** | **logL** | **AICc** | **Lambda** | **Alpha** | **Mu** | **Beta** | **Akaike_w** |
|---|---|---|---|---|---|---|---|---|
| BCSTDCST | 2 | -113.959 | 232.225 | 0.241 | - | 0.042 | - | 0.004 |
| BCO2VarDCST_EXPO | 3 | -113.425 | 233.482 | 2.082 | 3.8468 | 0.089 | - | 0.002 |
| BCSTDCO2Var_EXPO | 3 | -113.666 | 233.963 | 0.262 | - | 31.634 | 11.4936 | 0.002 |
| BCO2VarDCO2Var_EXPO | 4 | -111.721 | 232.523 | 107.023 | 10.8075 | 3932.347 | 18.7358 | 0.004 |
| BCO2VarDCST_LIN | 3 | -113.252 | 233.136 | 1.036 | 1.4182 | 0.101 | - | 0.003 |
| BCSTDCO2Var_LIN | 3 | -113.153 | 232.938 | 0.269 | - | 1.567 | 2.8522 | 0.003 |
| BCO2VarDCO2Var_LIN | 4 | -111.304 | 231.689 | 1.979 | 3.0687 | 3.204 | 5.7865 | 0.005 |
| BTempVarDCST_EXPO | 3 | -113.858 | 234.347 | 0.153 | -0.5448 | 0 | - | 0.001 |
| BCSTDTempVar_EXPO | 3 | -112.851 | 232.333 | 0.25 | - | 2.212 | 6.336 | 0.004 |
| BTempVarDTempVar_EXPO | 4 | -111.743 | 232.566 | 2.313 | 2.8253 | 5.657 | 4.9749 | 0.003 |
| BTempVarDCST_LIN | 3 | -113.76 | 234.152 | 0.104 | -0.1791 | 0 | - | 0.002 |
| BCSTDTempVar_LIN | 3 | -112.659 | 231.949 | 0.267 | - | 0.409 | 0.5242 | 0.005 |
| **BTempVarDTempVar_LIN** | **4** | **-106.653** | **222.386** | **3.383** | **3.8819** | **3.365** | **3.8776** | **0.558** |
| BTimeVarDCST_EXPO | 3 | -112.648 | 231.927 | 0 | 6.2943 | 0 | - | 0.005 |
| BCSTDTimeVar_EXPO | 3 | -113.541 | 233.714 | 0.265 | - | 249.803 | -5.6508 | 0.002 |
| BTimeVarDTimeVar_EXPO | 4 | -112.648 | 234.376 | 0 | 6.2999 | 254.805 | -17.0777 | 0.001 |
| BTimeVarDCST_LIN | 3 | -112.736 | 232.104 | -1.098 | 0.9235 | 0 | - | 0.004 |
| BCSTDTimeVar_LIN | 3 | -112.78 | 232.191 | 0.238 | - | 2.936 | -2.0269 | 0.004 |

| | | | | | | | | |
|---|---|---|---|---|---|---|---|---|
| BTimeVarDTimeVar_LIN | 4 | -112.734 | 234.549 | -0.839 | 0.7451 | -0.77 | 0.5313 | 0.001 |
| BElevVarDCST_EXPO | 3 | -113.154 | 232.939 | 0.269 | -0.0398 | 0 | - | 0.003 |
| BCST_DElevVar_EXPO | 3 | -112.804 | 232.239 | 0.246 | - | 0.012 | 0.1923 | 0.004 |
| BElevVar_DElevVar_EXPO | 4 | -112.801 | 234.684 | 0.244 | 0.0057 | 0.015 | 0.1832 | 0.001 |
| BElevVarDCST_LIN | 3 | -112.98 | 232.592 | 0.27 | -0.009 | 0 | - | 0.003 |
| BCST_DElevVar_LIN | 3 | -112.538 | 231.708 | 0.256 | - | 0.03 | -0.0171 | 0.005 |
| **BElevVar_DElevVar_LIN** | **4** | **-107.064** | **223.209** | **0.116** | **0.2831** | **0.071** | **-0.2822** | **0.37** |

**Table S6.** Model fit comparisons for BiSSE analyses. For each model, we denote the number of parameters (NP), the log-likelihood (logLik), the Akaike Information Criterion for sample size (AIC), chi-square, p-value (Pr > ChiSq), and the model parameter values: speciation (), extinction () and transitions (q) rates (in $Myr^{-1}$) for states 0 and 1. The best-fitting model for each trait is highlighted in bold, identified with the lowest AIC.

| States | Model.parameter | NP | logLik | AIC | ChiSq | Pr_chiSq | l0 | l1 | m0 | m1 | q01 | q10 |
|---|---|---|---|---|---|---|---|---|---|---|---|---|
| **Habitat** | full | 6 | -32684.01 | 65580.01 | - | - | 0.65 | 0.84 | 0.63 | 0.82 | 0.02 | 0.01 |
| state 0 (tropical) | mu.q free | 4 | -32709.5 | 65427 | 50.99 | 0 | 0.75 | 0.78 | - | 0.76 | - | 0.01 |
| | Lambda.mu free | 4 | -32701.12 | 65410.23 | 34.22 | 0 | - | 0.77 | - | 0.75 | 0.02 | 0.01 |
| state 1 (non-tropical) | Lambda.q free | 4 | -32710.95 | 65429.9 | 53.89 | 0 | - | 0.77 | 0.77 | 0.75 | - | 0.01 |
| | **Lambda free** | **5** | **-32697.3** | **65404.61** | **26.59** | **0** | **-** | **0.77** | **0.74** | **0.76** | **0.03** | **0.01** |
| **Life form** | full | 6 | -26369.73 | 52751.45 | - | - | 0.45 | 0.79 | 0.43 | 0.77 | 0 | 0 |

| | | | | | | | | | | | | |
|---|---|---|---|---|---|---|---|---|---|---|---|---|
| state 0 (woody) | mu.q free | 4 | -26466.33 | 52940.66 | 193.21 | 0 | 0.57 | 0.57 | - | 0.56 | - | 0 |
| | Lambda.mu free | 4 | -26462.53 | 52933.06 | 185.6 | 0 | - | 0.58 | - | 0.56 | 0 | 0 |
| state 1 (herb) | Lambda.q free | 4 | -26468.49 | 52944.99 | 197.53 | 0 | - | 0.58 | 0.56 | 0.56 | - | 0 |
| | **Lambda free** | **5** | **-26456.77** | **52923.54** | **174.09** | **0** | **-** | **0.58** | **0.57** | **0.56** | **0** | **0** |
| **Succulence** | full | 6 | -44300.25 | 88612.5 | - | - | 3.05 | 0.89 | 3.03 | 0.88 | 0.02 | 0 |
| state 0 (non-succulent) | mu.q free | 4 | -44691.87 | 89391.74 | 783.25 | 0 | 0.93 | 0.95 | - | 0.93 | - | 0 |
| | Lambda.mu free | 4 | -44492.04 | 88992.08 | 383.58 | 0 | - | 0.94 | - | 0.93 | 0.02 | 0 |
| state 1 (succulent) | Lambda.q free | 4 | -44686.04 | 89380.07 | 771.58 | 0 | - | 0.94 | 0.95 | 0.93 | - | 0 |
| | **Lambda free** | **5** | **-44462.37** | **88934.73** | **324.24** | **0** | **-** | **0.95** | **0.88** | **0.94** | **0.06** | **0** |
| **Aridity** | full | 6 | -15135.72 | 30283.44 | - | - | 0.86 | 0.4 | 0.85 | 0.38 | 0 | 0.01 |
| state 0 (dry) | mu.q free | 4 | -15207.61 | 30423.22 | 143.79 | 0 | 0.54 | 0.54 | - | 0.53 | - | 0 |
| | **Lambda.mu free** | **4** | **-15205.71** | **30419.42** | **139.98** | **0** | **-** | **0.54** | **-** | **0.52** | **0.01** | **0** |
| state 1 (humid) | Lambda.q free | 4 | -15206.3 | 30420.6 | 141.16 | 0 | - | 0.54 | 0.53 | 0.53 | - | 0 |
| | Lambda free | 5 | -15205.63 | 30421.27 | 139.83 | 0 | - | 0.54 | 0.53 | 0.52 | 0.01 | 0 |

### Supplementary References


1. N. F. Hughes, A. B. McDougall, Records of angiospermid pollen entry into the English Early Cretaceous succession. *Rev. Palaeobot. Palyno.* **50**, 255–272 (1987).
2. N. F. Hughes, A. B. McDougall, J. L. Chapman, Exceptional new record of Cretaceous Hauterivian angiospermid pollen from Southern England. *J. Micropalaeontol.* **10**, 75–82 (1991).
3. G. J. Brenner, Flowering Plant Origin, Evolution, and Phylogeny (Chapman and Hall, New York, 1996).
4. S. Magallón, K. W. Hilu, D. Quandt, Land plant evolutionary timeline: Gene effects are secondary to fossil constraints in relaxed clock estimation of age and substitution rates. *Am. J. Bot.* **100**, 556–573 (2013).
5. Li, H.-T., Yi, T.-S., Gao, L.-M., Ma, P.-F., Zhang, T., Yang, J.-B., Gitzendanner, M. A., Fritsch, P. W., Cai, J., Luo, Y., Wang, H., van der Bank, M., Berndt, R., Wang, Q.-F., Wang, J., Zhang, Z.-R., Fu, C.-N., Yang, J., Hollingsworth, M. L., Soltis, D. E. Origin of angiosperms and the puzzle of the Jurassic gap. *Nature Plants*, **5**, 461–470 (2019).
6. Wilf, P., Donovan, M. P., Cúneo, N. R., & Gandolfo, M. A. (2017). The fossil flip-leaves (Retrophyllum, Podocarpaceae) of southern South America. American Journal of Botany, 104(9), 1344–1369.
7. M. Salard-Cheboldaeff, Quelques grains de pollen peripores Tertiaires du Cameroun. *Rev. Micropaleontol*. **17**, 182–190 (1975).
8. R. C. Mehrotra, Study of plant megafossils from the Tura Formation of Nangwalbibra, Garo Hills, Meghalaya, India. *Palaeobotanist* **49**, 255–237 (2000).
9. C. A. Chmura, Upper Cretaceous (Campanian-Maastrichtian) angiosperm pollen from the western San Joaquin Valley, California, U.S.A. *Palaeontographica Abteilung B* **141**, 89–171 (1973).
10. S. R. Manchester, E. L. O'Leary, Phylogenetic distribution and identification of fin-winged fruits. *Bot. Rev.* **76**, 1–82 (2010).
11. L. Calvillo-Canadell, S. R. S. Cevallos-Ferriz, Reproductive structures of Rhamnaceae from the Cerro del Pueblo (Late Cretaceous, Coahuila) and Coatzingo (Oligocene, Puebla) Formations, Mexico. *Am. J. Bot.* **94**, 1658–1669 (2007).
12. S. R. Manchester, Biogeographical relationships of North American Tertiary floras. *Ann. Missouri Bot. Gard.* **86**, 472–522 (1999).
13. Magallón, S., Gómez-Acevedo, S., Sánchez-Reyes, L. L., & Hernández-Hernández, T. (2015). A metacalibrated time-tree documents the early rise of flowering plant phylogenetic diversity. New Phytologist, 207(2), 437–453.
14. M. E. J. Chandler, The Lower Tertiary floras of Southern England. III. Flora of the Bournemouth Beds; the Boscombe, and the Highcliff Sands. (British Museum, London, 1963).
15. J. H. Germeraad, C. A. Hopping, J. Muller, Palynology of Tertiary sediments from tropical areas. *Rev. Palaeobot. Palyno.* **6**, 189–348 (1968).

16. R. W. Brown, Additions to the flora of the Green River formation. *U.S. Geol. Surv. Prof. Pap.* **154**, 279–292 (1929).
17. W. L. Crepet, C. P. Daghlian, Castaneoid inflorescences from the Middle Eocene of Tennessee and the diagnostic value of pollen (at the subfamily level) in Fagaceae. *Am. J. Bot.* **67**, 739–757 (1980).
18. K. B. Pigg, M. L. DeVore, M. F. Wojciechowski, Paleosecuridaca curtisii gen. et sp. nov., Securidaca-like samaras (Polygalaceae) from the Late Paleocene of North Dakota and their significance to the divergence of families within the Fabales. *Int. J. Plant Sci.* **169**, 1304–1313 (2008).
19. M. E. J. Chandler, The Lower Tertiary floras of southern England. I. Paleocene floras. London Clay Flora (Supplement). Text and Atlas. (British Museum, London, 1961).
20. J. A. R. Anderson, J. Muller, Palynological study of a Holocene peat and a Miocene coal deposit from NW Borneo. *Rev. Palaeobot. Palyno.* **19**, 291–351 (1975).
21. R. C. Mehrotra, U. Prakash, M. B. Bande, Fossil woods of Lophopetalum and Artocarpus from the Deccan Intertrappean Beds of Mandla district, Madhya Pradesh, India. *Palaeobotanist* **32**, 310–320 (1984).
22. Hollick, The Tertiary floras of Alaska. *U.S. Geol. Surv. Prof. Pap.* **182**, 1–185 (1936).
23. M. E. Collinson, S. R. Manchester, V. Wilde, *Fossil fruits and seeds of the Middle Eocene Messel biota, Germany*. *Abh. Senckenberg Ges. Naturforsch.* **570**, 1–251 (2012).
24. E. M. Reid, M. E. J. Chandler, The London Clay flora. (British Museum, London, 1933).
25. E. Estrada-Ruiz, H. I. Martínez-Cabrera, S. R. S. Cevallos-Ferriz, Fossil woods from the late Campanian–early Maastrichtian Olmos Formation, Coahuila, Mexico. *Rev. Palaeobot. Palyno.* **145**, 123–133 (2007).
26. S. R. Manchester, D. K. Kapgate, J. Wen, Oldest fruits of the grape family (Vitaceae) from the Late Cretaceous Deccan Cherts of India. *Am. J. Bot.* **100**, 1849–1859 (2013).
27. R. W. Brown, Paleocene flora of the Rocky Mountains and Great Plains. *Geol. Surv. Prof. Pap.* **375**, 1–119 (1962).
28. E. Estrada-Ruiz, L. Calvillo-Canadell, S. R. S. Cevallos-Ferriz, Upper Cretaceous aquatic plants from Northern Mexico. *Aquat. Bot.* **90**, 282–288 (2009).
29. M. Takahashi, P. R. Crane, H. Ando, Esgueiria futabensis sp. nov., a new angiosperm flower from the Upper Cretaceous (Lower Coniacian) of northeastern Honshu, Japan. *Paleontol. Res.* **3**, 81–87 (1999).
30. L. Palazzesi, M. Gottschling, V. Barreda, M. Weigend, First Miocene fossils of Vivianiaceae shed new light on phylogeny, divergence times, and historical biogeography of Geraniales. *Biol. J. Linn. Soc.* **107**, 67–85 (2012).
31. G. R. Hernandez-Castillo, S. R. S. Cevallos-Ferriz, Reproductive and vegetative organs with affinities to Haloragaceae from the Upper Cretaceous Huepac Chert Locality of Sonora, Mexico. *Am. J. Bot.* **86**, 1717 (1999).
32. E. Knobloch, D. H. Mai, Neue Gattungen nach Früchten und Samen aus dem Cenoman bis Maastricht (Kreide) von Mitteleuropa. *Feddes Repert.* **95**, 3–41 (1984).

33. R. J. Carpenter, M. K. Macphail, G. J. Jordan, R. S. Hill, Fossil evidence for open, Proteaceae-dominated heathlands and fire in the late Cretaceous of Australia. *Am. J. Bot.* **102**, 2092–2107 (2015).
34. V. D. Barreda, L. Palazzesi, L. Katinas et al., An extinct Eocene taxon of the daisy family (Asteraceae): Evolutionary, ecological and biogeographical implications. *Ann. Bot*. **109**, 127–134 (2012).
35. F. Knowlton, Fossil floras of the Vermejo and Raton formations of Colorado and New Mexico. *U.S. Geol. Surv. Prof. Pap.* **101**, 223–435 (1917).
36. L. Grande, Paleontology of the Green River Formation, with a review of the fish fauna, second edition. Geol. Surv. Wyoming Bull. 63, 1–333 (1984).
37. M. E. Dettmann, H. T. Clifford, Monocotyledon fruits and seeds, and an associated palynoflora from Eocene–Oligocene sediments of coastal central Queensland, Australia. *Rev. Palaeobot. Palyno.* **110**, 141–173 (2000).
38. S. Y. Smith, M. E. Collinson, D. A. Simpson, P. J. Rudall, F. Marone, M. Stampanoni, Elucidating the affinities and habitat of ancient, widespread Cyperaceae: Volkeria messelensis gen. et sp. nov., a fossil mapanioid sedge from the Eocene of Europe. *Am. J. Bot*. **96**, 1506–1518 (2009).
39. R. W. J. M. van der Ham, J. H. A. van Konijnenburg-van Cittert, L. Indeherberge, Seagrass foliage from the Maastrichtian type area (Maastrichtian, Danian, NE Belgium, SE Netherlands). *Rev. Palaeobot. Palyno*. **144**, 301–321 (2007).
40. D. Bone, The stratigraphy of the Reading Beds (Palaeocene), at Felpham, West Sussex. *Tertiary Res*. **8**, 17–32 (1986).
41. B. Gomez, V. Daviero-Gomez, C. Coiffard, C. Martín-Closas, D. L. Dilcher, Montsechia, an ancient aquatic angiosperm. *Proc. Natl. Acad. Sci. USA* **112**, 10985–10988 (2015).
42. D. W. Taylor, G. J. Brenner, S. H. Basha, Scutifolium jordanicum gen. et sp. nov. (Cabombaceae), an aquatic fossil plant from the Lower Cretaceous of Jordan, and the relationships of related leaf fossils to living genera. *Am. J. Bot*. **95**, 340–352 (2008).
43. E. M. Friis, K. R. Pedersen, J. Schönenberger, Normapolles plants: a prominent component of the Cretaceous rosid diversification. *Plant Syst. Evol*. **260**, 107–140 (2006).
44. P. S. Herendeen, P. R. Crane, in Advances in Legume Systematics, part 4. The fossil record, P. S. Herendeen, D. L. Dilcher, Eds. (Royal Botanic Gardens, Kew, 1992), pp. 57–68.
45. C. D. Bell, D. E. Soltis, P. S. Soltis, The age and diversification of the angiosperms re-revisited. *Am. J. Bot*. **97**, 1296–1303 (2010).
46. E. D. Knobloch, D. H. Mai, Monograph of the fruits and seeds in the Cretaceous of Central Europe. *Rozpr. Ústř. Úst. Geol*. **47**, 1–219 (1986).
47. E. J. Hermsen et al., *Divisestylus* gen. nov. (aff. Iteaceae), a fossil saxifrage from the Late Cretaceous of New Jersey, USA. *Am. J. Bot*. **90**, 1373–1388 (2003).
48. K. C. Nixon, W. L. Crepet, Late Cretaceous fossil flowers of ericalean affinity. *Am. J. Bot.* **80**, 616–623 (1993).

49. M. Takahashi, P. R. Crane, S. R. Manchester, Hironoia fusiformis gen. et sp. nov.; a cornalean fruit from the Kamikitaba locality (Upper Cretaceous, Lower Coniacian) in northeastern Japan. *J. Plant Res*. **115**, 463–473 (2002).
50. D. Pan, B. F. Jacobs, J. Dransfield, W. J. Baker, The fossil history of palms (Arecaceae) in Africa and new records from the Late Oligocene (28–27 Mya) of north-western Ethiopia. *Bot. J. Linn. Soc*. **151**, 69–81 (2006).
51. D. M. Jarzen, The terrestrial palynoflora from the Cretaceous-Tertiary transition, Alabama, U.S.A. *Pollen Spores* **20**, 535–553 (1978).
52. P. R. Crane, E. M. Friis, K. R. Pedersen, Palaeobotanical evidence on the early radiation of magnoliid angiosperms. *Plant Syst. Evol*. **8**, 51–72 (1994).
53. C. Del Rio, T. Haevermans, D. De Franceschi, First record of an Icacinaceae Miers fossil flower from Le Quesnoy (Ypresian, France) amber. *Sci. Rep*. **7**, 1–8 (2017).
54. M. Lancucka-Srodoniowa, Macroscopical plant remains from the freshwater Miocene of the Nowy Sacz Basin (West Carpathians, Poland). *Acta Palaeobot*. **20**, 3–117 (1979).
55. J. Schönenberger, E. M. Friis, M. L. Matthews, P. K. Endress, Cunoniaceae in the Cretaceous of Europe: Evidence from fossil flowers. *Ann. Bot*. **88**, 423–437 (2001).
56. P. van Hoeken-Klinkenberg, A palynological investigation of some Upper Cretaceous sediments in Nigeria. *Pollen Spores* **6**, 209–231 (1964).
57. W. L. Crepet, K. C. Nixon, Fossil Clusiaceae from the Late Cretaceous (Turonian) of New Jersey and implications regarding the history of bee pollination. *Am. J. Bot*. **85**, 1122–1133 (1998).
58. H. F. Becker, Oligocene plants from the Upper Ruby River Basin, southwestern Montana. (Geological Society of America, New York, 1961).
59. M. A. Beilstein, N. S. Nagalingum, M. D. Clements et al., Dated molecular phylogenies indicate a Miocene origin for Arabidopsis thaliana. *Proc. Natl. Acad. Sci. USA* **107**, 1872–1877 (2010).
60. M. A. Gandolfo, K. C. Nixon, W. L. Crepet, Tylerianthus crossmanensis gen. et sp. nov. (aff. Hydrangeaceae) from the Upper Cretaceous of New Jersey. *Am. J. Bot*. **85**, 376–386 (1998).
61. E. M. Friis, Spirematospermum chandlerae sp. nov., an extinct species of Zingiberaceae from the North American Cretaceous. *Tert. Res*. **9**, 7–12 (1988).
62. M. E. Collinson, Fossil Plants of the London Clay I: Field Guide to Fossils. (Palaeontological Association, London, 1983).
63. H. W. Meyer, S. R. Manchester, Oligocene Bridge Creek Flora of the John Day Formation, Oregon. (University of California Press, Berkeley, 1997).
64. Takahashi, M., Friis, E. M., Uesugi, K., Suzuki, Y., & Crane, P. R. (2008). Floral evidence of Annonaceae from the Late Cretaceous of Japan. International Journal of Plant Sciences, 169(7), 908–917.
65. C. C. Wehr, D. Q. Hopkins, The Eocene orchards and gardens of Republic, Washington. *Wash. Geol*. **22**, 27–33 (1994).

66. M. E. J. Chandler, The Lower Tertiary Floras of Southern England IV: A Summary and Survey of Findings in Light of Recent Botanical Observations. (British Museum of Natural History, London, 1964).
67. D. H. Mai, Entwicklung der Wasser- und Sumpfplanzen-Gesellschaften Europas von der Kreide bis ins Quartär. *Flora* **176**, 449–511 (1985).
68. P. I. Dorofeev, Tertiary Floras in Western Siberia. (Akademia Nauk SSSR, Leningrad, 1963).
69. E. M. Friis, P. R. Crane, K. R. Pedersen, Early Flowers and Angiosperm Evolution. (Cambridge University Press, Cambridge, 2011).
70. M. Crawley, Angiosperm woods from British Lower Cretaceous and Palaeogene deposits. *Spec. Pap. Palaeontol*. **66**, 1–100 (2001).
71. E. M. Friis, K. R. Pedersen, P. R. Crane, Cretaceous diversification of angiosperms in the western part of the Iberian Peninsula. *Rev. Palaeobot. Palyno*. **162**, 341–361 (2010).
72. J. A. Doyle, P. K. Endress, G. R. Upchurch, Early Cretaceous monocots: A phylogenetic evaluation. *Acta Mus. Natl. Pragae, Ser. B Hist. Nat.* **64**, 59–87 (2008).
73. W. J. D. Iles, S. Y. Smith, M. A. Gandolfo et al., Monocot fossils suitable for molecular dating analyses. *Bot. J. Linn. Soc*. **178**, 346–374 (2015).
74. G. J. Jordan, M. K. Macphail, A Middle-Late Eocene inflorescence of Caryophyllaceae from Tasmania, Australia. *Am. J. Bot*. **90**, 761–768 (2003).
